# “Evolution of the mutation spectrum across a mammalian phylogeny”

**DOI:** 10.1101/2023.05.31.543114

**Authors:** Annabel C. Beichman, Jacqueline Robinson, Meixi Lin, Andrés Moreno-Estrada, Sergio Nigenda-Morales, Kelley Harris

## Abstract

Little is known about how the spectrum and etiology of germline mutagenesis might vary among mammalian species. To shed light on this mystery, we quantify variation in mutational sequence context biases using polymorphism data from thirteen species of mice, apes, bears, wolves, and cetaceans. After normalizing the mutation spectrum for reference genome accessibility and *k*-mer content, we use the Mantel test to deduce that mutation spectrum divergence is highly correlated with genetic divergence between species, whereas life history traits like reproductive age are weaker predictors of mutation spectrum divergence. Potential bioinformatic confounders are only weakly related to a small set of mutation spectrum features. We find that clocklike mutational signatures previously inferred from human cancers cannot explain the phylogenetic signal exhibited by the mammalian mutation spectrum, despite the ability of these clocklike signatures to fit each species’ 3-mer spectrum with high cosine similarity. In contrast, parental aging signatures inferred from human de novo mutation data appear to explain much of the mutation spectrum’s phylogenetic signal when fit to non-context-dependent mutation spectrum data in combination with a novel mutational signature. We posit that future models purporting to explain the etiology of mammalian mutagenesis need to capture the fact that more closely related species have more similar mutation spectra; a model that fits each marginal spectrum with high cosine similarity is not guaranteed to capture this hierarchy of mutation spectrum variation among species.

## Introduction

Germline mutations likely arise from a mixture of different processes including DNA replication errors and chemical DNA damage (Lindahl and Wood 1999; Hoeijmakers 2001). Although the relative contributions of these exogenous and endogenous processes are unknown, the action of specific mutagens can sometimes be inferred by classifying mutations into a spectrum of measurable mutation types, for example, SNPs occurring in different 3-mer contexts (Hwang and Green 2004; Alexandrov et al. 2013). Studies of somatic mutations in cancer have revealed that exogenous mutagens and DNA repair deficiencies can dramatically affect the mutation spectrum in a way that is informative about the biology of the cancer and its likely susceptibility to chemotherapies (Nik-Zainal et al. 2012). Many of the same mutational processes also affect normal tissues and provide insights into mechanisms of aging (Martincorena et al. 2015; Cagan et al. 2022).

Germline mutation spectra tend to be less variable than somatic mutation spectra, but subtle stratifications still exist (Harris 2015; Harris and Pritchard 2017; Mathieson and Reich 2017; Narasimhan et al. 2017; Moore et al. 2021; Gao et al. 2023). Some of these stratifications reflect the aging of parental gametes; for example, children born to older mothers tend to have more C>G de novo mutations (Goldmann et al. 2016; Wong et al. 2016; Jónsson et al. 2017). Measurements of germline mutation accumulation patterns are beginning to overturn long-held theories about the biology of reproduction, including the assumption that most genetic variation stems from DNA replication errors in the adult testis (Gao et al. 2019; Wu et al. 2020; Seplyarskiy and Sunyaev 2021; Hahn et al. 2023).

One source of information about germline mutagenesis is genetic variation: polymorphisms that are measured by sequencing unrelated individuals are relics of mutations that occurred generations ago in the ancestors of the sampled individuals. These mutation spectra can be complicated to interpret because of perturbations introduced by natural selection and biased gene conversion (Duret and Galtier 2009; Ratnakumar et al. 2010; Vollger et al. 2023). However, polymorphism data suggest that many species and populations have distinct mutation spectra (Moorjani et al. 2016; Harris and Pritchard 2017; Mathieson and Reich 2017; Dumont 2019; Jiang et al. 2021; Goldberg and Harris 2022; Sasani et al. 2022; Bloom et al. 2023) and these differences generally do not fit the classical profile of biased gene conversion (Harris and Pritchard 2017; Gao et al. 2023). Mutation spectrum variation is generally inferred from polymorphisms in non-conserved, non-coding genomic regions, meaning that natural selection is not likely to be the driving force behind these differences.

One pattern that has been qualitatively noted in humans and other great apes is that the mutation spectrum appears to be a phenotype with phylogenetic signal (DeWitt et al. 2021; Goldberg and Harris 2022), meaning that more distantly related lineages generally have less similar mutation spectra than more closely related lineages. This pattern is consistent with the hypothesis that the mutation spectrum is a genetically determined phenotype that evolves over time due to the emergence of new mutator alleles (Sturtevant 1937; Lynch 2010; Sung et al. 2012; Lynch et al. 2016) that each perturb different DNA repair pathways and tend to act in different sequence contexts. Mutator variants have been identified in human families (Robinson et al. 2021; Kaplanis et al. 2022) as well as certain populations of yeast, mice, and primates (Jiang et al. 2021; Sasani et al. 2022; Stendahl et al. 2023), but these variants can only explain a small proportion of the mutation spectrum variation that exists within these species. It is unclear whether the remaining variation was created by undiscovered mutators versus environmental or demographic forces, which might also create phylogenetic signal under certain circumstances (Thomas et al. 2018; Coll Macià et al. 2021; Wang et al. 2022; Wang et al. 2023).

Life history traits such as generation time have a clear impact on the germline mutation rate and spectrum (as well as the somatic mutation spectrum) (Risch et al. 1987; Sayres et al. 2011; Bromham et al. 2015; Cagan et al. 2022). Body size and longevity may also affect germline mutagenesis by incentivizing evolution of better DNA repair to avoid cancer growth (Nabholz et al. 2008; Caulin and Maley 2011; Abegglen et al. 2015; Vazquez and Lynch 2021); in rockfish, longevity appears to be correlated with the rate of CpG transition mutations (Kolora et al. 2021). A large recent study of vertebrate de novo mutations found support for the idea that generation time affects the mutation rate, though interestingly it found no support for the impact of body size (Bergeron et al. 2023). To better understand how genetics, environment, and age interact to shape the accumulation of mutations in the germline, more standardized mutation data from a variety of taxa will be needed.

In this study, we use publicly available whole-genome polymorphism data to study mutation spectrum evolution over a phylogeny that spans rodents, primates, cetaceans, and carnivorans. We generate mutation spectra from each species using a pipeline that is designed to minimize variation caused by reference genome composition, sample size, genome accessibility, and population history. Since bioinformatic batch effects are a significant obstacle to the reanalysis of data from multiple studies that were generated at different times using different technologies under different budgetary constraints, we explore the apparent dependence of the mutation spectrum on confounders, including genome assembly quality and resequencing read coverage (Taub et al. 2010; Tom et al. 2017; Leigh et al. 2018; Anderson-Trocmé et al. 2020). We then use these data to explore how the mutation spectrum might evolve as a function of biological variables like reproductive life history, testing the predictions of several key hypotheses about the origin and evolution of germline mutations.

## Results

### Standardizing the mutation spectrum for genome composition and genetic diversity

We estimated 1-mer, 3-mer, 5-mer and 7-mer mutation spectra (**Figure 1**) using polymorphisms annotated as high quality in whole-genome sequence data sampled from six primate species, two rodents, two cetaceans, and three carnivorans (**Figure 2A, Table S1**). These species span 100 million years of mammalian evolution (**Figure S1**). They also vary considerably in body size, generation time, lifespan and environment, which are important variables that have the potential to influence DNA damage, repair, and replication.

**Figure 1.**
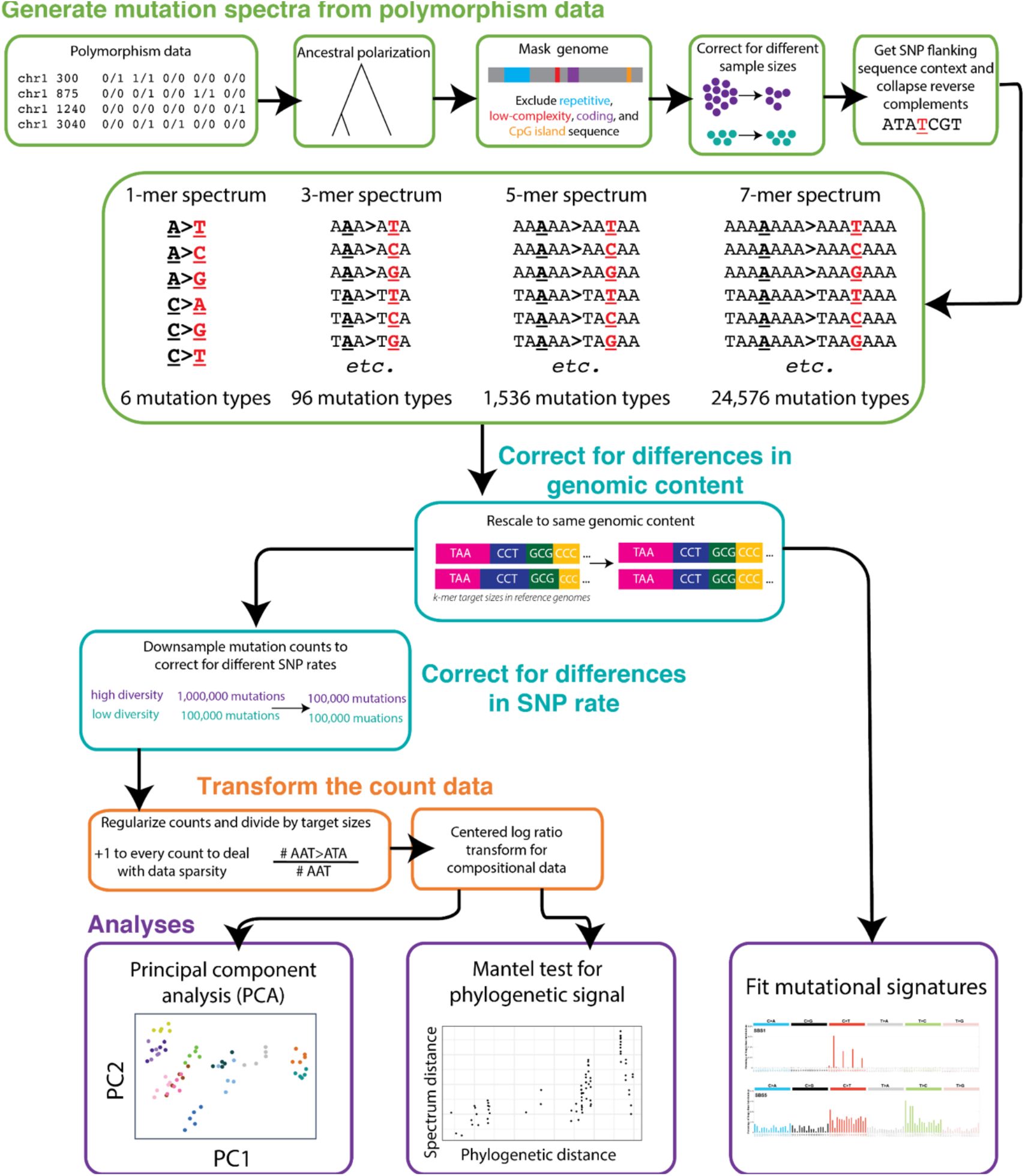
Analysis workflow. Our approach for comparing mutation spectra between species. Details in **Methods**.

**Figure 2.**
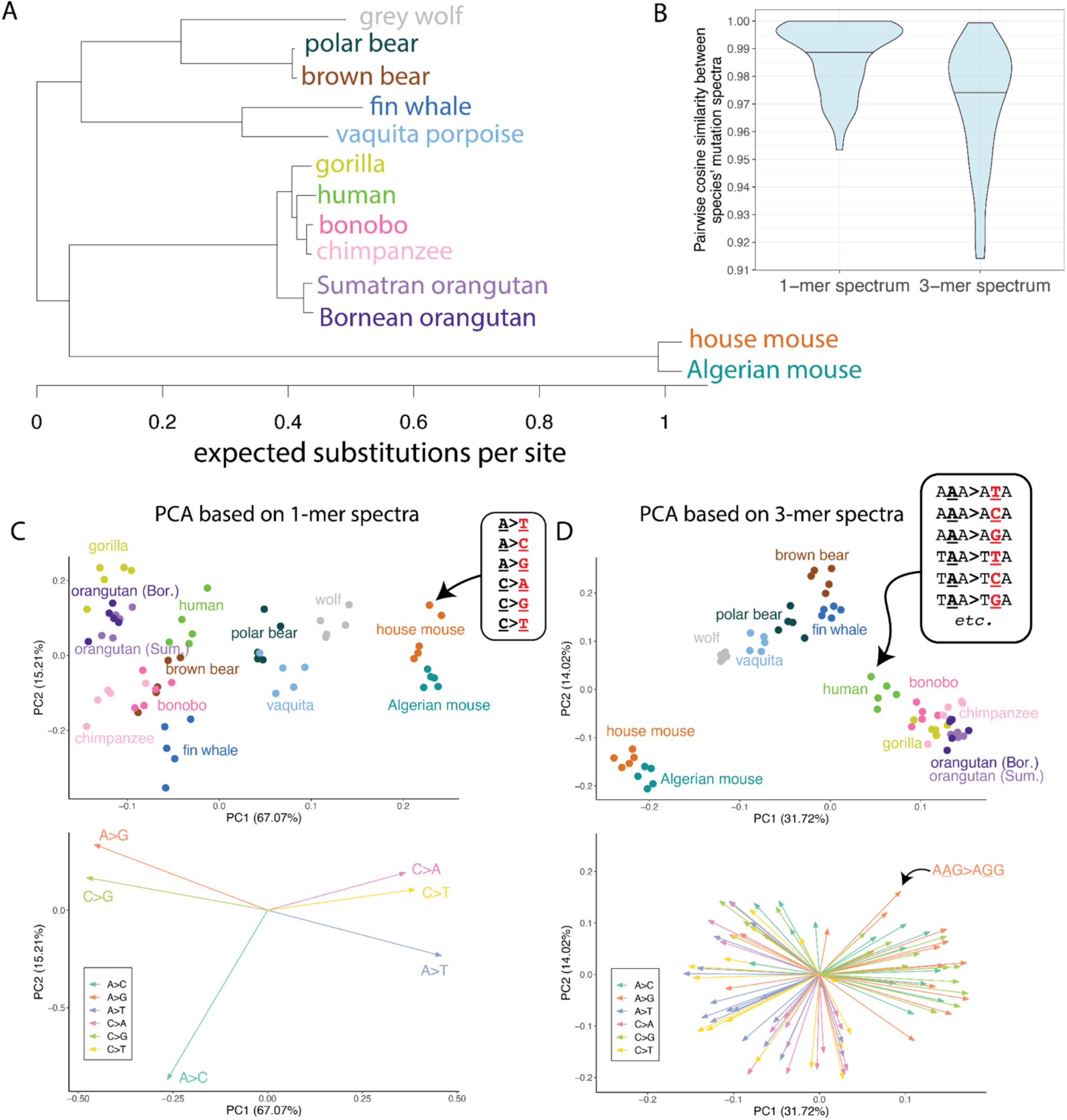
Principal components of 1-mer and 3-mer mutation spectrum variation reflect phylogenetic relationships among species. **A)** *RAxML* tree from Upham et al. (2019), restricted to species included in our study. Branch lengths represent the expected substitutions per site in Upham et al’s 31-gene sequence alignment. **B)** Distributions of cosine similarities between 1-mer and 3-mer mutation spectra for every pair of species in our dataset (**Table 1**). Horizontal lines denote the median. **C)** PCA of 1-mer mutation spectra. Each point represents a single individual’s 1-mer mutation spectrum. Points are colored and labeled according to species membership (**Table 1**). To avoid oversampling any species, five representative individuals were chosen at random from each species. SNPs were rescaled to the same genomic content across species and Poisson-downsampled to the minimum number of SNPs observed across all individuals. The resulting mutation spectra were CLR transformed as described in the **Methods**. PCA loadings are shown in the panel below, colored and labeled by mutation type (dark green: A>C; orange: A>G; purple: A>T; pink: C>A; light green: C>G; yellow: C>T). **D)** PCA of 3-mer spectra constructed as in (B). 3-mer mutation type loadings in the lower panel are colored by their central mutation types (e.g. A**<u>A</u>**G>A**<u>G</u>**G is labeled for illustration, and is colored orange as it is a type of A>G mutation). Plots including PC3 are shown in **Figure S2**. PCAs of isometric log-ratio transformed (ILR) spectra, which qualitatively resemble the centered-log-ratio (CLR) transformed spectra, are shown in **Figure S3**.

To interpret the counts of each *k*-mer-based mutation type as proxies for context-dependent mutation rates, we developed a novel standardized pipeline to normalize these counts for accessible genome composition. We first excluded genomic regions where SNP calls are likely to be unreliable (low complexity regions, repeat regions, CpG Islands) as well as regions subject to strong purifying selection (genic regions and surrounding regulatory regions) (**Figure 1**). We then transformed raw SNP counts to minimize differences between species caused by sample size, *k*-mer composition of the accessible part of the reference genome, and demographic history (**Figure 1**). Finally, we transformed the distance between the normalized mutation spectra of each pair of species via Aitchison’s centered log ratio to eliminate spurious correlations that can affect vectors of compositional data (Pearson 1897; Aitchison 1986).

### Principal component analysis of mutation spectra reveals clustering by species and higher-order clade

After filtering and normalizing all species’ mutation spectra, we explored several strategies for measuring their similarity to one another. One metric commonly used to compare mutation spectra is cosine similarity (Kucab et al. 2019; Alexandrov et al. 2020). Highly divergent mutation spectra from different cancer types typically have low cosine similarities, and a high cosine similarity between a model fit and data (e.g. higher than 0.9-0.95) is typically used as evidence that a cancer spectrum can be adequately explained as the composite of mutational signatures previously recorded in the COSMIC signature catalog (Gori and Baez-Ortega 2020; Kolora et al. 2021; Islam et al. 2022).

We observe a high degree of cosine similarity between pairs of species’ 1-mer and 3-mer spectra– roughly half of the comparisons have cosine similarity greater than 0.98 and might be considered essentially identical if cosine similarity were the only metric used to assess mutation spectrum differences (**Figure 2B**). However, since polymorphisms are much more numerous than somatic mutation counts derived from individual tumors, we hypothesized that polymorphism-based spectra exhibiting high cosine similarity might still exhibit differences that are robust and statistically significant, as previously seen using principal component analysis of human and great ape mutation spectra (Harris and Pritchard 2017; Goldberg and Harris 2022). This hypothesis is supported by a principal component analysis of our normalized 1-mer and 3-mer spectra, which reveals clustering of individuals by species and higher order clade (**Figure 2C-D**). Notably, bears, wolves, vaquitas, and fin whales cluster together as per phylogenetic expectation, despite the fact that these species were all sequenced as part of different studies with different bioinformatics protocols and different reference genome qualities. We note that the mice, which are outliers on PC1, also have the greatest genetic distance to all other clades due to a long internal branch in the phylogeny (**Figure 2A**). This long branch is likely caused by a shortening in the murine clade of the generation time (Martin and Palumbi 1993).

### Testing for the significance of phylogenetic signal

We used the Mantel test to quantify the correlation between phylogenetic distance and mutation spectrum divergence that is qualitatively seen in **Figure 2**. This involves permuting the matrix of pairwise mutation spectrum distances to construct a well-calibrated null for assessing the significance of the spectrum distance’s correlation with phylogenetic distance (Mantel 1967; Harmon and Glor 2010; Hardy and Pavoine 2012; Legendre and Legendre 2012). Phylogenetic branch lengths were calculated from a published *RAxML* tree (Upham et al. 2019) with branch lengths representing expected substitutions per site (**Figure 2A**; ultrametric timetree in **Figure S1**). Since the divergence in a trait evolving under a Brownian motion model is expected to scale with the square root of cophenetic distance (Hardy and Pavoine 2012), we tested for a significant correlation between each mutation spectrum distance and the square root of the substitution rates that Upham et al. (2019) estimated using a multispecies sequence alignment.

Using a Mantel Test with 9,999,999 permutations, we found that both the 6-dimensional 1-mer mutation spectrum and the 96-dimensional 3-mer spectrum exhibited significant phylogenetic signal (*r* = 0.68, *p* < 8e-6 and *r* = 0.82, *p* < 3e-7; respectively) (**Figure 3A, Figure S4** for labeled comparisons). This phylogenetic signal appears robust to many analysis variations, including using an ultrametric phylogenetic tree, substituting cosine distance or isometric log-ratio (ILR) distance for the CLR distance (Egozcue et al. 2003), and ‘folding’ the mutation spectrum to remove any effects of ancestral allele misidentification (**Figures S5-S8**). Due to their high mutation rates, CpG>TpG mutations are sometimes censored from polymorphism-based spectra or separated out as their own mutation class (Gao et al. 2023; Wang et al. 2023), and we find that neither of these choices appreciably reduces the phylogenetic signal (**Figure S9**).

**Figure 3.**
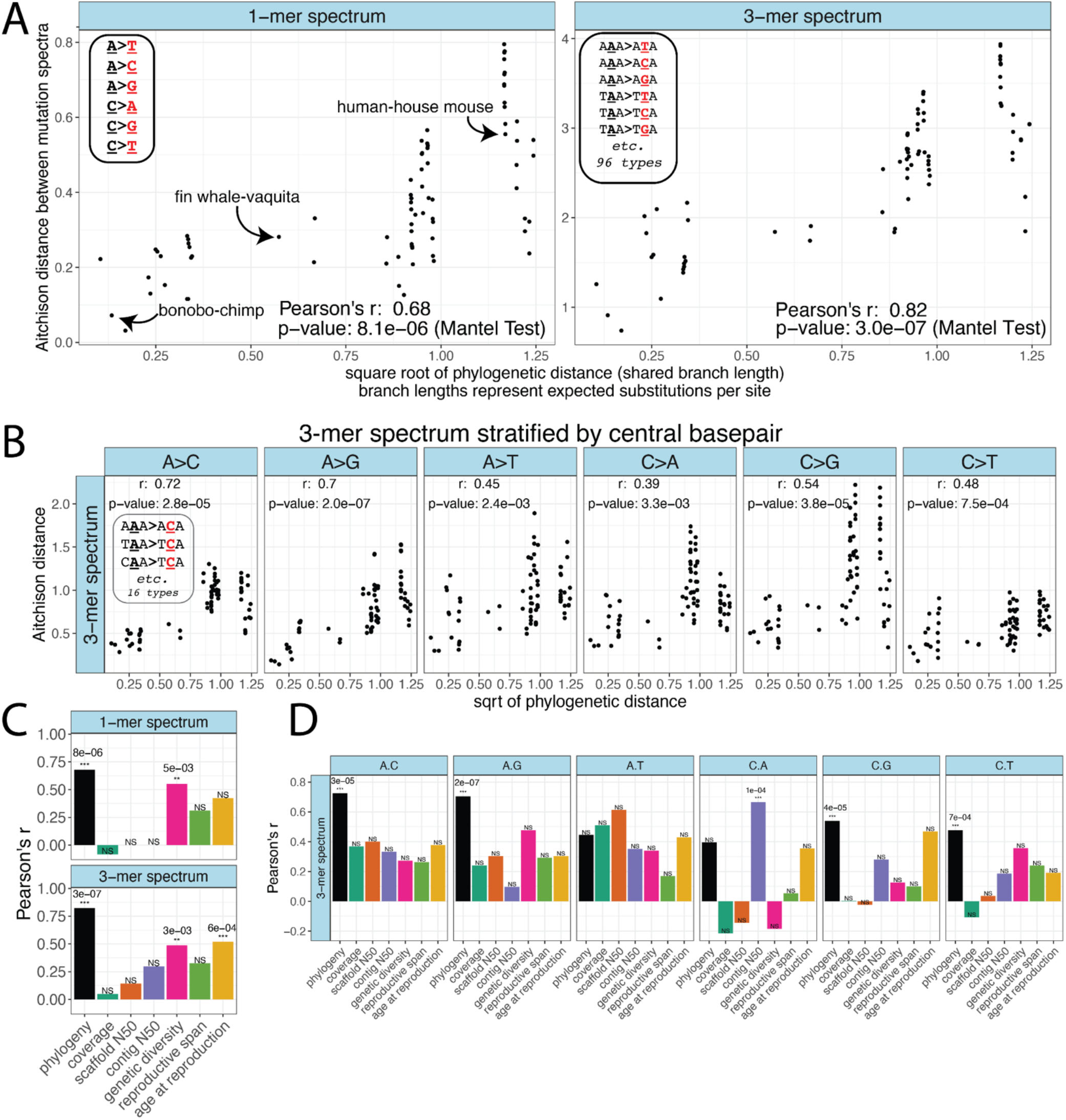
Mutation spectrum distance between species shows a phylogenetic signal. **A)** Correlation between pairwise mutation spectrum distance and the square root of phylogenetic distance between pairs of species. Distances reflect CLR-transformed 1-mer mutation spectra and 3-mer spectra for each species such that each point represents a single between-species comparison (e.g. human vs. house mouse). Note that unlike in Figure 2, mutation spectra are calculated across all five sampled individuals per species, rather than per individual. *p*-values are based on the Mantel test with 9,999,999 permutations. A version of these plots with each point labeled is shown in **Figure S4**. **B)** Correlation between mutation spectrum distance and phylogenetic distance exists within the 3-mer sub-spectrum of each 1-mer mutation type (e.g. distances in the “A>C” column distances are calculated based on the 16 A>C 3-mers only). **C)** Values of Pearson’s *r* for the correlations across species between technical and biological variables and mutation spectrum distance, calculated using the Mantel test with 99,999 permutations. The black “phylogeny” column represents the phylogenetic signal *p*-values reported in (A). “NS” (non-significant) denotes *p*-values that fell above a significance threshold Bonferroni-corrected for 7 tests (α = 0.007 = 0.05/7). *p*-values below this threshold are denoted using “**” if < 0.007 but > 0.001, and “***” if < 0.001, and the specific *p*-values are written above the corresponding column. **D)** Values of Pearson’s *r* for the correlation between different variables and the 3-mer spectrum when it is stratified by central 1-mer type (as in (B)). *p*-values from Mantel tests with 99,999 permutations. The black “phylogeny” columns represent the *r*-values in (B). NS (non-significant) designates *p*-values that fall above a significance threshold Bonferroni-corrected for 42 tests: 7 confounders for each of 6 mutation types (α = 0.05/(6*7) = 0.001). *p*-values below this threshold are denoted with “***” and the specific *p*-value is written above the corresponding column. Note that when carrying out the Mantel test for these potential confounders we used fewer permutations than in (A) since no confounding variable reached the minimum *p*-value for 99,999 permutations (1e-5) that would have required additional permutations. Phylogenetically-aware Mantel results for panels C and D are in **Figure S13**.

We additionally tested for the presence of phylogenetic signal using the *K_mult_* statistic (Adams 2014; Adams and Collyer 2018), a multivariate version of Blomberg’s *Κ* (Blomberg et al. 2003). We found significant phylogenetic signal at both the 1-mer and 3-mer level (1-mer: *K_mult_* = 0.31, *p* < 0.001; 3-mer: *K_mult_* = 0.26, *p <* 0.001; 999 permutations) (**Figure S10**). The values of *K_mult_* were less than 1 (1 being the expected value under Brownian motion). This deviation of the Brownian motion expectation is typical of most assayed multivariate traits and could be due to noise or natural selection. Another possibility for *K_mult_* being less than 1 is that the number of independently varying mutation spectrum components may be much smaller than the dimensionality of the full *k*-mer mutation spectrum (Adams and Collyer 2019), as seen in mutational signature decompositions where a relatively small number of mutational processes explain mutation spectrum variability across samples.

### Mutation spectrum divergence is not consistent with the GC-biased signature of gene conversion

One potential contributor to phylogenetic divergence between mutation spectra is GC-biased gene conversion (gBGC), a process that drives mutations from A/T to G/C to rise in frequency over time while driving mutations from G/C to A/T to decline in frequency (Duret and Galtier 2009). Species with the highest effective population sizes are expected to experience the strongest gBGC, leading to a well-understood distortion of the mutation spectrum. However, gBGC is not expected to affect C>G or A>T mutations, and it is not known to affect the *k*-mer sequence composition within each 1-mer mutation class.

When we performed Mantel tests on the spectrum of 3-mer mutation types partitioned into categories that experience different modalities of BGC-induced selection (neutral A>T and C>G; negatively selected C>A and C>T, and positively selected A>C, A>G), we found highly significant phylogenetic signal within each category, notably including the BGC-neutral (A>T + C>G) category (**Figure S11**; *r* = 0.79, *p* < 5e-6). We then partitioned 3-mers by 1-mer mutation class and still found significant phylogenetic signal, with Mantel test *p*-values ranging from 3.3e-3 (for C>A 3-mers) to 2e-7 (for A>G 3-mers; all less than the Bonferroni-corrected threshold α = 0.05/6 = 0.0083 appropriate for a set of 6 tests) (**Figure 3B**). This implies that GC-biased gene conversion cannot be the primary force that causes the mutation spectrum to have phylogenetic signal.

### Differences in bioinformatic data quality are unlikely to explain the observed mutation spectrum differences among species

A potential caveat to the above results is that if data quality and bioinformatic processing tend to be more similar among more closely related species, this could create the false appearance of a correlation between mutation spectrum similarity and phylogenetic relatedness. There are several reassuring indications that our dataset does not have this property: for example, the best quality chromosomal genome assemblies by several metrics are human and vaquita, two species which are not closely related, and the oldest-generated datasets (the great apes (published in 2013), mice (2016), and bears (2012-2018)) are likewise dispersed across the phylogeny.

To formally test whether technical artifacts are phylogenetically distributed across our dataset, we used the Mantel test to measure the correlation of phylogenetic distance with three technical variables: average sequencing coverage, reference genome scaffold N50, and reference genome contig N50. Differences between species’ scaffold N50 and sequence coverage showed no significant correlation with phylogenetic distance (**Figure S12A, S12C**). Differences in contig N50 (the relative contiguity of contigs prior to scaffolding) showed a moderate phylogenetic signal (Pearson’s *r* = 0.42, *p*-value < 0.007, Mantel test with 99,999 permutations) (**Figure S12B**), but this correlation coefficient and *p*-value are more modest than those of the correlation between mutation spectrum distance and phylogenetic distance. While technical confounders may explain a small portion of the phylogenetic signal in the dataset, they do not appear sufficient to explain the results shown in **Figure 3A**.

To more directly assess what role (if any) these technical confounders may play in causing differences between our species’ mutation spectra, we directly tested each technical confounder for correlation with mutation spectrum distance (Mantel test with 99,999 permutations) **(Figure 3C)**. Our aim here was to test whether any of these measurements explains mutation spectrum divergence *better* than the phylogeny does. We found that differences in contig N50, scaffold N50, and sequence coverage between species are *not* significantly correlated with 1-mer and 3-mer mutation spectrum distance after correction for multiple testing (**Figure 3C**). Our result indicates that these confounders cannot be responsible for the differences between mutation spectra we observe, though any correlations between the mutation spectrum and these technical measurements could result from a shared phylogenetic signal. We obtain qualitatively similar results using a phylogenetically-aware Mantel test that asks whether each technical covariate explains additional mutation spectrum divergence on top of what is explained by the phylogeny (**Figure S13**).

When we look at the 3-mer sub-spectra of individual 1-mer mutations (particularly A>T and C>A 3-mers), it is less consistently clear that phylogeny explains sub-spectrum variation better than technical factors (**Figure 3D**). After stratifying the spectra by central basepair, we observed that contig N50 is more significantly correlated with differences in the C>A 3-mer spectrum than phylogenetic distance is (*r* = 0.67, *p* < 1.1e-4 for the correlation between C>A 3-mers and contig N50 compared to *r* = 0.39, *p* < 3.3e-3 for C>A 3-mers and phylogenetic distance; **Figure 3D, Figure S13** for phylogenetically-aware Mantel results). Scaffold N50 is more correlated with A>T 3-mer spectrum distances than phylogenetic distance is (*r* = 0.6, *p* < 2.1e-3 for A>T 3-mers and scaffold N50 compared to *r* = 0.45, *p* < 2.4e-3 for A>T 3-mers and phylogenetic distance), though neither correlation passes the significance threshold after correction for multiple tests (α = 1e-3). However, after correcting for multiple testing, the phylogeny is the only significant covariate with 3-mer mutation spectrum divergence within each of the remaining four mutation types (A>C, A>G, C>G, and C>T).

### Differences in reproductive age and effective population size do not appear to drive the mutation spectrum’s phylogenetic signal

Recent studies of human and animal de novo mutagenesis have found that the mutation rate and spectrum depend on age at reproduction (Goldmann et al. 2016; Wong et al. 2016; Jónsson et al. 2017; Thomas et al. 2018; Bergeron et al. 2023). Motivated by this, we performed additional tests to calculate the correlation of mutation spectrum divergence with maximum reproductive lifespan and age at first reproduction (Jones et al. 2009; Pacifici et al. 2013). We also measured the correlation between mutation spectrum distance and the genetic diversity metric Watterson’s *θ*, since diversity is strongly correlated with the strength of GC-biased gene conversion. We expected these biological confounders to be at least partially phylogenetically distributed across our dataset, and so first tested each variable for a significant phylogenetic signal (**Figure S14**). Age at first reproduction exhibited no significant phylogenetic signal in this dataset after correction for multiple testing (Pearson’s *r* = 0.32, *p* < 0.024, Bonferroni α = 0.008 as appropriate for six tests for phylogenetic signal across the six confounders, **Figure 14B**), but reproductive lifespan and Watterson’s *θ* each exhibited moderate phylogenetic signal (Pearson’s *r* = 0.40, *p* < 0.005; *r* = 0.6, *p* < 0.0013; respectively) (**Figure S14A, S14C**).

To determine what impact these biological variables may have in shaping mutation spectrum distances across our species, we then directly correlated each of them with pairwise mutation spectrum distances (**Figure 3C**). Age at first reproduction is significantly correlated with differences in the 3-mer spectrum after correction for multiple testing (age at first reproduction: *r* = 0.52, *p* < 0.00057; Bonferroni α = 0.007 as appropriate for a set of seven tests for correlation with different biological and technical variables) (**Figure 3C**), but correlation of phylogenetic distance with the 3-mer mutation spectrum is stronger. Reproductive lifespan is marginally significantly correlated with the 3-mer spectrum after correction for multiple tests (r = 0.33, p < 0.007, Bonferroni α = 0.007) (**Figure 3C**). Age at first reproduction is also marginally significantly correlated with the 1-mer spectrum (*r* = 0.42, *p* < 0.0076, Bonferroni α = 0.007) (**Figure 3C**). Interestingly, the significant relationships of generation time and reproductive lifespan with the mutation spectrum persist after carrying out a phylogenetically-aware version of the Mantel test (**Figure S13**), meaning that it is unlikely that these correlations are driven by shared phylogenetic signal and may instead reflect a role of generation time in shaping mutation spectrum patterns between species.

We found Watterson’s *θ*, an indicator of recent effective population size, to be correlated with the 1-mer and 3-mer mutation spectra at significance levels *p* < 0.005 (*r* = 0.55) and *p* < 0.003 (*r* = 0.48) (**Figure 3C**). These are weaker than the correlation between mutation spectrum distance and phylogenetic distance (**Figure 3C**), further indicating that differences in biased gene conversion strength driven by effective population size are likely not strong enough to fully explain the phylogenetic signal we observe in the 1- and 3-mer mutation spectra. When we recalculated the correlation between mutation spectrum distance and genetic diversity accounting for shared phylogenetic signal affecting both traits, both the 1-mer and 3-mer correlations remained significant (*p*-values < 0.0008 and 0.0004, respectively) (**Figure S13**). Genetic diversity is generally uncorrelated with 3-mer composition after partitioning by 1-mer type (**Figure 3D, Figure S13**). This is consistent with our expectation that genetic diversity affects the mutation spectrum primarily through GC-biased gene conversion, and that biased gene conversion affects mutation spectrum composition at the 1-mer level, but not the 3-mer level.

### Evolution of mutation rate dependence on extended sequence context

Although 1-mer and 3-mer mutation spectra are commonly used to study mutational patterns in datasets of modest size, a few studies of human genetic variation have found that mutation rate can depend on extended sequence context over *k*-mers of size 7 or greater (Aggarwala and Voight 2016; Carlson et al. 2018; Liu and Samee 2021; Adams et al. 2023). In theory, more mutational categories could yield greater power to resolve distinct mutagenic processes, but this power can only be realized given sufficient data to fill out rare mutational categories. A study of variation in 5-mer and 7-mer spectra of human populations yielded mixed results, finding some indications that 3-mer sequence context varied more among populations than extended sequence context did (Aikens et al. 2019). A few differences in 5-mer and 7-mer context dependence were observed among human populations, but it is unclear whether these results are robust to quality issues later identified in the low coverage 1000 Genomes data (Anderson-Trocmé et al. 2020).

The first two principal components of higher-dimensional mutation spectrum variation explain less variance than we observed in PCAs of 1-mer and 3-mer spectra, but clusters within species and higher-order clades appear more visually distinct, with lower cosine similarities between the most distant pairs of species (**Figure S15-S16**). Higher-dimensional mutation spectra have significant but somewhat weaker phylogenetic signal compared to the 3-mer mutation spectrum (**Figures 5A**, see additional 5-mer and 7-mer spectrum and sub-spectrum phylogenetic signal details in **Note S1** and **Figures S17-S22)**.

If phylogenetic signal exists at the 3-mer level, a Mantel test on the 5-mer mutation spectrum is likely to show phylogenetic signal even if the effects of extended sequence context on mutation rate are invariant among species. We therefore devised a permutation test to investigate whether each 5-mer’s species-specific mutation rate is conditionally independent of the phylogeny after controlling for variation of the 3-mer mutation rate among species. The test involves randomizing the distribution of 5-mer counts within 3-mer equivalence classes to generate 5,000 control spectra per species where the dependence of mutation rate on non-adjacent nucleotides is eliminated (**Methods**). We then compared the empirical correlation of mutation spectrum distance and phylogenetic distance to that of the randomized 5-mer control spectra (**Figure 4B**; see **Figure S22** for an example comparing the phylogenetic signal of a single randomized 5-mer replicate to that of the empirical 5-mer spectrum).

**Figure 4.**
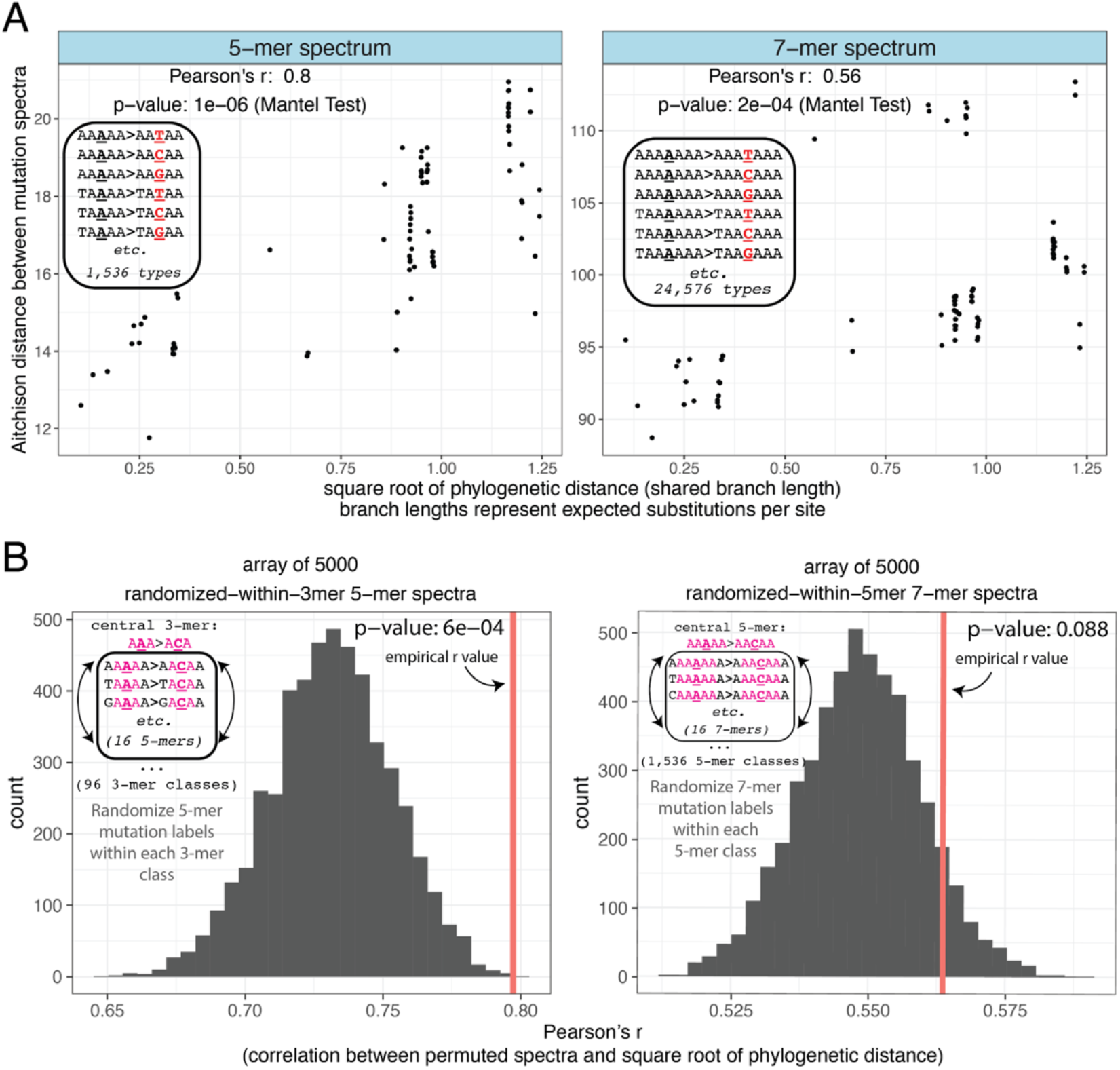
The 5-mer mutation spectrum exhibits additional phylogenetic signal beyond what is inherited from the 3-mer mutation spectrum. **A)** 5-mer and 7-mer spectra show significant phylogenetic signal. Distances were calculated based on CLR-transformed mutation spectra such that each point represents a single between-species comparison (e.g. human-house mouse). *p*-values were calculated using the Mantel Test with 9,999,999 permutations. A version of these plots with each point labeled is shown in **Figure S17**. **B)** The empirical 5-mer mutation spectrum is significantly more correlated with the phylogeny than a distribution of 5,000 control spectra generated by randomizing 5-mer mutation counts based on genomic target size within each 3-mer equivalence class (*p* < 6e-4, left panel). In contrast, we observe no significant difference in phylogenetic signal between the empirical 7-mer spectrum and 7-mer spectra that are randomized within 5-mer equivalence classes (*p* > 0.088).

**Figure 5.**
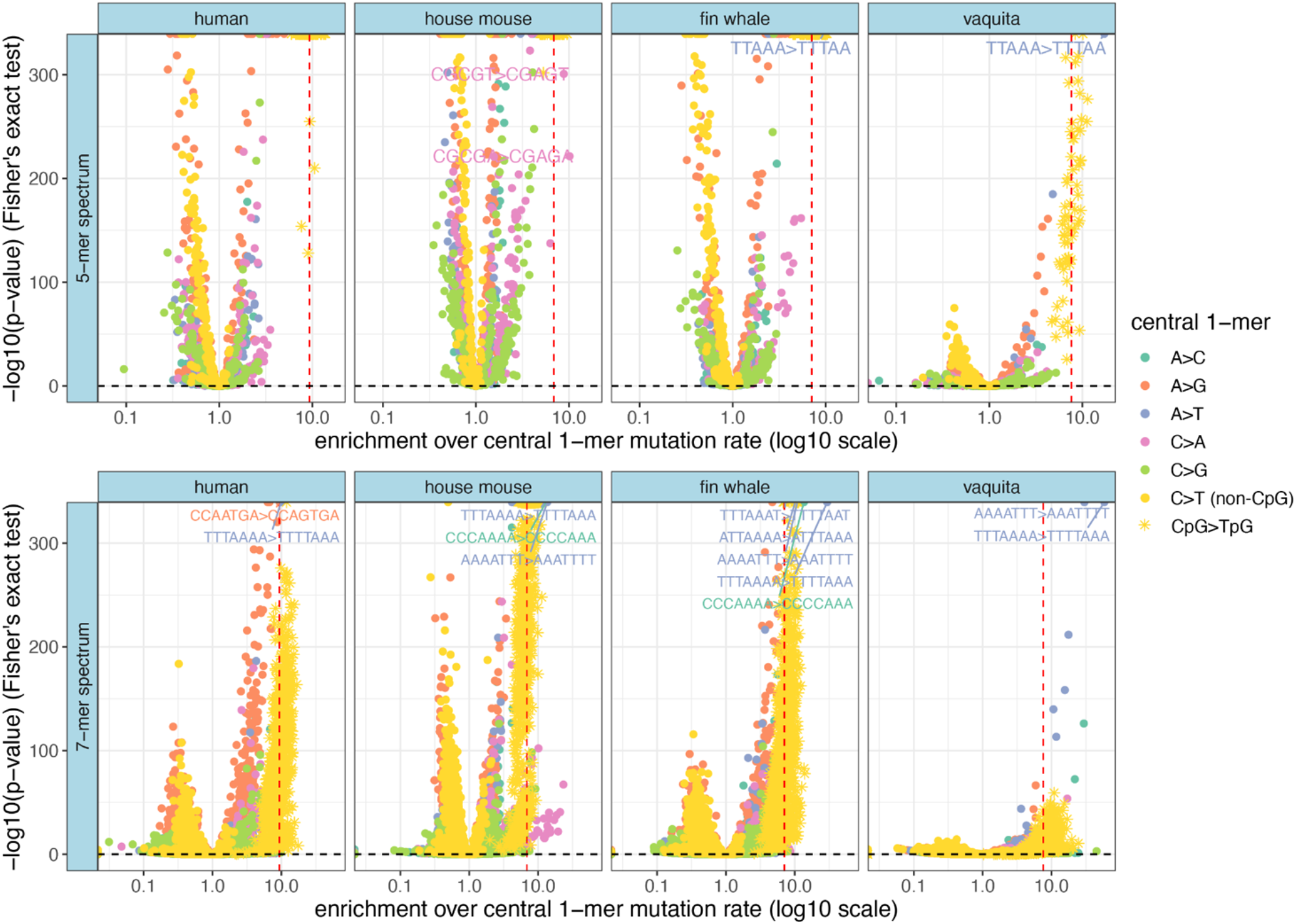
A small subset of 5-mers and 7-mers are hypermutable above the CpG>TpG level. A comparison of the mutabilities of each 5-mer or 7-mer mutation type relative to the background mutation rate of its central 1-mer (e.g. AT**<u>C</u>**CA>AT**<u>T</u>**CA rate divided by C>T rate). The *x*-axis represents the ratio of a particular *k*-mer’s rate (counts divided by target size) over the rate of its central 1-mer (counts divided by target size). The *y*-axis is the -log10(*p*-value) from a two-sided Fisher’s exact test. The horizontal black dashed line represents the Bonferroni-corrected statistical significance threshold, and the red vertical dashed line is the species-specific mutability of CpG>TpG dimers relative to the background C>T rate. *k*-mers are colored by central mutation type. Counts are not rescaled to human targets or downsampled since cross-species comparisons are not occurring. Only four species are shown: human as a baseline for comparison, house mouse which has a large number of outlying *k*-mers compared to other species, and the two whale species (fin whale and vaquita) which show extraordinary enrichment of a particular *k*-mer (TTT**<u>A</u>**AAA>TTT**<u>T</u>**AAA enriched >30-fold above the A>T rate). A small subset of the ∼100 outlying non-CpG>TpG 7-mers are labeled. **Tables S3-S4** have lists of significantly enriched 5-mers and 7-mers that exceed the CpG>TpG enrichment level. Note that enrichment *p*-values depend on the number of observed SNPs as well as fold enrichment, as evident from the trend toward lower *p*-values in the low-diversity vaquita. See **Figures S27-S29** for other species’ enrichment profiles.

We find that the empirical 5-mer spectrum distances are significantly more correlated with phylogeny compared to the 5-mer data that was randomized within each 3-mer class (p < 6e-4, 5,000 permutations; **Figure 4B, left panel)**. A similar analysis indicates that the 7-mer spectrum does *not* contain more phylogenetic signal than the dataset that is randomized to remove information beyond the 5-mer level, at least at our limited sample sizes (p < 0.09, 5,000 permutations; **Figure 4B, right panel**). After removing the lowest-diversity species in order to sample more SNPs, we see an increased separation between the empirical and permuted 5-mer spectra (*p* < 2e-4) but a decrease in the suggestive difference between the permuted and empirical 7-mer spectra (*p* > 0.75) (**Figure S23**).

As seen for 1-mer and 3-mer spectra, the phylogeny explains 5-mer and 7-mer mutation spectrum divergence consistently better than our list of technical confounders (such as reference genome contiguity) and biological confounders (such as age at first reproduction) (**Figure S24, Note S1**). However, technical confounders appear to play an important role in shaping various 5-mer and 7-mer sub-spectra, indicating that these extended context dependencies should be interpreted with caution in modestly sized datasets such as this one (**Note S1, Figures S25-S26**).

### Variation in motif hypermutability and hypomutability across species

A few sequence motifs, such as CpGs, are extremely hypermutable, with mutation rates nearly an order of magnitude above baseline. To measure how the frequency and magnitude of extreme hypermutability varies among species, we used a two-sided Fisher’s exact test to systematically compare *k*-mer-based mutation rates to the average mutation rate of the nested 1-mer (**Figure 5**). Unsurprisingly, many *k*-mer mutation rates are significantly different from the nested 1-mer rate after Bonferroni correction, but the four N**<u>C</u>**G>N**<u>T</u>**G 3-mers are consistently the most hypermutable 3-mer types across species: enrichment of CpG>TpG above the background C>T rate ranges from 6.6x (orangutan) to 9.3x (human) across the species surveyed (**Figure S27**).

As previously seen in humans, certain non-CpG 5-mer and 7-mer motifs actually have mutation rates that are more significantly elevated than those of CpG-containing motifs. We observed five distinct 5-mer mutation types that fall into this category (**Figure 5**, **Table S3**, **Figure S28**): in both whale species (fin whale and vaquita) and the polar bear, TT**<u>A</u>**AA>TT**<u>T</u>**AA is enriched ∼9-17x over the A>T rate (p-values < 2e-308). The rest of the non-CpG-transition 5-mer hypermutability is observed in mice and consists of C>A mutations in CpG rich motifs: in *Mus musculus* and *Mus spretus*, CG**<u>C</u>**GT>CG**<u>A</u>**GT (8-10x, *p*-values < 1e-200) and CG**<u>C</u>**GA>CG**<u>A</u>**GA (8-15x, *p*-values < 1e-148) were significantly enriched above the C>A rate. In *Mus spretus*, CG**<u>C</u>**AA>CG**<u>A</u>**AA (7x, *p*-value < 2e-308) and CG**<u>C</u>**GG>CG**<u>A</u>**GG (8x, p-value < 1e-190) were additionally enriched. Note that mice have the highest genetic diversity of any species in our dataset, which should allow for detection of hypermutable motifs with greater precision and recall than can be achieved with data from less diverse species.

At the 7-mer level, we observe over 100 distinct non-CpG>TpG mutation types that exceed the CpG>TpG fold-enrichment threshold in at least one species (**Figure 5, Table S4, Figure S29)**. As seen for 5-mers, mice have the largest number of hypermutable 7-mers, with *Mus spretus* having over 60 types with greater enrichment than CpG>TpG sites, the majority of which are C>A 7-mers. *Mus musculus* had over 40 enriched types (majority C>A).

In humans, Carlson, et al. (2018) previously reported that the only 7-mer more hypermutable than CpG-containing 7-mers was TTT**<u>A</u>**AAA>TTT**<u>T</u>**AAA. Aikens, et al. (2019) also noted that this motif appears to be slightly more hypermutable in Africans compared to Europeans. We find that TTT**<u>A</u>**AAA>TTT**<u>T</u>**AAA is one of the most hypermutable 7-mer types in every species in our study, with enrichments ranging from 9x-59x above species-specific A>T rates (**Figure 5B**, **Figure S29**). Its hypermutability is most extreme (30-59x above background) in the fin whale and vaquita sister lineages. We note that the fin whale and vaquita datasets were processed by different researchers and mapped to reference genomes of very different assembly quality (a highly fragmented assembly was used for the fin whale study, while a highly contiguous chromosomal-level assembly was available for the vaquita). Despite this discrepancy, fin whale and vaquita have similar hyper-enrichments of TTT**<u>A</u>**AAA>TTT**<u>T</u>**AAA, as well as shared enrichments of other repetitive 7-mers.

### Clocklike COSMIC signatures are not phylogenetically distributed

The underlying mutational processes that generate mutation spectrum patterns can be described as “mutational signatures” of known or unknown etiology. Mutational signatures are frequently used in the cancer literature to link particular environmental exposures or DNA proofreading defects to observed 3-mer somatic mutation spectrum patterns (Alexandrov et al. 2013), and the resulting signatures are maintained in the COSMIC cancer database (Tate et al. 2019). Only two COSMIC single base substitution (SBS) signatures are consistently inferred to contribute to germline mutagenesis: SBS1 and SBS5 (**Figure S30**) (Alexandrov et al. 2015; Rahbari et al. 2016; Hamidi et al. 2021; Moore et al. 2021). SBS1 has a known etiology: the deamination of methylated cytosine, resulting in C>T mutations in 3-mers that contain a central CpG sequence. SBS5 has an unknown etiology but is thought to represent a background endogenous mutational process, given its ubiquity and clock-like accumulation pattern.

To determine whether the combination of these two signatures could explain the variation in our mammalian 3-mer spectrum data, we used the *R* package *sigfit* (Gori and Baez-Ortega 2020), to model our empirical mutation spectra as linear combinations of “exposures” to SBS1 and SBS5.We found that *sigfit* inferred highly similar levels of exposure to SBS1 and SBS5 in each species in our dataset (**Figure 6A**, left panel). The corresponding mutation spectrum reconstructions fit the empirical data with high cosine similarity (0.95-0.99) (**Figure 6B**), albeit with biased residuals (across species, the model consistently underestimates the fractions of A>Gs and C>Ts while overestimating the abundance of C>As and C>Gs) (**Figure 6C**).

**Figure 6.**
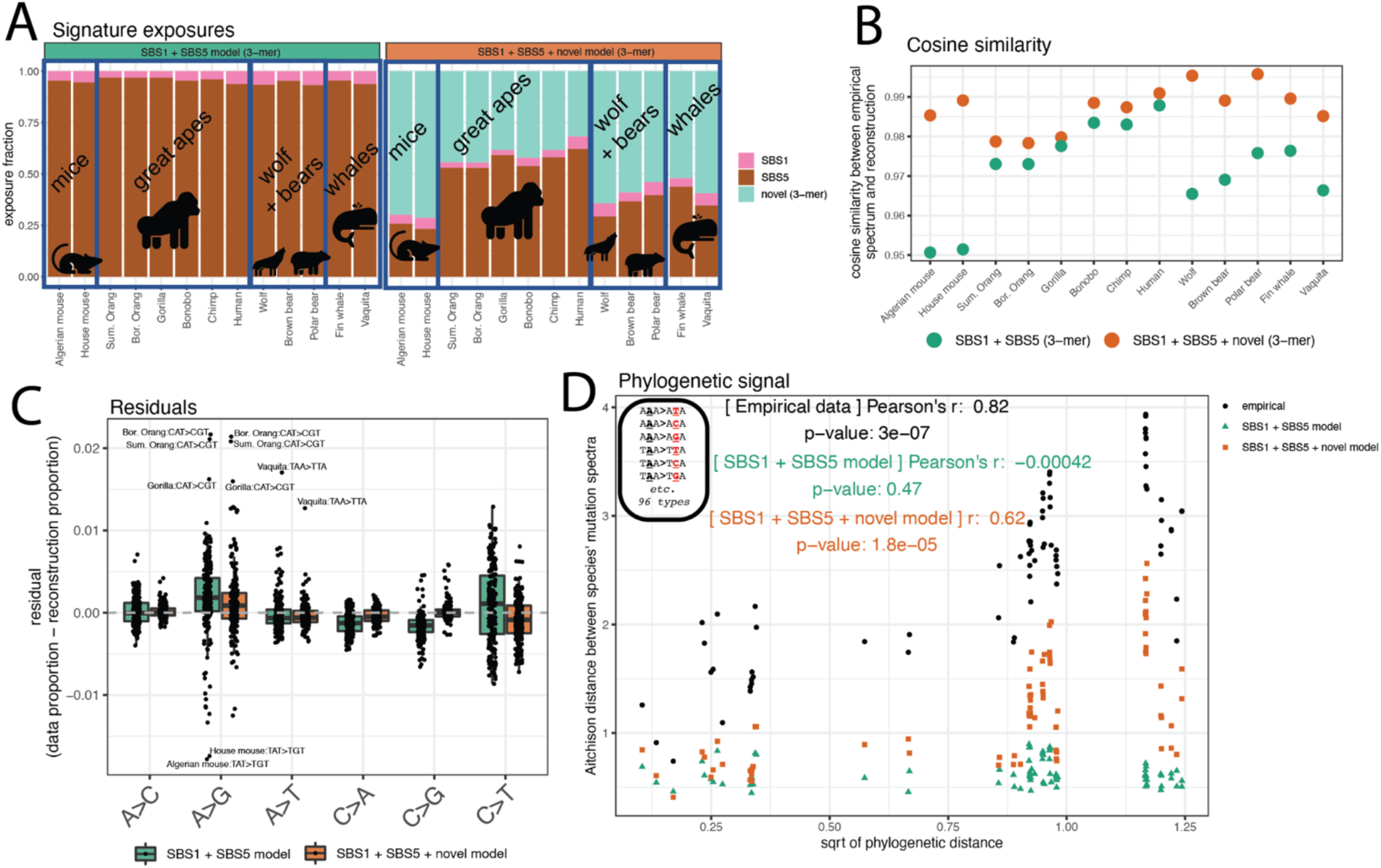
Clocklike COSMIC cancer signatures fail to reconstruct 3-mer spectra. **A)** Left panel: exposures to COSMIC cancer signatures SBS1 (pink) and SBS5 (brown), derived from human somatic data. Right panel: exposures to a model containing SBS1, SBS5, and an additional third novel signature (teal) extracted from the data. Broad phylogenetic clades are outlined and labeled. **B)** Cosine similarity between the empirical spectrum and the spectrum reconstructed using either only SBS1 + SBS5 or the SBS1 + SBS5 + novel signature. Higher cosine similarity indicates a better reconstruction of the data. **C)** Residuals (observed proportion - reconstructed proportion) for each spectrum. Boxplots are made up of the 96 3-mer mutation types for all species, separated by their central 1-mer type, with the most extreme outliers labeled. Negative residuals indicate the model reconstruction overestimating a proportion of a mutation type, positive residuals indicate an underestimate. The simple SBS1 + SBS5 model is indicated in green, the SBS1 + SBS5 + novel signature model in orange. **C)** Result of Mantel test on the correlation between Aitchison distance between species reconstructed 3-mer spectra under each model (green triangles and orange squares) and the square root of phylogenetic distance, overlaid on results based on empirical spectra (black circles). *p*-values based on 9,999,999 permutations.

Despite the high cosine similarity between the empirical mutation spectra and SBS1+SBS5 reconstructions, we find that the reconstructed mutation spectra have no significant phylogenetic signal (*r* = 0, *p* > 0.4, Mantel test with 9,999,999 permutations; **Figure 6D**). Given the strength of the phylogenetic signal in the empirical data, our results indicate that an important source of clade-specific mutation spectrum variation is missing from the SBS1+SBS5 model.

To investigate what the SBS1+SBS5 model is failing to capture in the data, we used *sigfit* to infer exposures to SBS1 and SBS5 jointly with an additional novel signature (signatures shown in **Figure S30**). Although the novel signature inferred by *sigfit* is still fairly similar to SBS5 (cosine similarity 0.94), introducing this signature increased the phylogenetic signal predicted by the model. Exposure to the novel signature is inferred to be highest in mice and lowest in great apes (**Figure 6A**, right panel), perhaps reflecting the fact that SBS5 was inferred from human data and is less tailored to the mutational processes of more distantly related species. This 4-signature model still fails to reconstruct the full phylogenetic signal of the empirical 3-mer data (**Figure 6D**), indicating that greater complexity is needed to model the cladistic mutation spectrum patterns we observe between mammals’ 3-mer spectra.

### Reproductive aging signatures inferred from human data capture 1-mer mutation spectrum differences among mammalian species

After seeing that SBS1 and SBS5 cannot explain the phylogenetic signal of the 3-mer mutation spectrum, we turned our attention to a second model of mutation spectrum etiology that attempts to explain differences at the less complex 1-mer mutation spectrum level. A previous human de novo mutation study trained a Poisson regression model to capture the dependence of the mutation spectrum on paternal and maternal age (due to data sparsity, 3-mer mutation spectrum effects were not inferred) (Jónsson et al. 2017), and several studies have argued that this reproductive aging model may largely explain 1-mer mutation spectrum differences observed among populations and species (Coll Macià et al. 2021; Wang et al. 2022; Wang et al. 2023). However, others have questioned the ability of a parental age model to explain the full range of human mutation spectrum variation (Gao et al. 2023; Ragsdale and Thornton 2023).

To test whether a reproductive aging model derived from human data is able to explain variation we observe at the 1-mer level, we attempted to reconstruct our species’ 1-mer mutation spectra using a linear combination of exposures to three parental aging signatures derived from mutation patterns observed in human families by Jónsson et al. (2017). Since Jónsson et al. modeled parental mutation contributions to the 1-mer mutation spectrum using a regression, we transformed this regression model into a mutational signature model by translating the slopes into maternal and paternal age signatures and combining the maternal and paternal regression intercepts into a “young parent” signature whose exposure is expected to be highest in the children of young parents (signatures shown in **Figure S31**). As in Wang et al. (2023) and Gao et al. (2023), we excluded CpG>TpG mutations from the signatures and data to avoid confounding by potentially elevated levels of homoplasy and mismapping affecting CpG polymorphisms.

We used *sigfit* to infer exposures to these reproductive aging signatures in our 1-mer mammalian mutation spectrum data (**Figure 7A**, left panel). The reproductive aging model fits all species’ 1-mer mutation spectra with high cosine similarities (>0.99), but also with biased residuals (across species, the model predicts too many C>Gs and too few A>Gs (**Figure 7B-C**)). This bias may be due in part to the fact that the model was trained on de novo mutation data and then fit to polymorphism data. The residuals do not conform to the expected action of biased gene conversion, as C>G is a GC-conservative mutation type.

**Figure 7.**
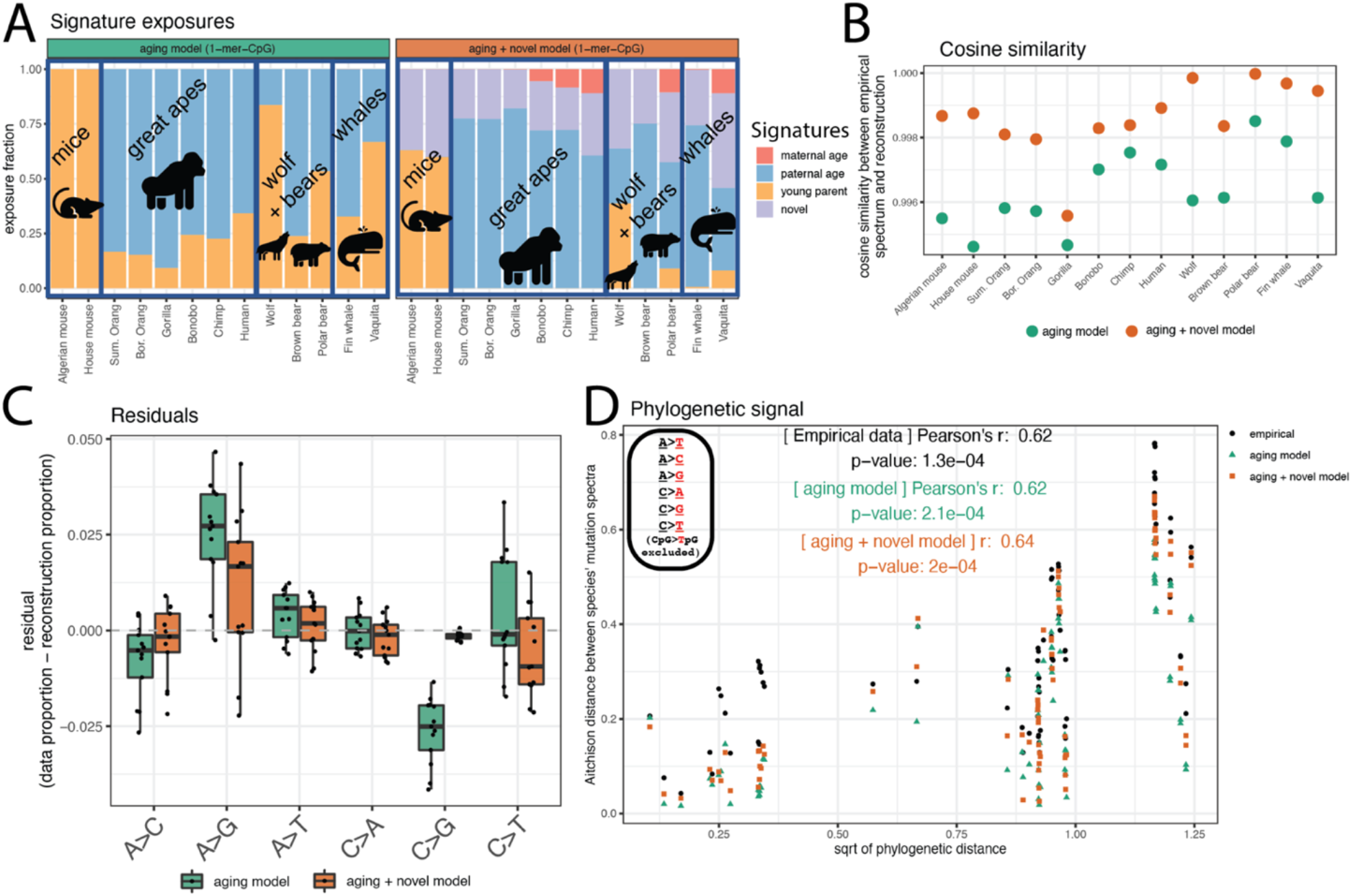
Human reproductive aging signatures can reconstruct the phylogenetic signal of the 1-mer mutation spectrum. **A)** Left panel: exposures to three reproductive aging signatures derived from human trio data in Jónsson et al. (2017) maternal age (red), paternal age (blue), young parents (gold). Right panel: exposures to a model containing an additional fourth novel signature extracted from the data (purple). Broad phylogenetic clades are outlined and labeled. **B)** Cosine similarity between the empirical spectrum and the spectrum reconstructed using either the aging signatures alone, or the aging signatures + novel signature. Higher cosine similarity indicates a better reconstruction of the data. **C)** Residuals (observed proportion - reconstructed proportion) for each spectrum. Boxplots are made up of the 6 1-mer mutation types (CpG>TpG mutations excluded) for all species. Negative residuals indicate the model reconstruction overestimating a proportion of a mutation type, positive residuals indicate an underestimate. The simple 3-signature aging model is indicated in green, the aging + additional novel signature in orange. **C)** Result of Mantel test measuring correlation between species’ reconstructed 1-mer-CpG spectra under each model (green triangles and orange squares) and the square root of phylogenetic distance, overlaid on results based on empirical spectra (black circles). *p*-values based on 9,999,999 permutations.

Despite these biased residuals, the reproductive aging model is able to capture the phylogenetic signal that is present in the empirical 1-mer mutation spectrum data (**Figure 7D**; reproductive aging model *r* = 0.62, *p* < 2.4e-4; empirical 1-mer (minus CpG>TpG) data *r* = 0.62, *p* < 1.3e-4). We found that adding an additional novel signature extracted from the data (**Figure S31**) improved the fit to the data and reduced the residual bias (**Figure 7B-C**), without changing the strength of phylogenetic signal captured by the reconstructed spectra (**Figure 7D**).

### Early reproduction is associated with similar mutational biases across disparate clades of mammals

It is particularly intriguing that the reproductive aging exposures assign most mouse mutations to the young parent signature (**Figure 7A**, left panel), possibly because mice reproduce at much younger ages (9 to 11 weeks of age) than any of the other species. The second- and third-highest young parent exposures are inferred in wolves and vaquita, which are the second- and third-youngest reproducers in our dataset on average, respectively (∼1 year and ∼5 years age at first reproduction). The mice, wolves, and vaquita all have lower Aitchison distances between their 1-mer and 3-mer mutation spectra than comparisons between species with similar or greater phylogenetic distance (**Figure S4**, **Figure S32**), which given their similar levels of exposure to the young parent signature, may be driven by similarities between these species’ reproductive times relative to other species.

We were able to reproduce this trend using de novo mutation data from mice (Lindsay et al. 2019) and an independently generated wolf polymorphism dataset (Mooney et al. 2023), consistently finding that the species’ 1-mer spectra that were most similar to that of mice were the wolf’s and vaquita’s 1-mer spectra (**Figure S32**). These results suggest that although the reproductive aging model cannot fully reconstruct these species’ polymorphism spectra in an unbiased way, the reproductive aging signatures are consistent with some of the major 1-mer-level differences that exist between different mammalian clades, which may make species with more similar reproductive strategies have more similar 1-mer spectra.

## Discussion

### Mammalian polymorphisms reveal a hierarchy of clade-specific mutational processes

We have extracted mutation spectra from polymorphism data in 13 species spanning 96 million years of mammalian evolution, standardizing these spectra to remove differences caused by composition of the accessible reference genome. Our pipeline is designed to facilitate comparison between genomically divergent species that have been sampled and sequenced using different methodologies; we expect that this flexibility will enable expansion of these analyses to more species as new data become available. Although we identified a few correlations between mutation spectrum features and bioinformatic confounding variables, we consistently found phylogenetic distance to be a stronger predictor of mutation spectrum divergence between species. Our results support the hypothesis that the mutation spectrum evolves over time due to slight increases in the rates of some mutation types and slight decreases in the rates of other mutation types.

Previous papers have used principal component analysis to demonstrate that closely related populations appear to have more similar mutation spectra than more distantly related populations (Harris and Pritchard 2017; Dumont 2019; Goldberg and Harris 2022). Here, we utilized the classical phylogenetic Mantel test to quantify the significance of this correlation, a technique that we recently used to quantify mutation spectrum evolution across the SARS-CoV-2 phylogeny (Bloom et al. 2023). We expect this test to scale well to future datasets containing even more species and populations, and it is adaptable for testing how much variation among mutation spectra is captured by particular mutational signature models or sub-spectra of interest.

We investigated several nested *k*-mer mutation spectra and found the strongest phylogenetic signal at the 3-mer context level. However, we also found that a small number of 7-mer mutation types exhibit hypermutability that varies conspicuously among species. In particular, the 7-mer TTTAAAA, whose A>T mutation rate is extremely elevated in humans, has a consistently high mutation rate across mammals but an exceptionally high mutation rate within whales. A previous study (Carlson et al. 2018) hypothesized that TTT**<u>A</u>**AAA>TTT**<u>T</u>**AAA hypermutability is caused by LINE-1 transposase activity, which preferentially cuts into specific genomic motifs in a manner that is susceptible to error-prone repair. Since LINE-1 elements have a documented pattern of hyperactivity in Minke whales (Ivancevic et al. 2016), the transposase nicking mechanism might explain the observed interspecies differences in the rate of this outlier mutation type. Although repetitive *k*-mers like TTTAAAA might have an elevated susceptibility to sequencing errors, the hypermutability of this mutation type does not appear to correlate with genome assembly quality or sequencing coverage.

### Power and limitations of polymorphism data for mutation spectrum analysis in non-model organisms

A limitation of our study is the fact that we inferred mutation spectra from polymorphisms, which typically exhibit systematic differences from de novo mutation spectra ascertained in the same species (Zhu et al. 2014; Carlson et al. 2018; Wang et al. 2022; Ragsdale and Thornton 2023). These polymorphisms are descended from mutations that occurred many generations ago and are impacted by GC-biased gene conversion (gBGC), which enriches the spectrum for A>G and A>C mutations while depleting it of C>A and C>T. Since the strength of gBGC is proportional to the effective population size, it has potential to create mutation spectrum differences between large, outbred populations (such as mouse populations) and more inbred species such as the vaquita porpoise. However, gBGC is not known to impact A>T or C>G mutations, or to influence the sub-spectra of any 1-mer mutation type. In contrast to this expectation, we consistently find phylogenetic signal in sub-spectra of mutations that each have the same gBGC selective modality (positively selected A>G plus A>C; negatively selected C>A plus C>T; neutrally evolving A>T plus C>G). This finding, combined with the correlation between effective population size and mutation spectrum divergence being weaker than that of phylogenetic distance, suggests that mammalian mutation spectrum variation must be driven by forces other than gBGC. Gao et al. (2023) recently used a similar partitioning strategy to confirm that gBGC cannot be the driver of most signals of human mutation spectrum evolution.

It is possible that these mutation spectra are confounded by selective forces other than GC-biased gene conversion, such as selection to preserve gene regulatory motifs that might be unique to particular lineages. This possibility will be important to investigate as gene regulatory grammar becomes better understood in non-model species.

Although polymorphisms are biased by evolutionary processes, they will likely continue to be indispensable for the study of mutation spectrum evolution given that comparably large numbers of de novo mutations are not possible to sample. A sample of five unrelated individuals per species allowed us to sample hundreds of thousands of variants and quantify context-dependent mutation spectra with higher precision than would be possible using de novo mutation sets that typically number in the high tens or low hundreds. This precision is essential given that normal germline mutation spectra are much more similar to one another than pathological cancer mutation spectra – all of the 1-mer and 3-mer mutation spectra extracted from species included in this study have pairwise cosine similarities in excess of 0.92. Although these differences are small in magnitude, our Mantel test results show that they exist along a robust hierarchy where more closely related species accumulate mutations in more similar sequence contexts.

### Existing models of germline mutational signature etiology do not fully capture phylogenetic signal

Mutational signature deconvolutions of cancer spectra typically assume that a model fits a dataset well if the cosine similarity between the model and the data is greater than 0.95 (Blokzijl et al. 2018; Alexandrov et al. 2020; Gori and Baez-Ortega 2020). Our results indicate that this threshold is likely too permissive for reconstructing germline mutational spectra, at least at the level of precision that is needed to capture the drivers of mammalian mutation spectrum evolution. We can see this by considering the SBS1+SBS5 clocklike cancer signature model, which captures none of the 3-mer spectrum’s phylogenetic signal yet fits each species’ spectrum with cosine similarity greater than 0.95.

Although the SBS1+SBS5 model and the reproductive aging model achieve similar cosine similarity fits to the data, the significant phylogenetic signal captured by the reproductive aging model suggests that it better encapsulates some of the forces that are driving mutation spectrum evolution. It is particularly intriguing that the mouse and the grey wolf have the shortest ages at first reproduction in our dataset and also the lowest exposures to the paternal and maternal aging signatures. However, if the parental age model captured all mutation spectrum variation among mammals, we would expect this model to explain all the variation of the 1-mer spectrum between species with unbiased residuals, which is not the case. This implies that either parental aging cannot explain all mutation spectrum variation between species, or the Jónsson et al. (2017) model does not fully capture the mutational effects of parental aging, perhaps due to the sparsity and bioinformatic complexity of the underlying human mutation data.

Importantly, it is not known how human reproductive aging affects the 3-mer mutation spectrum, as currently available de novo mutation data are too sparse to determine this with precision. It is possible that maternal and paternal aging signatures have different context dependencies that might explain the phylogenetic signal of the 3-mer and 5-mer spectra. In the absence of such data, our results suggest that the reproductive aging model may be an important contributor to mammalian mutation spectrum variation, particularly at the 1-mer level, but that additional factors such as mutator alleles or environmental mutagens may be needed to fully explain the 3-mer phylogenetic signal we observe. The correlation of 3-mer and higher-order mutation spectra with reproductive lifespan and age at first reproduction adds additional evidence that these life history features play a role in shaping mutation spectra. However, since these correlations are weaker than the mutation spectrum’s phylogenetic signal, they do not support the idea that life history is the primary driver of mutation spectrum differences between species.

In summary, the observed variation of the mutation spectrum across the phylogeny is consistent with theoretical expectations about a heritable polygenic trait. If germline mutations are generated by several different mutational processes and the rate of each process is subject to weak selective constraint, then as species evolve, each of their mutation spectra should perform a random walk through a multidimensional space. Other explanations of the data are also possible: to the extent that closely related species tend to inhabit similar environments and reproduce via similar strategies, environmental mutagens and conserved reproductive aging signatures might create phylogenetic signal that is consistent with our expectations of a mutator allele, particularly for the lower-dimensional 1-mer spectrum. To disentangle these possibilities, it will be necessary to sample mutation data from a wider variety of species that independently came to inhabit similar environments or reproductive niches, as well as species that recently came to inhabit new environments (as humans did). We anticipate that such mutation data will become increasingly available over the coming years, and that the methodology presented in this paper will allow for nuanced modeling of the mutagenic effects of genotype versus environment.

## Methods

### Data acquisition

#### Humans

Byrska-Bishop et al. (2022) sequenced 2,504 human genomes to 30x coverage (Illumina NovaSeq 6000). They mapped these data to the human genome assembly GRCh38 and called genotypes using *GATK HaplotypeCaller*, with variant quality score recalibration (VQSR) carried out as described in their paper. We downloaded the resulting genotype dataset in .*vcf* format (“1000 Genomes 30x on GRCh38” dataset) from the 1000 Genomes FTP site (https://www.internationalgenome.org/data-portal/data-collection/30x-grch38). We used superpopulation assignments for downstream analyses with the following sample sizes: African (AFR; *n* = 661), European (EUR; *n* = 503), South Asian (SAS; *n* = 489), East Asian (EAS; *n* = 504), Admixed Americans (AMR; *n* = 347).

#### Non-human great apes

Prado-Martinez et al. (2013) carried out whole genome sequencing to mean 25x coverage (Illumina HiSeq 2000) of 79 great ape individuals from the species chimpanzee (*Pan troglodytes*), bonobo *(Pan paniscus*), gorilla (*Gorilla gorilla* and *Gorilla beringei*), Sumatran orangutan (*Pongo abelii*), and Bornean orangutan (*Pongo pygmaeus*). They mapped the short-read sequencing data to the human genome hg18 (NCBI Build 36) and called genotypes using *GATK UnifiedGenotyper* with the following filter criteria: DP < (mean_read_depth/8.0) || DP > (mean_read_depth*3), QUAL < 33, FS > 26.0, MQ < 25, MQ0 ≥ 4 && ((MQ0 / (1.0 * DP)) > 0.1), and sites within 5bp of an indel (Prado-Martinez et al. 2013). We downloaded their processed genotype *.vcf* files from the Great Ape Genome Project (GAGP) (https://www.biologiaevolutiva.org/greatape/). For chimp, one individual (Pan_troglodytes_ellioti-Banyo) was excluded from downstream analyses as it was listed as low quality in the GAGP documentation. For the Western Lowland gorilla (*Gorilla gorilla gorilla*), two individuals listed in the GAGP documentation as being low quality were removed (Gorilla_gorilla_gorilla-X00108_Abe and Gorilla_gorilla_gorilla-KB7973_Porta). Additionally, individuals from the *Gorilla beringei graueri* and *Gorilla gorilla dielhi* subspecies were excluded to reduce population structure in the dataset (Gorilla_beringei_graueri-9732_Mkubwa, Gorilla_beringei_graueri-A929_Kaisi, Gorilla_beringei_graueri-Victoria, Gorilla_gorilla_dielhi-B646_Nyango).

After exclusion of these individuals, the resulting samples sizes were chimpanzee (*n* = 24), bonobo (*n* = 13), gorilla (*n* = 25), Sumatran orangutan (*n* = 5), and Bornean orangutan (*n* = 5).

#### Wild mice

Harr et al. (2016) sequenced mouse whole genomes to 11-26x coverage (Illumina HiSeq 2000) from natural populations of house mouse (*Mus musculus*), including the subspecies western house mouse (*M. m. domesticus*) and eastern house mouse (*M. m. musculus*) and Algerian mouse (*Mus spretus*). They also included publicly available data from the southeast-Asian house mouse (*M. m. castaneus*) in their genotype calling pipeline. They mapped reads to the mm10 mouse reference genome and called genotypes using *GATK* best practices, including VQSR. Sample sizes for each species/subspecies are *M. m. domesticus* (Mmd; *n* = 27), *M. m. musculus* (Mmm; *n* = 22); *M. m. castaneus* (Mmc; *n* = 10), *M. spretus* (Ms; *n* = 8).

#### Grey wolf

The Broad Institute sequenced 676 canids to 10-94x coverage (Morrill et al. 2022). The reads were mapped to the dog reference genome (canfam3) and genotypes were called using *GATK HaplotypeCaller*, with hard filters QD < 2.0, FS > 60.0, MQ < 40.0, MQRankSum < - 12.5, ReadPosRankSum < -8.0, and GQ < 14.0. We selected a subset of grey wolves (*Canis lupus*) from the dataset and one coyote (*Canis latrans*) to act as an outgroup for polarization (wolves: Wolf03, Wolf08, Wolf19, Wolf20, Wolf24, Wolf27, Wolf29, Wolf31, Wolf32, Wolf33, Wolf34, Wolf35, Wolf36, WO001_895, WO002_732, WO003_636, Wolf40, Wolf41, Wolf42, and coyote: cal_coy). The resulting sample size was grey wolf (*n* = 19).

#### Fin whale

Nigenda-Morales et al. (in press) sequenced 50 fin whale (*Balaenoptera physalus*) whole genomes (20x coverage; Illumina HiSeqX and NovaSeq6000) from two populations, the Eastern North Pacific (ENP) and the Gulf of California (GOC). They mapped reads to the outgroup minke whale genome (BalAcu1.0) and called genotypes using *GATK HaplotypeCaller*. Additionally, they mapped one humpback whale (*Megaptera novaeangliae*) and one blue whale (*Balaenoptera musculus*) individual to the minke whale genome for use in polarization. Variants were filtered using hard filters, with minimum genotype depth of 8, and maximum as 250% of the mean per-individual depth. Genotypes had to have a minimum Phred score of 20 and have 0.2 ≤ allelic balance ≤ 0.8 for heterozygous genotypes, allelic balance ≥ 0.9 for homozygous reference genotypes, and allelic balance ≤ 0.1 for homozygous alternate genotypes. They also filtered sites based on the *GATK* recommended hard filters (QD < 2.0, FS > 60.0, MQ < 40.0, MQRankSum < -12.5, ReadPosRankSum < -8.0, SOR > 3.0, and QUAL < 30). Sites where >75% of individuals were heterozygous were also excluded. They excluded six individuals that had low genotype quality, high degrees of admixture, or high kinship to other individuals from the dataset (ENPOR12, ENPCA01, ENPCA09, GOC010, GOC080, GOC111). We excluded one additional low-coverage individual (ENPAK28) from downstream analyses. The resulting sample sizes were fin whales from the Gulf of California (GOC; *n* = 17) and fin whales from the Eastern North Pacific (ENP; *n* = 26).

#### Vaquita porpoise

Robinson et al. (2022) sequenced 20 vaquita porpoise (*Phocoena sinus*) whole genomes from the Gulf of California to 60x coverage (Illumina HiSeqX). They mapped reads to the vaquita reference genome (mPhoSin1.pri, GCF_008692025.1) and called genotypes using *GATK HaplotypeCaller*. They filtered genotypes using the hard filters: QD < 4, FS > 60, MQ < 40, MQRankSum < -12.5, ReadPosRankSum < -8, SOR > 3. Sites at which >75% of individuals were heterozygous were excluded. Heterozygous sites with allele balance < 0.2 or > 0.8 were excluded, as were homozygous genotypes with allele balance > 0.9 or < 0.1. They filtered genotypes with depth less than 1/3 of the mean read depth or greater than 2x mean read depth of the individual. After we excluded five relatives of individuals left in the dataset (z0001663, z0004380, z0004393, z0004394, z0185383), the resulting sample size was vaquita (*n* = 15).

For all datasets, the genotype data were restricted to biallelic SNPs with the “PASS” designation, indicating that they have passed all filters.

### Polar bear and brown bear genotype calling

Since a unified dataset for polar bear (*Ursus maritimus*) and brown bear (*Ursus arctos*) was not available, we carried out genotype calling ourselves. Fastq files for polar bear and brown bear individuals (**Table S2**) were downloaded from the European Nucleotide Archive (ENA). Additionally, one black bear (*Ursus americanus*) individual was included for use in polarization (see below). Paired-end reads were mapped to the brown bear (Acc. # GCA_003584765.1) and polar bear (Acc. # GCA_000687225.1) genomes using *paleomix* (Schubert et al. 2014), which removes adapters and PCR duplicates and filters out reads with quality < 30. Variants were called for all scaffolds ≥ 1Mb, corresponding to 2.218Gb of sequence in the polar bear reference genome (out of a total genome size of 2.301 Gb) and 2.253Gb in the brown bear genome (out of 2.328Gb). Note that reads were mapped to the set of all scaffolds, but variants were only called for those ≥ 1Mb. using *GATK* (v. 3.7) *HaplotypeCaller*, with all sites emitted and sites with mapQ < 20 filtered out (Van der Auwera et al. 2013). Sites were then joint-genotyped across all individuals using *GATK GenotypeGVCFs*, emitting all sites (variant and invariant).

*GATK* hard filters were then used to filter sites, excluding variants that had: QD < 2.0, FS > 60.0, SOR > 3.0, MQ < 40.0. MQRankSum < -12.5, ReadPosRankSum < -8.0. Additionally, at the genotype level to be retained the individual genotype had to have GQ ≥ 20, genotype depth that is ≥ 8 and ≤ 250% of the mean depth of the individual, have 0.2 ≤ allelic balance ≤ 0.8 for heterozygous genotypes, allelic balance > 0.9 for homozygous reference genotypes, and allelic balance < 0.1 for homozygous alternate genotypes. Finally, sites at which > 75% of individuals were heterozygous were filtered out. After filtering, thirteen lower-coverage individuals had < 19.5Gb of sites passing filters, and so were excluded from downstream analyses (006_UARC_SW_BGI_brownbear_20105373,015_UARC_GEO_Ge 017_UARC_RUS_S235,020_UARC_AK_Uarc_EP040,021_UARC_ABC_Uarc_EP050040_UMAR_SVL_PB1_N23531,041_UMAR_SVL_PB10_N23997,042_UMAR_SVL_PB2_N23604,043_UMAR_SVL_PB3_N23719,044_UMAR_SVL_PB4_N23917,045_UMAR_SVL_P B6_N26028,047_UMAR_SVL_PB8_N7968,048_UMAR_SVL_PB9_N23985).

Brown bears were classified into two geographic groups–the Alaskan Admiralty, Baranof, and Chicagof (ABC) islands and Europe. Two individuals from Alaska or Montana were excluded as outliers originating from outside of these two main populations. After exclusion of these individuals, the resulting sample sizes were polar bear (*n* = 18) and brown bear from ABC Islands (*n* = 8) and brown bears from Europe (*n* = 6).

### Polarization

To assign mutation type (e.g., T**<u>A</u>**C > T**<u>G</u>**C), the ancestral allele state must first be determined. The software *mutyper* (DeWitt et al. 2023), which is used to assign mutation type, requires an ancestral genome fasta file in which the reference genome has been updated with ancestral states at polarized SNP positions (for example, if the reference genome contains an “A” at a particular site, but the ancestral state is determined to be “C”, the ancestral fasta should have a “C” at that position).

Ancestral genome fasta files were previously generated for the human genome as part of the 1000 Genomes Project using the inferred great ape phylogeny at each site (Auton et al. 2015) (details in that paper’s Supplement 8.3.1). Ancestral genome states were generated for the great ape species in Goldberg & Harris (2022) using parsimony. The remaining species had ancestral states assigned using *est-sfs* (Keightley and Jackson 2018), a program which takes SNPs from a focal species and one or more outgroups aligned to the same reference genome and uses parsimony and allele frequencies to estimate the probability that the major allele of the focal species is the ancestral state. Ancestral alleles can then be assigned probabilistically using a single draw from a binomial distribution per SNP, where the probability of success is the probability that the major allele is ancestral for the focal species: if the binomial draw yields ‘1’ (success) then the major allele is assigned as likely to be ancestral; if it yields ‘0’ then the minor allele is assigned as likely to be ancestral. Robinson et al. (2022) polarized the vaquita (*Phocoena sinus*; focal) in this manner with harbor porpoise (*Phocoena phocoena*; outgroup 1) and Indo-Pacific finless porpoise (*Neophocaena phocaenoides;* outgroup 2) as outgroups, and provided us with these previously unpublished data. We implemented *est-sfs* using the Kimura 2-parameter model and 10 maximum likelihood runs for the species in our dataset that had not been polarized by previous studies, using 1-2 outgroups depending on availability:

- polar bear (*U. maritimus*; focal), brown bear (*U. arctos*; outgroup 1), black bear (*U. americanus;* outgroup 2)
- brown bear (*U. arctos*; focal), polar bear (*U. maritimus*; outgroup 1), black bear (*U. americanus*; outgroup 2)
- Western European house mouse (*M. m. domesticus*; focal); southeastern Asian house mouse (*M. m. castaneus;* outgroup 1); Algerian mouse (*M. spretus;* outgroup 2)
- Algerian mouse (*M. spretus*; focal); Western European house mouse (*M. m. domesticus*; outgroup 1)
- Grey wolf (*C. lupus;* focal), coyote (*C. latrans;* outgroup1)
- Fin whale (*B. physalus;* focal), humpback whale (*Megaptera novaeangliae*; outgroup 1), blue whale (*Balaenoptera musculus*; outgroup 2)

### Genome masking

To avoid regions under selection, regions that may be susceptible to poor genome quality calls due to repeat content, and regions known to have very atypical mutational processes compared to the rest of the genome, we generated mask files for every genome in order to mask:

1) exons ± 10kb on either side of each exon
2) repeat regions identified using *RepeatMasker* (Smit et al. 2013)
3) low-complexity regions identified using *Tandem Repeat Finder* (*TRF*) (Benson 1999)
4) CpG islands identified using *CpGPlot* (Larsen et al. 1992) or the UCSC pipeline (http://genome.ucsc.edu/cgi-bin/hgTrackUi?g=cpgIslandExt).

For most reference genomes in our study, the coordinates of these regions were available for download from the UCSC genome browser (https://genome.ucsc.edu/). For vaquita and fin whale, these masks are not available on the UCSC browser but were previously annotated and provided by the data generators (Robinson et al. (2022) and Nigenda-Morales et al. (in press), respectively). We generated the remaining files as follows: for the brown bear and polar bear we ran *RepeatMasker* (with species database set to “carnivore”) (Smit et al. 2013). For polar bear, brown bear, and minke whale we ran *Tandem Repeat Finder* (*TRF*) (Benson 1999) to identify low-complexity regions using recommended parameters (and a maximum period size of 12 as in UCSC pipelines (http://genomewiki.ucsc.edu/index.php/TRF_Simple_Repeats) (parameters: match:2, mismatch: 5, delta: 7, PM: 80, PI: 10, minscore: 50, maxperiod: 12) and converted the output using a conversion script (https://github.com/Adamtaranto/TRF2GFF). Finally, for the brown bear, polar bear and minke whale genomes, we ran *EMBOSS CpGPlot* (Larsen et al. 1992) to identify CpG Islands using default parameters (window size: 100, minimum length of island: 200, minimum observed/expected ratio: 0.6, minimum percentage: 50).

We then combined all of these regions into one negative mask file in *.bed* format for each reference genome using *bedtools sort* and *bedtools merge* (Quinlan and Hall 2010). These regions were then masked out from all downstream analyses.

### Generating mutation spectra from polymorphism data

The mutation spectrum represents the distribution of relative abundances of different mutation types, sometimes calculated from all derived alleles present in one individual genome and sometimes calculated from all sites that are variable within a larger population sample. In the simplest form of the mutation spectrum, which we call the 1-mer spectrum, mutations are classified into 6 types (A>T, A>C, A>G, C>T, C>G, C>A) with DNA strand complements collapsed. This can be expanded into a 7-dimensional “1-mer+CpG spectrum” by separating C>T mutation types into those that are found in a CpG context and those that are not. CpG>TpG mutations can also be removed from the spectrum entirely (“1-mer-CpG spectrum”).

Finer-grained mutation spectra can be computed by subdividing these basic mutation types by their flanking sequence context. In the popular 3-mer spectrum, mutations are classified by their immediate 5’ and 3’ flanking basepairs, yielding 96 mutation types once reverse complements are collapsed (e.g., T**<u>C</u>**C>T**<u>G</u>**T, A**<u>A</u>**C>A**<u>T</u>**C, A**<u>C</u>**G > A**<u>T</u>**G, etc.). Higher-dimensional spectra are possible as well, including 5-mers (2bp on either side of the mutating base, e.g., TA**<u>C</u>**CT>TA**<u>T</u>**CT, resulting in 1,536 mutation types), or 7-mers (3bp on either side of the mutating base, e.g., TTA**<u>C</u>**CTA > TTA**<u>T</u>**CTA, resulting in 24,576 mutation types). These higher-dimensional spectra have considerably more mutation types, which can perhaps aid in the detection of more subtle mutation signatures, but which can also lead to issues of data sparsity.

For each individual, we estimated a nested series of spectra at the 1-mer, 1-mer+CpG, 1-mer-CpG, 3-mer, 5-mer and 7-mer levels from our polymorphism data, as described below.

Where sufficient individuals existed within a single population within a species, one population was chosen as representative for each species for downstream analyses: humans (African/AFR continental group), fin whale (Gulf of California population), and brown bear (ABC Islands population). First, to standardize sample sizes across datasets, five individuals were drawn at random from each population and used for all downstream analysis (**Table S1**). Sites that were missing genotypes across the selected individuals within a species were removed.

Each remaining variant’s 7-mer mutation context (3bp on either side of each SNP) was extracted using the function *mutyper variants* (DeWitt et al. 2023), restricting to biallelic SNPs with no missing data that are not fixed for the alternate allele and excluding the masked regions described above. In the human ancestral genome *.fasta* file, lower-case letters indicate sites with uncertain polarization; these were excluded using the --strict setting of *mutyper variants*. To enable the correction of these mutation spectra for the *k*-mer content of each species’ reference genome, we also used the function *mutyper targets* to count the number of times each 7-mer was observed as part of each species’ ancestral state *.fasta* file.

To generate individual-level mutation spectra we calculated each individual’s 7-mer mutation spectrum (counts of each 7-mer mutation type) using the function *mutyper spectrum*. SNPs with derived alleles appearing in more than one of these five individuals were randomly assigned to a single individual’s mutation spectrum using the –*randomize* parameter to prevent shared variation from driving similarity between related individuals. These spectra were used to perform principal component analysis as described below.

To summarize the mutation spectrum of each population sample as a whole, we generated per-population spectra using *mutyper spectrum* with the –population parameter, which is the equivalent of summing the randomized per-individual spectra.

To generate lower-dimensional mutation spectra, the 7-mer spectra generated per-individual and species described above were collapsed down into their 5-mer, 3-mer or 1-mer content (e.g., all 7-mers containing T**<u>A</u>**A>T**<u>G</u>**A as their central 3-mer were summed up to get the count of that mutation type for the 3-mer mutation spectrum).

### Correcting for genome content and amount of genetic diversity

To compare mutation spectra between individuals and species, the mutation spectra must be corrected for several factors, including genomic content and genetic diversity. The 7-mer, 5-mer, 3-mer and 1-mer spectra were corrected in the same manner, as described below.

Genomic *k*-mer content may differ across species. For example, one species may have more “ACC” 3-mers in its genome, causing more A**<u>C</u>**C>A**<u>N</u>**C mutations to accumulate due to a larger ACC target size rather than a higher A**<u>C</u>**C>A**<u>N</u>**C mutation rate). To correct for differences in genomic content, we transformed the SNP count *x_m→j,A_* of 7-mer mutation type *m → j (k*-mer *m* mutates to *k*-mer *j*) into a rescaled SNP count *x^(r)^_m→j,A_* that is what we would expect to observe if the mutation rate were unchanged but the fraction of the ancestral *k*-mer of mutation type *m* were changed to match the human reference genome:

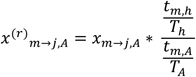

where *t_m,A_* is the number of times the ancestral *k*-mer of mutation type *m* is observed in species A’s reference genome, *T_A_* is the total target count (sum of all *k*-mer targets) in species A’s reference genome, *t_m,h_* is the number of times the ancestral *k*-mer of mutation type *m* is observed in the human reference genome, and *T_h_* is the total target count (sum of all *k*-mer targets) in the human reference genome.

After rescaling all counts to the same genomic content, the mutation counts are normalized to fractions. Then, multinomial downsampling is carried out using the *rmultinom* function in *R* to sample the number of SNPs found in the lowest diversity species using the fraction of each mutation type as the multinomial probabilities. The vaquita had the lowest diversity, and so all species were downsampled to match the ∼130,000 SNPs observed in the vaquita dataset. This downsampling corrects for different levels of diversity across species caused by differences in effective population size, overall mutation rate, or generation time, and adds more noise to the spectra of species with higher diversity so that they are comparable to the noisier spectra of lower diversity species. At these lower numbers of SNPs, some mutation categories have mutation counts of 0 (particularly in the high-dimensional 7-mer spectrum, and to a lesser extent, the 5-mer spectrum).

### Center log ratio (CLR) transformation of mutation spectrum data

The mutation spectrum is an example of a *compositional* data type, meaning that it is a distribution vector constrained to sum to 1. This internal dependency structure can cause downstream analyses of compositional data to be confounded by spurious correlations, but this problem can be largely eliminated by applying a geometrical transformation called the centered log ratio transform (CLR) that was developed by John Aitchison (Aitchison 1986).

We transformed our mutation spectrum data using the function *clr* from the *compositions R* package (van den Boogaart and Tolosana-Delgado 2008). The CLR transformation divides each value within the compositional spectrum by the geometric mean of the spectrum, then takes the natural logarithm. CLR-transformed spectra are used to compute the PCA and distance analyses described below. Because the Aitchison distance represents the difference between two logarithmic values, it makes comparisons between species’ mutation counts proportional, instead of the absolute differences used in calculated un-transformed Euclidean distance. This has the additional advantage that variation in the most abundant mutation types does not dominate the differences between species to the degree that they can dominate distances between raw mutation spectrum vectors (**Figure S33**).

Since the CLR transformation is incompatible with mutation counts of zero, we regularized our data by adding a pseudocount of 1 to the number of mutations observed within each type category in each species. This regularization is done separately for 1-mer, 3-mer, 5-mer and 7-mer spectra. It contributes very little to 1-mer and 3-mer spectra since these have relatively high mutation counts per mutation type, even after downsampling, but is more critical for 5-mer and particularly 7-mer spectra which are sparser and therefore have many missing mutation types after downsampling.

We then converted the mutation counts to approximate ‘rates’ by dividing each count by the human-rescaled genomic target size of each k-mer.

An apparent limitation of this approach is that the final distances between species’ mutation spectra will depend on the particular *k*-mer composition chosen for normalization (specifically, the human reference genome composition). However, we found that the CLR distance between mutation spectra is actually not dependent on this choice of *k*-mer composition as long as the species’ spectra are rescaled to the same target sizes; an algebraic proof in **Note S2** shows that we could normalize our variant counts to any other nonzero *k*-mer distribution and obtain the same distance matrix. This proof also shows that as long as the species’ spectra are rescaled to the same genomic target composition, that same result would be obtained using mutation counts, mutation fractions, or mutation ‘rates’ (counts/targets) to represent the mutation spectrum (**Figure S34**). This implies that our results are not intrinsically sensitive to the genomic *k*-mer composition chosen to rescale all species’ spectra to, nor to the particular set of species, genome assemblies or accessibility masks chosen for our study, except to the extent that they cause inclusion or exclusion of genomic regions with very different mutation spectra or systematic errors.

In addition to using the CLR, we also repeated our results using the ILR transform (isometric log-ratio) (Egozcue et al. 2003), with qualitatively similar results.

### Principal component analysis

Principal component analysis was carried out on the per-individual mutation spectra (for the five randomly sampled individuals from each species) using *prcomp* in *R* with variables shifted to be 0-centered and scaled to have unit variance (scale = *T*, center = *T*). The PCA results were then plotted for every combination of PCs 1, 2 and 3 using *autoplot* from the *ggfortify R* package (Tang et al. 2016), coloring each point by its species identity. PCA loadings showing the contributions of each mutation type to the first two variance components are plotted below the main PCA figure.

### Aitchison distance (Euclidean distance of CLR-transformed vectors)

Aitchison distance (the Euclidean distance between clr-transformed compositional vectors) was calculated between every pair of per-species spectra generated above using the *dist* function in R.

### Phylogenetic distance

To calculate the phylogenetic distances between species, we used the maximum likelihood tree that Upham et al. (2019) generated using *RAxML* (Stamatakis 2014) from an alignment of 31 genes across 4,098 mammal species, including all the species included in our study. Upham et al. rooted the tree with *anolis* as its outgroup and constructed using the GTRCAT model within *RAxML*, with five independent replicates. The branch lengths of the *RAxML* tree represent the expected number of substitutions per site in the alignment. We subsetted the tree to include only our study species ("Balaenoptera_physalus","Phocoena_sinus","Ursus_maritimus","Ursus_arctos","Mus_musculus","Mus_spretus","Homo_sapiens","Pongo_pygmaeus","Pongo_abelii","Pan_troglodytes","Pan_pa niscus","Gorilla_gorilla","Canis_lupus") using the *keep.tip* function in the R package *ape* (Paradis and Schliep 2019), and then calculated pairwise patristic distance (the sum of branch lengths between species) for every pair of species using the *cophenetic* function in *ape*. It is worth noting that the branch leading to the mouse species is very long (**Figure 2A**), a known feature of trees based on genetic distance that include rodents, likely due to their short generation times.

Additionally, we generated a second tree for our species using the tool *TimeTree* (Kumar et al. 2022) in which branch lengths represent divergence times between species scaled in millions of years (**Figure S1**). In this tree, the rodent branch is not longer than the other branches as the tree is scaled by time rather than genetic distance.

### Cosine similarity

We also measured cosine similarity between the non-transformed proportional mutation spectra, as this is a commonly used metric in cancer biology for comparing mutation signatures and spectra, using *sigfit’s cosine_sim* function (Gori and Baez-Ortega 2020).

### Mantel test for phylogenetic signal

Many methods, including Blomberg’s *K* (Blomberg et al. 2003), have been developed to quantify phylogenetic signal: the degree to which a trait varies continuously across a phylogeny. However, few of them are capable of comparing high-dimensional traits, since they generally require each trait to be summarized by a single number at each tip of the phylogeny. Since each species’ mutation spectrum is multidimensional and cannot be easily reduced to a single value, we are restricted to using a test for phylogenetic signal that is based on pairwise distances between tips. One test satisfying this constraint is the Mantel test for phylogenetic signal. The Mantel test calculates the correlation between two symmetrical distance matrices (one of pairwise distances in trait values, another of pairwise phylogenetic distances), and then permutes one matrix and recalculates the correlation. This is repeated for thousands of permutations, and a *p*-value is calculated based on the number of permutations which produce a correlation coefficient that equals or exceeds that of the real data. The Mantel test is underpowered compared to other tests for phylogenetic signal (Harmon and Glor 2010; Hardy and Pavoine 2012) but does not suffer from excess Type I error (Harmon and Glor 2010), even though the comparisons are not independent. It is important to note that the Mantel test, while an appropriate test for phylogenetic signal as performed here, should *not* be used in cases when two traits are being correlated without taking underlying phylogenetic signal into account (e.g. body weight and metabolism), as in that case it would suffer from excess Type I error (Harmon and Glor 2010; Guillot and Rousset 2013).

We tested for phylogenetic signal by quantifying the correlation between the pairwise Aitchison distance measured between species’ mutation spectra and the square root of the pairwise phylogenetic distance calculated from either the *RAxML* tree (genetic distance) or the timetree (distance in evolutionary time). We used the square root of phylogenetic distance because under a Brownian motion model the trait distance is expected to scale linearly with the square root of phylogenetic distance (Hardy and Pavoine 2012).

We used the *mantel* function from the *VEGAN* (Dixon 2003) package in R to carry out the Mantel test. We calculated the Pearson correlation coefficient between genetic distance and mutation spectrum distance for each of 9,999,999 permutations. If no permutation yields a correlation coefficient higher than the correlation coefficient of the non-permuted data, *VEGAN* returns a *p*-value of 1 / (1+9,999,999) = 1e-7. We carried out a separate Mantel test for each mutation spectrum (1-mer, 1-mer+CpG, 1-mer-CpG, 3-mer, 5-mer and 7-mer).

To confirm that our results were not due to the choice of a substitution-based tree, we repeated our analyses using cophenetic distances from the ultrametric time tree described above.

To confirm that our results were not due to how the data were polarized (ancestral states determined), we repeated the analysis using ‘folded’ mutation spectra, in which counts of reciprocal mutation types are added together (so that A**C**T > A**A**T and A**A**T>A**C**T mutations are not counted as separate categories, but are added together, meaning that mispolarization will not impact the results).

We also repeated our Mantel tests using a different distance measure between spectra: instead of Aitchison distance, we used the commonly-used metric cosine distance (1-cosine similarity) between spectra.

We additionally stratified the 3-mer, 5-mer and 7-mer spectra by central mutation type (treating all A>T *k*-mers separately from C>T *k*-mers when normalizing relative mutation rates, etc.) and carried out the Mantel test for distances calculated based on each central mutation type’s *k*-mers. We set a significance threshold of α = 0.05/6 for these tests to correct for multiple testing (6 central mutation types).

We also stratified the 3-mer, 5-mer and 7-mer spectra into three mutation categories depending on whether the ancestral and derived base pairs are “weak” (A:T) versus “strong” (G:C). This creates a class of weak-to-strong mutations that are positively selected by GC-biased gene conversion (A>C and A>G), a class of strong-to-weak mutations that are negatively selected (C>A and C>T), and a GC-conservative class that is neutrally evolving with respect to gBGC (A>T and C>G). We repeated the Mantel test within these stratified categories to verify that the mutation spectrum still exhibits phylogenetic signal after removal of the component that might be the result of gBGC. We set the significance threshold as α = 0.05/3 for these tests (three BGC categories).

In order to determine how much phylogenetic signal is contained in the 5-mer spectrum beyond what is driven by the phylogenetic signal of the central 3-mer, we randomized the counts of 5-mers within each central 3-mer category. We first stratified our 5-mer dataset by central 3-mer category (e.g. the A**<u>A</u>**A>A**<u>T</u>**A 3-mer category contains 16 5-mers: AA**<u>A</u>**AA>AA**<u>T</u>**AA, TA**<u>A</u>**AA>TA**<u>T</u>**AA, CA**<u>A</u>**AC>CA**<u>T</u>**AC, etc.) and calculated the relative proportion of each of the 5-mers target size in the genome within a 3-mer category. We then carried out multinomial sampling to assign each of the total number of mutations in that 3-mer category to a 5-mer mutation type, using its relative target proportion in genome as its multinomial probability of occurring. Thus, mutations are assigned to 5-mers solely based on their relative k-mer frequency in the genome, not due to any underlying differences in mutation rate. This removes any possible phylogenetic signal that could exist due to differences in 5-mer mutation rate, with any residual phylogenetic signal driven by the phylogenetic signal of the 5-mers’ central 3-mer. We generated 5,000 of these control spectra per species, then compared the distribution of Pearson’s *r* values between these control spectra and phylogenetic distance with the *r* value of the empirical 5-mer spectrum. We calculated a *p*-value as [(the number of randomized control spectra with *r* >= empirical) +1] / (5000 randomized datasets +1). We repeated this procedure for the 7-mer spectrum, randomizing 7-mer counts within each central 5-mer category.

To determine whether sparsity of 7-mers was driving the lack of phylogenetic signal beyond the 5-mer level, we repeated these analyses excluding the lowest-diversity species/populations in our dataset (removing the vaquita and polar bear and substituting the higher diversity Eastern North Pacific fin whales for the Gulf of California fin whales), resulting in the minimum number of SNPs in our dataset being ∼890,000 instead of ∼130,000. However, even with this higher number of SNPs (though reduced power due to fewer species), the 7-mer spectrum did not show a higher degree of phylogenetic signal than when randomized across 5-mer categories.

### *K_mult_* test for phylogenetic signal

We also tested for phylogenetic signal using a multivariate version of Blomberg’s *K*, called *K_mult_* (Adams 2014), as implemented in the *physignal* command in the R package *geomorph* (v. 4.0.5) (Adams et al. 2016; Baken et al. 2021) with 999 permutations.

### Other possible sources of phylogenetic signal

We identified several bioinformatic “phenotypes” (such as genome assembly quality) that vary between species, as well as several biological phenotypes that are suspected to impact mutagenesis, and used the Mantel test to determine whether any of these confounders has as much phylogenetic signal as the mutation spectrum itself. We reasoned that any potential confounder that exhibits less phylogenetic signal than the mutation spectrum can be ruled out as a major driver of mutation spectrum variation between species.

To measure key bioinformatic phenotypes, we first calculated the average per-individual SNP sequence depth of the five individuals chosen at random from each species’ dataset using *vcftools --depth* (Danecek et al. 2011) (the vaquita *.vcf* file did not contain depth information, so we used the average per-individual depth reported by Robinson et al. (2022)), and downloaded reference genome contig and scaffold N50 values from NCBI. To explore some biological phenotypes that might impact mutagenesis, we obtained estimates of age at first reproduction (in days) and reproductive lifespan (in days) for each species from (Pacifici et al. 2013). Vaquita was not included in that dataset and so we used estimates for harbor porpoise (*Phocoena phocoena*) as a proxy. Humans were also not included in that dataset, and so we added rough estimates of age at first reproduction and lifespan of age 22 years (8030 days) and ∼18 years of reproductive lifespan (6570 days). We then calculated the absolute value difference between every pair of species for each of these technical and biological phenotypes, and carried out the Mantel test to quantify correlation between these distances and phylogenetic distance (99,999 permutations). Note that fewer permutations were needed for these tests than the Mantel tests for phylogenetic signal of the mutation spectrum, as no variable reached the *p*-value threshold (1-e5) that would have necessitated additional permutations.

We then used the Mantel test to directly measure correlation between these technical and biological confounding variables and the mutation spectrum, in both a phylogeny-unaware and phylogeny-aware manner. For the phylogenetically-unware manner (in which shared phylogenetic signal could contribute to a correlation between the variables and the mutation spectrum), we carried out the Mantel test as described above, with 99,999 permutations. For the phylogenetically-aware method, we used the *PhyloMantel* function from the *evolqg R* package (v. 0.2-9) (Melo et al. 2016), to carry out the Mantel test with phylogenetic permutations (in which more closely related species are more likely to be permuted). We could then compare the results from both of these approaches to see how much of the correlation between these variables and our species’ mutation spectra appears to be due to shared phylogenetic signal.

### Enrichment or depletion of particular *k*-mers

To look for lineage-specific enrichment or depletion of particular *k*-mer mutation types, we calculated the relative mutation rate of each mutation type within each species based on their original projected mutation spectrum counts (not rescaled, downsampled, or regularized). The relative rates were calculated as the rate of each particular *k*-mer (e.g. TTT**<u>A</u>**AAA>TTT**<u>T</u>**AAA count divided by the target size of TTTAAAA targets in the genome) divided by the rate of its central base pair mutation type (count of A>T mutations divided by total accessible A bases in the genome). The ratio of these two rates indicates whether a particular *k*-mer is enriched (ratio > 1) or depleted (ratio < 1) relative to its central 1-mer rate. We tested whether each of these ratios differs significantly from 1 using a two-sided Fisher’s exact test (*fisher.test*() function in *R*) (this test was chosen over the Chi-Squared test because downsampled counts of some *k*-mers may be below 5). We then plotted the -log(10) *p*-values for each *k*-mer and looked for outlier *k*-mers that exceeded Bonferroni-corrected significance thresholds.

### Using prior mutational models to explain phylogenetic signal

We used the R package *sigfit* (Gori and Baez-Ortega 2020) to fit *a priori* mutation signatures to our germline mutation spectra. *sigfit* quantifies the goodness of fit of each signature decomposition as the cosine similarity between the input mutation data and the mutation counts expected given the output signature model. To use *sigfit*, we first reverse-complemented our 1-mer and 3-mer mutation types that have “A” as the ancestral allele into the equivalent mutation types where “T” is the ancestral allele in keeping with mutation labeling conventions from the COSMIC database (Tate et al. 2019). Since COSMIC signatures are defined relative to human genome composition, we used the species’ mutation counts that had been rescaled based on human genome content.

We fit two COSMIC cancer signatures to our 3-mer spectrum data, SBS1 and SBS5, which have been previously found in somatic and germline polymorphism data across species (multinomial model; 10,000 iterations). We then fit a more complex model in which *sigfit* extracts a third novel 3-mer mutation signature from the data while simultaneously fitting SBS1 and SBS5 using the *fit_extract_signatures* function (multinomial model; 10,000 iterations; adapt_delta = 0.99).

We then inferred mutation signatures related to human reproductive aging from Poisson regressions carried out by Jónsson et al. (2017) from Icelandic human trio data, which measured the effects of maternal and paternal age on de novo mutation counts for each 1-mer+CpG mutation class (C>A, C>G, C>T, CpG>T, T>A, T>C, T>G in Jonsson et al.). As in Wang et al. (2023) and Gao et al. (2023), we excluded CpG>TpG sites from the signatures we inferred. We took the maternal and paternal age slope and intercept values from Jonsson et al.’s Poisson regressions (located in Table S9 in Jonsson et al.) and used them to infer three mutational signatures. First, a paternal age signature was obtained by dividing the inferred paternal age slope for a particular mutation type (e.g., C>A) by the sum of slopes for all mutation types. The paternal age mutational signature is the 6-dimensional vector of these mutation fraction slopes. A mutational signature of maternal aging was calculated in the same manner using the inferred slopes describing the association between maternal age and each component of the de novo mutation spectrum. Finally, we generated a “young parent” signature that is representative of the mutations occurring in the offspring of two parents who reproduce directly after puberty. Such offspring are expected to be affected by mutations that occur during their own embryonic development as well as the development of their parents’ gametes prior to puberty, but are not expected to have accumulated mutations during cell division of mature spermatocytes or during the post-puberty aging of male or female gametes. We calculated this signature by using the Jonsson et al. regression equation to obtain expected counts of each mutation type (in the following example, C>A mutations) in a child of two 13-year-old parents (a typical puberty age in humans):

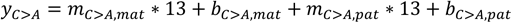

where *y_c>A_* is the expected total count of C>A mutations observed in the child, m_c>A,mat,_ is the maternal slope for C>A mutations, *b_c>A,mat_*, is the maternal intercept for C>A mutations, *m_c>A,pat_*, is the paternal slope for C>A mutations, and *b_c>A,pat,_* is the paternal intercept for C>A mutations, all from Jonsson et al.’s Table S9. The “young parent” mutation signature was then calculated by normalizing *y* for each mutation type by the sum of *y* across all mutation types, resulting in the expected mutation spectrum (under the Jonsson, et al. model) of a child conceived when its parents are pubescent.

We then fit these three parental age mutation signatures to our empirical 1-mer spectrum (minus CpG>TpG mutations, which we call the “1-mer-CpG spectrum”) using *sigfit* (multinomial model; 10,000 iterations). We then fit a more complex model using *fit_extract_signatures* in which *sigfit* inferred a fourth novel 1-mer-CpG signature from the empirical data while simultaneously fitting the three parental aging signatures (multinomial model; 10,000 iterations; adapt_delta = 0.99).

For each of the four models described above (“SBS1 + SBS5”, “SBS1 + SBS5 + novel”, “aging”, “aging + novel”), we calculated cosine similarities between each species’ mean reconstructed mutation spectrum (based on mutation signatures and exposures inferred by *sigfit*) and their empirical spectrum. We calculated per-species per-mutation-type residuals by subtracting the reconstructed mutation fraction from the empirical fraction (data - reconstruction).

To measure whether the reconstructions accurately capture phylogenetic signal, we carried out Mantel tests (9,999,999 permutations) to measure the correlation between species’ reconstructed spectra and the square root of their phylogenetic distance. We then compared these results to Mantel test results of the correlation between the empirical 1-mer-CpG and 3-mer spectra and phylogenetic distance.

### Investigating mouse-wolf spectra similarity

To confirm that the similarities we observe between the mouse and wolf spectra are not dataset specific or due to demographic history, we acquired two additional datasets: one dataset of mouse de novo mutation (DNM) 1-mer spectra (Lindsay et al. 2019) from whole genome sequencing of family trios, and a second wolf polymorphism dataset (Mooney et al. 2023) based on whole genome sequencing at the population level (which we call “wolf (UCLA)” as it was generated by a research group at that university, as opposed to “wolf (Broad)” which we used in our main analyses which was generated by the Broad Institute). We measured the Aitchison distance between each of these datasets’ 1-mer spectra to those of all our other species datasets. The mouse DNM data had elevated Aitchison distance compared to all other interspecies comparisons, likely due to excess noise in the 1-mer spectrum due to sparse data, and/or to systemic difference between DNMs and polymorphisms. However, reassuringly, the closest polymorphism-based spectra to the mouse DNMs were our two mouse polymorphism datasets (house mouse and Algerian mouse), and remarkably, the next-closest were the two wolf datasets and the vaquita porpoise, confirming the similarities between these species’ 1-mer spectra both at the DNM and polymorphism level, and across datasets.

## Code & data availability

Code used to generate spectra and analyses are available on https://github.com/harrispopgen/mammal_mutation_spectra. Mutation spectra with different transformations, genome masking bed files, polarized vcf files for species polarized in this study, ancestral fasta files, and reproductive aging mutation signatures will be made available on Dryad.

## Author contributions

ACB and KH conceived the study. ACB carried out all analyses and generated all figures, with supervision from KH. JR generated, processed, and polarized the vaquita data. ML, SN-M, and AME generated and processed the fin whale data. JR provided insights and code for ancestral polarization and provided insights during study design. ACB and KH wrote the manuscript, with input from all co-authors.

## Acknowledgements

ACB was supported by the National Institute of Health (NIH) “Biological Mechanisms of Healthy Aging” training program (T32 AG066574). KH received support from NIH/NIGMS grant R35GM133428, a Burroughs Wellcome Fund Career Award at the Scientific Interface, a Searle scholarship, a Pew Scholarship, and a Sloan Fellowship. We are very grateful for advice on analyses from Will DeWitt, Michael Goldberg, Pengyao Jiang, and Joe Felsenstein. We are grateful for detailed comments and suggestions on the manuscript from Joshua Schraiber. We sincerely thank those who shared data and insights with us, particularly: Phil Morin, Barbara Taylor, Lorenzo Rojas-Bracho, and the Southwest Fisheries Science Center’s Marine Mammal and Sea Turtle Research Collection, Eline Lorenzen, Mick Westbury, Jorge Urban, Frederick Archer, Kirk Lohmueller, Eduardo Amorim, Robert Wayne, and Daniel Promislow.

## Works cited

Abegglen LM, Caulin AF, Chan A, Lee K, Robinson R, Campbell MS, Kiso WK, Schmitt DL, Waddell PJ, Bhaskara S, et al. 2015. Potential Mechanisms for Cancer Resistance in Elephants and Comparative Cellular Response to DNA Damage in Humans. JAMA [Internet] 314:1850–1860. Available from: https://doi.org/10.1001/jama.2015.13134

Adams CJ, Conery M, Auerbach BJ, Jensen ST, Mathieson I, Voight BF. 2023. Regularized sequence-context mutational trees capture variation in mutation rates across the human genome. :2022.10.14.512160. Available from: https://www.biorxiv.org/content/10.1101/2022.10.14.512160v2

Adams DC. 2014. A Generalized K Statistic for Estimating Phylogenetic Signal from Shape and Other High-Dimensional Multivariate Data. Syst. Biol. [Internet] 63:685–697. Available from: https://doi.org/10.1093/sysbio/syu030

Adams DC, Collyer M, Kaliontzopoulou A, Sherratt E. 2016. Geomorph: Software for geometric morphometric analyses.

Adams DC, Collyer ML. 2018. Multivariate Phylogenetic Comparative Methods: Evaluations, Comparisons, and Recommendations. Syst. Biol. [Internet] 67:14–31. Available from: https://doi.org/10.1093/sysbio/syx055

Adams DC, Collyer ML. 2019. Phylogenetic comparative methods and the evolution of multivariate phenotypes. Annu. Rev. Ecol. Evol. Syst. 50:405–425.

Aggarwala V, Voight BF. 2016. An expanded sequence context model broadly explains variability in polymorphism levels across the human genome. Nat. Genet. [Internet] 48:349–355. Available from: https://www.nature.com/articles/ng.3511

Aikens RC, Johnson KE, Voight BF. 2019. Signals of Variation in Human Mutation Rate at Multiple Levels of Sequence Context. Mol. Biol. Evol. [Internet] 36:955–965. Available from: https://doi.org/10.1093/molbev/msz023

Aitchison J. 1986. The Statistical Analysis of Compositional Data. Springer Netherlands

Alexandrov LB, Jones PH, Wedge DC, Sale JE, Campbell PJ, Nik-Zainal S, Stratton MR. 2015. Clock-like mutational processes in human somatic cells. Nat. Genet. [Internet] 47:1402–1407. Available from: https://www.nature.com/articles/ng.3441

Alexandrov LB, Kim J, Haradhvala NJ, Huang MN, Tian Ng AW, Wu Y, Boot A, Covington KR, Gordenin DA, Bergstrom EN, et al. 2020. The repertoire of mutational signatures in human cancer. Nature [Internet] 578:94–101. Available from: https://www.nature.com/articles/s41586-020-1943-3

Alexandrov LB, Nik-Zainal S, Wedge DC, Aparicio SAJR, Behjati S, Biankin AV, Bignell GR, Bolli N, Borg A, Børresen-Dale A-L, et al. 2013. Signatures of mutational processes in human cancer. Nature [Internet] 500:415–421. Available from: https://www.nature.com/articles/nature12477

Anderson-Trocmé L, Farouni R, Bourgey M, Kamatani Y, Higasa K, Seo J-S, Kim C, Matsuda F, Gravel S. 2020. Legacy Data Confound Genomics Studies. Mol. Biol. Evol. [Internet] 37:2–10. Available from: https://doi.org/10.1093/molbev/msz201

Auton A, Abecasis GR, Altshuler DM, Durbin RM, Abecasis GR, Bentley DR, Chakravarti A, Clark AG, Donnelly P, Eichler EE, et al. 2015. A global reference for human genetic variation. Nature [Internet] 526:68–74. Available from: https://www.nature.com/articles/nature15393

Baken EK, Collyer ML, Kaliontzopoulou A, Adams DC. 2021. geomorph v4.0 and gmShiny: Enhanced analytics and a new graphical interface for a comprehensive morphometric experience. Methods Ecol. Evol. [Internet] 12:2355–2363. Available from: https://onlinelibrary.wiley.com/doi/abs/10.1111/2041-210X.13723

Benson G. 1999. Tandem repeats finder: a program to analyze DNA sequences. Nucleic Acids Res. [Internet] 27:573–580. Available from: https://doi.org/10.1093/nar/27.2.573

Bergeron LA, Besenbacher S, Zheng J, Li P, Bertelsen MF, Quintard B, Hoffman JI, Li Z, St. Leger J, Shao C, et al. 2023. Evolution of the germline mutation rate across vertebrates. Nature [Internet] 615:285–291. Available from: https://www.nature.com/articles/s41586-023-05752-y

Blokzijl F, Janssen R, van Boxtel R, Cuppen E. 2018. MutationalPatterns: comprehensive genome-wide analysis of mutational processes. Genome Med. [Internet] 10:33. Available from: https://doi.org/10.1186/s13073-018-0539-0

Blomberg SP, Garland JR. T, Ives AR. 2003. Testing for Phylogenetic Signal in Comparative Data: Behavioral Traits Are More Labile. Evolution [Internet] 57:717–745. Available from: https://onlinelibrary.wiley.com/doi/abs/10.1111/j.0014-3820.2003.tb00285.x

Bloom JD, Beichman AC, Neher RA, Harris K. 2023. Evolution of the SARS-CoV-2 Mutational Spectrum. Mol. Biol. Evol. [Internet] 40:msad085. Available from: https://doi.org/10.1093/molbev/msad085

van den Boogaart KG, Tolosana-Delgado R. 2008. “compositions”: A unified R package to analyze compositional data. Comput. Geosci. [Internet] 34:320–338. Available from: https://www.sciencedirect.com/science/article/pii/S009830040700101X

Bromham L, Hua X, Lanfear R, Cowman PF. 2015. Exploring the Relationships between Mutation Rates, Life History, Genome Size, Environment, and Species Richness in Flowering Plants. Am. Nat. [Internet] 185:507–524. Available from: https://www.journals.uchicago.edu/doi/abs/10.1086/680052

Byrska-Bishop M, Evani US, Zhao X, Basile AO, Abel HJ, Regier AA, Corvelo A, Clarke WE, Musunuri R, Nagulapalli K, et al. 2022. High-coverage whole-genome sequencing of the expanded 1000 Genomes Project cohort including 602 trios. Cell [Internet] 185:3426–3440.e19. Available from: https://www.cell.com/cell/abstract/S0092-8674(22)00991-6

Cagan A, Baez-Ortega A, Brzozowska N, Abascal F, Coorens THH, Sanders MA, Lawson ARJ, Harvey LMR, Bhosle S, Jones D, et al. 2022. Somatic mutation rates scale with lifespan across mammals. Nature [Internet] 604:517–524. Available from: https://www.nature.com/articles/s41586-022-04618-z

Carlson J, Locke AE, Flickinger M, Zawistowski M, Levy S, Myers RM, Boehnke M, Kang HM, Scott LJ, Li JZ, et al. 2018. Extremely rare variants reveal patterns of germline mutation rate heterogeneity in humans. Nat. Commun. [Internet] 9:3753. Available from: https://www.nature.com/articles/s41467-018-05936-5

Caulin AF, Maley CC. 2011. Peto’s Paradox: evolution’s prescription for cancer prevention. Trends Ecol. Evol. [Internet] 26:175–182. Available from: https://www.sciencedirect.com/science/article/pii/S0169534711000152

Coll Macià M, Skov L, Peter BM, Schierup MH. 2021. Different historical generation intervals in human populations inferred from Neanderthal fragment lengths and mutation signatures. Nat. Commun. [Internet] 12:5317. Available from: https://www.nature.com/articles/s41467-021-25524-4

Danecek P, Auton A, Abecasis G, Albers CA, Banks E, DePristo MA, Handsaker RE, Lunter G, Marth GT, Sherry ST, et al. 2011. The variant call format and VCFtools. Bioinformatics [Internet] 27:2156–2158. Available from: https://doi.org/10.1093/bioinformatics/btr330

DeWitt WS, Harris KD, Ragsdale AP, Harris K. 2021. Nonparametric coalescent inference of mutation spectrum history and demography. Proc. Natl. Acad. Sci. [Internet] 118:e2013798118. Available from: https://www.pnas.org/doi/abs/10.1073/pnas.2013798118

DeWitt WS, Zhu L, Vollger MR, Goldberg ME, Talenti A, Beichman AC, Harris K. 2023. mutyper: assigning and summarizing mutation types for analyzing germline mutation spectra. J. Open Source Softw. [Internet] 8:5227. Available from: https://joss.theoj.org/papers/10.21105/joss.05227

Dixon P. 2003. VEGAN, a package of R functions for community ecology. J. Veg. Sci. [Internet] 14:927–930. Available from: https://onlinelibrary.wiley.com/doi/abs/10.1111/j.1654-1103.2003.tb02228.x

Dumont BL. 2019. Significant Strain Variation in the Mutation Spectra of Inbred Laboratory Mice. Mol. Biol. Evol. [Internet] 36:865–874. Available from: https://doi.org/10.1093/molbev/msz026

Duret L, Galtier N. 2009. Biased Gene Conversion and the Evolution of Mammalian Genomic Landscapes. Annu. Rev. Genomics Hum. Genet. [Internet] 10:285–311. Available from: https://doi.org/10.1146/annurev-genom-082908-150001

Egozcue JJ, Pawlowsky-Glahn V, Mateu-Figueras G, Barceló-Vidal C. 2003. Isometric Logratio Transformations for Compositional Data Analysis. Math. Geol. [Internet] 35:279–300. Available from: https://doi.org/10.1023/A:1023818214614

Gao Z, Moorjani P, Sasani TA, Pedersen BS, Quinlan AR, Jorde LB, Amster G, Przeworski M. 2019. Overlooked roles of DNA damage and maternal age in generating human germline mutations. Proc. Natl. Acad. Sci. [Internet] 116:9491–9500. Available from: https://www.pnas.org/doi/abs/10.1073/pnas.1901259116

Gao Z, Zhang Y, Cramer N, Przeworski M, Moorjani P. 2023. Limited role of generation time changes in driving the evolution of the mutation spectrum in humans. Messer PW, Perry GH, Duret L, editors. eLife [Internet] 12:e81188. Available from: https://doi.org/10.7554/eLife.81188

Goldberg ME, Harris K. 2022. Mutational Signatures of Replication Timing and Epigenetic Modification Persist through the Global Divergence of Mutation Spectra across the Great Ape Phylogeny. Genome Biol. Evol. [Internet] 14:evab104. Available from: https://doi.org/10.1093/gbe/evab104

Goldmann JM, Wong WSW, Pinelli M, Farrah T, Bodian D, Stittrich AB, Glusman G, Vissers LELM, Hoischen A, Roach JC, et al. 2016. Parent-of-origin-specific signatures of de novo mutations. Nat. Genet. [Internet] 48:935–939. Available from: https://www.nature.com/articles/ng.3597

Gori K, Baez-Ortega A. 2020. sigfit: flexible Bayesian inference of mutational signatures. :372896. Available from: https://www.biorxiv.org/content/10.1101/372896v2

Guillot G, Rousset F. 2013. Dismantling the Mantel tests. Methods Ecol. Evol. [Internet] 4:336–344. Available from: https://onlinelibrary.wiley.com/doi/abs/10.1111/2041-210x.12018

Hahn M, Peña-Garcia Y, Wang RJ. 2023. The “faulty male” hypothesis: implications for evolution and disease. Available from: https://ecoevorxiv.org/repository/view/5373/

Hamidi H, Alinejad-Rokny H, Coorens T, Sanghvi R, Lindsay SJ, Rahbari R, Ebrahimi D. 2021. Signatures of Mutational Processes in Human DNA Evolution. :2021.01.09.426041. Available from: https://www.biorxiv.org/content/10.1101/2021.01.09.426041v1

Hardy OJ, Pavoine S. 2012. Assessing phylogenetic signal with measurement error: a comparison of Mantel tests, Blomberg et al.’s K, and phylogenetic distograms. Evol. Int. J. Org. Evol. 66:2614–2621.

Harmon LJ, Glor RE. 2010. Poor statistical performance of the Mantel test in phylogenetic comparative analyses. Evolution [Internet] 64:2173–2178. Available from: https://doi.org/10.1111/j.1558-5646.2010.00973.x

Harr B, Karakoc E, Neme R, Teschke M, Pfeifle C, Pezer Ž, Babiker H, Linnenbrink M, Montero I, Scavetta R, et al. 2016. Genomic resources for wild populations of the house mouse, Mus musculus and its close relative Mus spretus. Sci. Data [Internet] 3:160075. Available from: https://www.nature.com/articles/sdata201675

Harris K. 2015. Evidence for recent, population-specific evolution of the human mutation rate. Proc. Natl. Acad. Sci. [Internet] 112:3439–3444. Available from: https://www.pnas.org/doi/abs/10.1073/pnas.1418652112

Harris K, Pritchard JK. 2017. Rapid evolution of the human mutation spectrum.McVean G, editor. eLife [Internet] 6:e24284. Available from: https://doi.org/10.7554/eLife.24284

Hoeijmakers JHJ. 2001. Genome maintenance mechanisms for preventing cancer. Nature [Internet] 411:366–374. Available from: https://www.nature.com/articles/35077232

Hwang DG, Green P. 2004. Bayesian Markov chain Monte Carlo sequence analysis reveals varying neutral substitution patterns in mammalian evolution. Proc. Natl. Acad. Sci. [Internet] 101:13994–14001. Available from: https://www.pnas.org/doi/abs/10.1073/pnas.0404142101

Islam SMA, Díaz-Gay M, Wu Y, Barnes M, Vangara R, Bergstrom EN, He Y, Vella M, Wang J, Teague JW, et al. 2022. Uncovering novel mutational signatures by de novo extraction with SigProfilerExtractor. :2020.12.13.422570. Available from: https://www.biorxiv.org/content/10.1101/2020.12.13.422570v3

Ivancevic AM, Kortschak RD, Bertozzi T, Adelson DL. 2016. LINEs between Species: Evolutionary Dynamics of LINE-1 Retrotransposons across the Eukaryotic Tree of Life. Genome Biol. Evol. [Internet] 8:3301–3322. Available from: https://doi.org/10.1093/gbe/evw243

Jiang P, Ollodart AR, Sudhesh V, Herr AJ, Dunham MJ, Harris K. 2021. A modified fluctuation assay reveals a natural mutator phenotype that drives mutation spectrum variation *within Saccharomyces cerevisiae*.Nordborg M, Przeworski M, editors. eLife [Internet] 10:e68285. Available from: https://doi.org/10.7554/eLife.68285

Jones KE, Bielby J, Cardillo M, Fritz SA, O’Dell J, Orme CDL, Safi K, Sechrest W, Boakes EH, Carbone C, et al. 2009. PanTHERIA: a species-level database of life history, ecology, and geography of extant and recently extinct mammals. Ecology [Internet] 90:2648– 2648. Available from: https://onlinelibrary.wiley.com/doi/abs/10.1890/08-1494.1

Jónsson H, Sulem P, Kehr B, Kristmundsdottir S, Zink F, Hjartarson E, Hardarson MT, Hjorleifsson KE, Eggertsson HP, Gudjonsson SA, et al. 2017. Parental influence on human germline de novo mutations in 1,548 trios from Iceland. Nature [Internet] 549:519–522. Available from: https://www.nature.com/articles/nature24018

Kaplanis J, Ide B, Sanghvi R, Neville M, Danecek P, Coorens T, Prigmore E, Short P, Gallone G, McRae J, et al. 2022. Genetic and chemotherapeutic influences on germline hypermutation. Nature [Internet] 605:503–508. Available from: https://www.nature.com/articles/s41586-022-04712-2

Keightley PD, Jackson BC. 2018. Inferring the Probability of the Derived vs. the Ancestral Allelic State at a Polymorphic Site. Genetics [Internet] 209:897–906. Available from: https://doi.org/10.1534/genetics.118.301120

Kolora SRR, Owens GL, Vazquez JM, Stubbs A, Chatla K, Jainese C, Seeto K, McCrea M, Sandel MW, Vianna JA, et al. 2021. Origins and evolution of extreme life span in Pacific Ocean rockfishes. Science [Internet] 374:842–847. Available from: https://www.science.org/doi/full/10.1126/science.abg5332

Kucab JE, Zou X, Morganella S, Joel M, Nanda AS, Nagy E, Gomez C, Degasperi A, Harris R, Jackson SP, et al. 2019. A Compendium of Mutational Signatures of Environmental Agents. Cell [Internet] 177:821–836.e16. Available from: https://www.sciencedirect.com/science/article/pii/S0092867419302636

Kumar S, Suleski M, Craig JM, Kasprowicz AE, Sanderford M, Li M, Stecher G, Hedges SB. 2022. TimeTree 5: An Expanded Resource for Species Divergence Times. Mol. Biol. Evol. [Internet] 39:msac174. Available from: https://doi.org/10.1093/molbev/msac174

Larsen F, Gundersen G, Lopez R, Prydz H. 1992. CpG islands as gene markers in the human genome. Genomics [Internet] 13:1095–1107. Available from: https://www.sciencedirect.com/science/article/pii/088875439290024M

Legendre P, Legendre L. 2012. Numerical Ecology. Elsevier

Leigh DM, Lischer HEL, Grossen C, Keller LF. 2018. Batch effects in a multiyear sequencing study: False biological trends due to changes in read lengths. Mol. Ecol. Resour. [Internet] 18:778–788. Available from: https://onlinelibrary.wiley.com/doi/abs/10.1111/1755-0998.12779

Lindahl T, Wood RD. 1999. Quality Control by DNA Repair. Science [Internet] 286:1897–1905. Available from: https://www.science.org/doi/full/10.1126/science.286.5446.1897

Lindsay SJ, Rahbari R, Kaplanis J, Keane T, Hurles ME. 2019. Similarities and differences in patterns of germline mutation between mice and humans. Nat. Commun. [Internet] 10:4053. Available from: https://www.nature.com/articles/s41467-019-12023-w

Liu Z, Samee MAH. 2021. Mutation Rate Variations in the Human Genome are Encoded in DNA Shape. :2021.01.15.426837. Available from: https://www.biorxiv.org/content/10.1101/2021.01.15.426837v2

Lynch M. 2010. Evolution of the mutation rate. Trends Genet. [Internet] 26:345–352. Available from: http://www.sciencedirect.com/science/article/pii/S0168952510001034

Lynch M, Ackerman MS, Gout J-F, Long H, Sung W, Thomas WK, Foster PL. 2016. Genetic drift, selection and the evolution of the mutation rate. Nat. Rev. Genet. [Internet] 17:704– 714. Available from: https://www.nature.com/articles/nrg.2016.104

Mantel N. 1967. The Detection of Disease Clustering and a Generalized Regression Approach. Cancer Res. 27:209–220.

Martin AP, Palumbi SR. 1993. Body size, metabolic rate, generation time, and the molecular clock. Proc. Natl. Acad. Sci. [Internet] 90:4087–4091. Available from: https://www.pnas.org/doi/abs/10.1073/pnas.90.9.4087

Martincorena I, Roshan A, Gerstung M, Ellis P, Van Loo P, McLaren S, Wedge DC, Fullam A, Alexandrov LB, Tubio JM, et al. 2015. High burden and pervasive positive selection of somatic mutations in normal human skin. Science [Internet] 348:880–886. Available from: https://www.science.org/doi/abs/10.1126/science.aaa6806

Mathieson I, Reich D. 2017. Differences in the rare variant spectrum among human populations. PLOS Genet. [Internet] 13:e1006581. Available from: https://journals.plos.org/plosgenetics/article?id=10.1371/journal.pgen.1006581

Melo D, Garcia G, Hubbe A, Assis AP, Marroig G. 2016. EvolQG - An R package for evolutionary quantitative genetics. F1000Research [Internet] 4:925. Available from: https://f1000research.com/articles/4-925/v3

Mooney JA, Marsden CD, Yohannes A, Wayne RK, Lohmueller KE. 2023. Long-term Small Population Size, Deleterious Variation, and Altitude Adaptation in the Ethiopian Wolf, a Severely Endangered Canid. Mol. Biol. Evol. [Internet] 40:msac277. Available from: https://doi.org/10.1093/molbev/msac277

Moore L, Cagan A, Coorens THH, Neville MDC, Sanghvi R, Sanders MA, Oliver TRW, Leongamornlert D, Ellis P, Noorani A, et al. 2021. The mutational landscape of human somatic and germline cells. Nature [Internet] 597:381–386. Available from: https://www.nature.com/articles/s41586-021-03822-7

Moorjani P, Amorim CEG, Arndt PF, Przeworski M. 2016. Variation in the molecular clock of primates. Proc. Natl. Acad. Sci. [Internet] 113:10607–10612. Available from: https://www.pnas.org/doi/full/10.1073/pnas.1600374113

Morrill K, Hekman J, Li X, McClure J, Logan B, Goodman L, Gao M, Dong Y, Alonso M, Carmichael E, et al. 2022. Ancestry-inclusive dog genomics challenges popular breed stereotypes. Science [Internet] 376:eabk0639. Available from: https://www.science.org/doi/full/10.1126/science.abk0639

Nabholz B, Glémin S, Galtier N. 2008. Strong Variations of Mitochondrial Mutation Rate across Mammals—the Longevity Hypothesis. Mol. Biol. Evol. [Internet] 25:120–130. Available from: https://doi.org/10.1093/molbev/msm248

Narasimhan VM, Rahbari R, Scally A, Wuster A, Mason D, Xue Y, Wright J, Trembath RC, Maher ER, van Heel DA, et al. 2017. Estimating the human mutation rate from autozygous segments reveals population differences in human mutational processes. Nat. Commun. [Internet] 8:303. Available from: https://www.nature.com/articles/s41467-017-00323-y

Nigenda-Morales, Lin, Nuñez-Valencia, Kyriazis, Beichman, Robinson, Ragsdale, Urbán, Archer, Viloria-Gómora, et al. in press. The Genomic Footprint of Whaling and Isolation in Fin Whale Populations. Nat. Commun.

Nik-Zainal S, Alexandrov LB, Wedge DC, Van Loo P, Greenman CD, Raine K, Jones D, Hinton J, Marshall J, Stebbings LA, et al. 2012. Mutational Processes Molding the Genomes of 21 Breast Cancers. Cell [Internet] 149:979–993. Available from: https://www.cell.com/cell/abstract/S0092-8674(12)00528-4

Pacifici M, Santini L, Di Marco M, Baisero D, Francucci L, Marasini GG, Visconti P, Rondinini C. 2013. Generation length for mammals. Nat. Conserv. 5:89–94.

Paradis E, Schliep K. 2019. ape 5.0: an environment for modern phylogenetics and evolutionary analyses in R. Bioinformatics [Internet] 35:526–528. Available from: https://doi.org/10.1093/bioinformatics/bty633

Pearson K. 1897. Mathematical contributions to the theory of evolution.—On a form of spurious correlation which may arise when indices are used in the measurement of organs. Proc. R. Soc. Lond. [Internet] 60:489–498. Available from: https://royalsocietypublishing.org/doi/abs/10.1098/rspl.1896.0076

Prado-Martinez J, Sudmant PH, Kidd JM, Li H, Kelley JL, Lorente-Galdos B, Veeramah KR, Woerner AE, O’Connor TD, Santpere G. 2013. Great ape genetic diversity and population history. Nature 499:471–475.

Quinlan AR, Hall IM. 2010. BEDTools: a flexible suite of utilities for comparing genomic features. Bioinformatics [Internet] 26:841–842. Available from: https://doi.org/10.1093/bioinformatics/btq033

Ragsdale AP, Thornton KR. 2023. Multiple sources of uncertainty confound inference of historical human generation times. :2023.02.23.529751. Available from: https://www.biorxiv.org/content/10.1101/2023.02.23.529751v1

Rahbari R, Wuster A, Lindsay SJ, Hardwick RJ, Alexandrov LB, Al Turki S, Dominiczak A, Morris A, Porteous D, Smith B, et al. 2016. Timing, rates and spectra of human germline mutation. Nat. Genet. [Internet] 48:126–133. Available from: https://www.nature.com/articles/ng.3469

Ratnakumar A, Mousset S, Glémin S, Berglund J, Galtier N, Duret L, Webster MT. 2010. Detecting positive selection within genomes: the problem of biased gene conversion. Philos. Trans. R. Soc. B Biol. Sci. [Internet] 365:2571–2580. Available from: https://royalsocietypublishing.org/doi/abs/10.1098/rstb.2010.0007

Risch N, Reich EW, Wishnick MM, McCarthy JG. 1987. Spontaneous mutation and parental age in humans. Am. J. Hum. Genet. [Internet] 41:218–248. Available from: https://www.ncbi.nlm.nih.gov/pmc/articles/PMC1684215/

Robinson JA, Kyriazis CC, Nigenda-Morales SF, Beichman AC, Rojas-Bracho L, Robertson KM, Fontaine MC, Wayne RK, Lohmueller KE, Taylor BL, et al. 2022. The critically endangered vaquita is not doomed to extinction by inbreeding depression. Science [Internet] 376:635–639. Available from: https://www.science.org/doi/full/10.1126/science.abm1742

Robinson PS, Coorens THH, Palles C, Mitchell E, Abascal F, Olafsson S, Lee BCH, Lawson ARJ, Lee-Six H, Moore L, et al. 2021. Increased somatic mutation burdens in normal human cells due to defective DNA polymerases. Nat. Genet. [Internet] 53:1434–1442. Available from: https://www.nature.com/articles/s41588-021-00930-y

Sasani TA, Ashbrook DG, Beichman AC, Lu L, Palmer AA, Williams RW, Pritchard JK, Harris K. 2022. A natural mutator allele shapes mutation spectrum variation in mice. Nature [Internet] 605:497–502. Available from: https://www.nature.com/articles/s41586-022-04701-5

Sayres MAW, Venditti C, Pagel M, Makova KD. 2011. Do variations in substitution rates and male mutation bias correlate with life-history traits? A study of 32 mammalian genomes. *Evolution* [Internet] 65:2800–2815. Available from: https://doi.org/10.1111/j.1558-5646.2011.01337.x

Schubert M, Ermini L, Sarkissian CD, Jónsson H, Ginolhac A, Schaefer R, Martin MD, Fernández R, Kircher M, McCue M, et al. 2014. Characterization of ancient and modern genomes by SNP detection and phylogenomic and metagenomic analysis using PALEOMIX. Nat. Protoc. [Internet] 9:1056–1082. Available from: https://www.nature.com/articles/nprot.2014.063

Seplyarskiy VB, Sunyaev S. 2021. The origin of human mutation in light of genomic data. Nat. Rev. Genet. [Internet] 22:672–686. Available from: https://www.nature.com/articles/s41576-021-00376-2

Smit A, Hubley R, Green P. 2013. RepeatMasker 4.0. *Seattle WA Inst*. Syst. Biol.

Stamatakis A. 2014. RAxML version 8: a tool for phylogenetic analysis and post-analysis of large phylogenies. Bioinformatics [Internet] 30:1312–1313. Available from: https://doi.org/10.1093/bioinformatics/btu033

Stendahl AM, Sanghvi R, Peterson S, Ray K, Lima AC, Rahbari R, Conrad DF. 2023. A naturally occurring variant of MBD4 causes maternal germline hypermutation in primates. :2023.03.27.534460. Available from: https://www.biorxiv.org/content/10.1101/2023.03.27.534460v1

Sturtevant AH. 1937. Essays on Evolution. I. On the Effects of Selection on Mutation Rate. Q. Rev. Biol. [Internet] 12:464–467. Available from: https://www.journals.uchicago.edu/doi/abs/10.1086/394543

Sung W, Ackerman MS, Miller SF, Doak TG, Lynch M. 2012. Drift-barrier hypothesis and mutation-rate evolution. Proc. Natl. Acad. Sci. [Internet] 109:18488–18492. Available from: https://www.pnas.org/doi/abs/10.1073/pnas.1216223109

Tang Y, Horikoshi M, Li W. 2016.ggfortify: unified interface to visualize statistical results of popular R packages. R J 8:474.

Tate JG, Bamford S, Jubb HC, Sondka Z, Beare DM, Bindal N, Boutselakis H, Cole CG, Creatore C, Dawson E, et al. 2019. COSMIC: the Catalogue Of Somatic Mutations In Cancer. Nucleic Acids Res. [Internet] 47:D941–D947. Available from: https://doi.org/10.1093/nar/gky1015

Taub MA, Corrada Bravo H, Irizarry RA. 2010. Overcoming bias and systematic errors in next generation sequencing data. Genome Med. [Internet] 2:87. Available from: https://doi.org/10.1186/gm208

Thomas GWC, Wang RJ, Puri A, Harris RA, Raveendran M, Hughes DST, Murali SC, Williams LE, Doddapaneni H, Muzny DM, et al. 2018. Reproductive Longevity Predicts Mutation Rates in Primates. Curr. Biol. [Internet] 28:3193–3197.e5. Available from: https://www.sciencedirect.com/science/article/pii/S0960982218311345

Tom JA, Reeder J, Forrest WF, Graham RR, Hunkapiller J, Behrens TW, Bhangale TR. 2017. Identifying and mitigating batch effects in whole genome sequencing data. BMC Bioinformatics [Internet] 18:351. Available from: https://doi.org/10.1186/s12859-017-1756-z

Upham NS, Esselstyn JA, Jetz W. 2019. Inferring the mammal tree: Species-level sets of phylogenies for questions in ecology, evolution, and conservation. PLOS Biol. [Internet] 17:e3000494. Available from: https://journals.plos.org/plosbiology/article?id=10.1371/journal.pbio.3000494

Van der Auwera GA, Carneiro MO, Hartl C, Poplin R, Del Angel G, Levy-Moonshine A, Jordan T, Shakir K, Roazen D, Thibault J, et al. 2013. From FastQ data to high confidence variant calls: the Genome Analysis Toolkit best practices pipeline. Curr. Protoc. Bioinforma. 43:11.10.1-33.

Vazquez JM, Lynch VJ. 2021. Pervasive duplication of tumor suppressors in Afrotherians during the evolution of large bodies and reduced cancer risk.Rokas A, Wittkopp PJ, Stearns SC, Gorbunova V, editors. eLife [Internet] 10:e65041. Available from: https://doi.org/10.7554/eLife.65041

Vollger MR, Dishuck PC, Harvey WT, DeWitt WS, Guitart X, Goldberg ME, Rozanski AN, Lucas J, Asri M, Munson KM, et al. 2023. Increased mutation and gene conversion within human segmental duplications. Nature [Internet] 617:325–334. Available from: https://www.nature.com/articles/s41586-023-05895-y

Wang RJ, Al-Saffar SI, Rogers J, Hahn MW. 2023. Human generation times across the past 250,000 years. Sci. Adv. [Internet] 9:eabm7047. Available from: https://www.science.org/doi/full/10.1126/sciadv.abm7047

Wang RJ, Raveendran M, Harris RA, Murphy WJ, Lyons LA, Rogers J, Hahn MW. 2022. De novo Mutations in Domestic Cat are Consistent with an Effect of Reproductive Longevity on Both the Rate and Spectrum of Mutations. Mol. Biol. Evol. [Internet] 39:msac147. Available from: https://doi.org/10.1093/molbev/msac147

Wong WSW, Solomon BD, Bodian DL, Kothiyal P, Eley G, Huddleston KC, Baker R, Thach DC, Iyer RK, Vockley JG, et al. 2016. New observations on maternal age effect on germline de novo mutations. Nat. Commun. [Internet] 7:10486. Available from: https://www.nature.com/articles/ncomms10486

Wu FL, Strand AI, Cox LA, Ober C, Wall JD, Moorjani P, Przeworski M. 2020. A comparison of humans and baboons suggests germline mutation rates do not track cell divisions. PLOS Biol. [Internet] 18:e3000838. Available from: https://journals.plos.org/plosbiology/article?id=10.1371/journal.pbio.3000838

Zhu YO, Siegal ML, Hall DW, Petrov DA. 2014. Precise estimates of mutation rate and spectrum in yeast. Proc. Natl. Acad. Sci. [Internet] 111:E2310–E2318. Available from: https://www.pnas.org/doi/abs/10.1073/pnas.1323011111

