## Supplementary notes and figures for "“Evolution of the mutation spectrum across a mammalian phylogeny”"

#### Note S1. Additional information about 5-mer and 7-mer spectrum results.

As in our analyses of the phylogenetic signal of 1-mer and 3-mer spectra, we obtained qualitatively consistent phylogenetic signal results for 5-mer and 7-mer spectra (**Figure 4**) when using an ultrametric tree (**Figure S18**), stratifying spectra by biased gene conversion categories (**Figure S19**), using cosine distance (1- cosine similarity) instead of Aitchison distance to measure distance between spectra (**Figure 20**), and folding the spectrum to eliminate sensitivity to incorrect ancestral allele inference (**Figure S21**).

**$K_{mult}$ test.** Values of $K_{mult}$ for 5-mer and 7-mer spectra are significant, but considerably lower than those of 1-mer and 3-mer spectra (5-mer: $K_{mult} = 0.1$, $p < 0.001$; 7-mer: $K_{mult} = 0.07$, $p < 0.001$; 999 permutations), which may be due to weaker phylogenetic signal, or indicate that the dimensionality of the mutation spectrum is growing much faster than the number of distinct mutational signatures that make up these mutation spectra (Adams & Collyer 2019) (**Figure S10**).

**Sub-spectra, biological and technical confounders.** Differences in genetic diversity and age at first reproduction are significantly correlated with 5-mer spectrum distances (**Figure S24**). The phylogeny is also more correlated with 7-mer spectrum distances than any other confounder, but reference genome scaffold N50 is also significantly correlated with 7-mer spectrum distances (**Figure S24**). When 5-mer or 7-mer spectra are separated into sub-spectra based on central 1-mer mutation type (e.g. 256 5-mers that have A>T as the central 1-mer mutation), most sub-spectra continue to show a significant phylogenetic signal, though A>T 5-mers and A>T, A>C and C>T 7-mers do not have significant phylogenetic signal after correcting for multiple testing (**Figure S25-S26**). Reference genome scaffold N50 is more correlated than phylogenetic distance for A>C, A>T, C>A and C>T 7-mer sub-spectra (**Figure S25-S26**).

This dependence on scaffold N50 may be partially driven by the vaquita, which has the highest genome scaffold contiguity of any species (**Table 1**), but also the sparsest 7-mer mutation spectrum due to its extremely low diversity, making it have elevated 7-mer mutation spectrum distance from all other species (an outlier on PC1 in the 7-mer based PCA (**Figure S15B**) and the elevated points in the scatter plot in **Figure 4A**, right panel). When low-diversity species, including the vaquita, are excluded from the analysis as described below, scaffold N50 is *no longer* significantly correlated with the 7-mer spectrum or any of its sub-spectra, though the 7-mer spectrum remains correlated with genetic diversity and age at first reproduction (**Figure S23**).

**Data sparsity.** The decrease in phylogenetic signal with increased spectrum dimensionality may be due in part to data sparsity. We were able to somewhat mitigate this data sparsity issue by excluding two low-diversity species (vaquita and polar bear) and substituting the higher-diversity Eastern North Pacific fin whale population for the low diversity Gulf of California population. The elimination of these species results in downsampling to ~890k SNPs across species instead of ~130k (**Figure S23**; 5-mer spectrum $r = 0.86$, $p < 2.8e{-5}$; 7-mer spectrum $r = 0.74$, $p < 3.7e{-5}$), which increases the amount of phylogenetic signal we measure when looking at the 7-mer spectrum, but still does not yield a better fit than the 7-mer spectrum permuted across 5-mer categories ($p > 0.75$) (**Figure S23B**).

**Note S2. Proof that mutation counts and relative rates yield identical estimates of CLR mutation spectrum distances.**

The mutation spectrum is sometimes computed using raw mutation type counts, but it is sometimes computed using SNP counts that have been rescaled by target size (i.e. the number of A**A**A>A**C**A SNPs in the genome divided by the number of AAA 3-mers where a SNP might be called if one exists). We refer to these rescaled SNP counts as relative rates since in the absence of natural selection, they should be proportional to the rates at which different mutation types arise. When using relative rates, we have found that Euclidean distances between mutation spectra are largely determined by the abundance of high-rate CpG mutations, while low-rate mutation types, which proportionally may differ greatly between species, have negligible impact (**Figure S33**). Intuitively, mutation spectra that are computed using relative rates look slightly "spikier" than spectra computed from raw counts, with larger contributions due to hypermutable mutation types such as CpGs. In contrast, we have found that CLR-transformed mutation spectrum distance depends more strongly on differences in the abundance on rarer mutation types, and we can show that the value of this distance does not actually depend on whether SNP counts are normalized for genome content or not, as long as all species have been corrected to have the same genome content.

To prove this, we represent the human-genome-content rescaled mutation count of a *k*-mer $m$ mutating to *k*-mer $j$ in species A as ${x^{(r)}}_{m \to j,A}$ ($r$ designates that the count has been rescaled by human genome target content, as in the **Methods**).

A full vector of all rescaled mutation counts for species A is $x_A^{(r)}$.

The target size of *k*-mer $m$ in the human genome (which all species' counts have been rescaled relative to) is $t_{m,h}$, and the vector of target sizes for all mutation types is $t_h$ (note that since each *k*-mer can mutate to three possible mutation types, the length of the vector $t_h$ is three times longer than the vector just of targets themselves, with each target repeated 3x).

The Euclidean distance between the raw SNP vectors $x_A^{(r)}$ and $x_B^{(r)}$ will generally be different from the Euclidean distance between the normalized vectors $x_A^{(r)}/t_h$ and $x_B^{(r)}/t_h$.

However, we will show that the Aitchison distances between these two different ways of scaling the mutation spectra are identical, provided we have rescaled our mutation counts to account for any differences between the target sizes of the genomes from which the two different mutation spectra were sampled (in our study, all counts are rescaled to reflect human reference genome composition; see **Methods**).

CLR transformation involves dividing a value by the geometric mean of the full compositional vector, and taking the natural log:

$$CLR_I = ln\,(\frac{x_I}{GM(x)})$$

where $x_I$ is a value of a compositional vector, and $x$ is the full compositional vector, and $GM$ is the geometric mean function.

The geometric mean (GM) has a very useful property:

$$GM\left(\frac{X}{Y}\right) = \frac{GM(X)}{GM(Y)}$$

The Aitchison distance between CLR-transformed vectors $x_A$ and $x_B$ is defined as:

$$d(x_A, x_B) = \sqrt{\sum_{i=1}^{S}\left[ln\left(\frac{x_{i,A}}{GM(x_A)}\right) - ln\left(\frac{x_{i,B}}{GM(x_B)}\right)\right]^2}$$

where S is the length of each vector, $x_{i,A}$ is the i[th] value of vector $x_A$ and $x_{i,B}$ is the i[th] value of vector $x_B$.

Due to the logarithmic property that ln(X)-ln(Y) = ln(X/Y), this equation can also be written as:

$$d(x_A, x_B) = \sqrt{\sum_{i=1}^{S}\left[ln\left(\frac{\frac{x_{i,A}}{GM(x_A)}}{\frac{x_{i,B}}{GM(x_B)}}\right)\right]^2}$$

In the above equation, we can define the mutation spectrum vectors $x_A$ and $x_B$ to either be unnormalized count vectors $x_A^{(r)}$ or to be normalized vectors of the form $x_A^{(r)}/t_h$.

If we use the unnormalized mutation count vector $x_A^{(r)}$ to calculate Aitchison distance, then the CLR value for the count (denoted $CLR_C$ for 'count') of mutation type $m \to j$ in species A is:

$$CLR_{C,m\to j,A} = ln\left(\frac{{x^{(r)}}_{m\to j,A}}{GM(x_A^{(r)})}\right)$$

However, if we use the target-normalized mutation count vector

$$x_A^{(r)}/t_h\ ,$$

the CLR value for the mutation count for mutation type $m \to j$ divided by the target size of $m$ (denoted $CLR_R$ for 'rate') in species A is:

$$CLR_{R,m\to j,A} = ln\left(\frac{\frac{{x^{(r)}}_{m\to j,A}}{t_{m,h}}}{GM\left(\frac{x_A^{(r)}}{t_h}\right)}\right)$$

We can then separate out the numerator and denominator of the geometric mean to obtain:

$$CLR_{R,m\to j,A} = ln\left(\frac{\frac{{x^{(r)}}_{m\to j,A}}{t_{m,h}}}{\frac{GM\left(x_A^{(r)}\right)}{GM(t_h)}}\right)$$

Some algebraic rearrangement yields:

$$CLR_{R,m\to j,A} = ln\left(\frac{{x^{(r)}}_{m\to j,A} * GM(t_h)}{t_{m,h} * GM\left(x_A^{(r)}\right)}\right)$$

Similarly, for species B:

$$CLR_{R,m\to j,B} = ln\left(\frac{{x^{(r)}}_{m\to j,B} * GM(t_h)}{t_{m,h} * GM\left(x_B^{(r)}\right)}\right)$$

Note that the target sizes $t_{m,h}$ are identical for species A and B, because both species have had their spectra rescaled to match the human genomic target content.

To calculate the Aitchison distance, we compute the difference between the two CLR values and observe that the target sizes cancel out as follows:

$$ln\left(\frac{x^{(r)}{}_{m\to j,A} * GM(t_h)}{t_{m,h} * GM\left(x_A^{(r)}\right)}\right) - ln\left(\frac{x^{(r)}{}_{m\to j,B} * GM(t_h)}{t_{m,h} * GM\left(x_B^{(r)}\right)}\right) = ln\left(\frac{\frac{x^{(r)}{}_{m\to j,A} * GM(t_h)}{t_{m,h} * GM\left(x_A^{(r)}\right)}}{\frac{x^{(r)}{}_{m\to j,B} * GM(t_h)}{t_{m,h} * GM\left(x_B^{(r)}\right)}}\right)$$

This simplifies to:

$$ln\left(\frac{x^{(r)}{}_{m\to j,A} * \cancel{GM(t_h)} * \cancel{t_{m,h}} * GM\left(x_B^{(r)}\right)}{\cancel{t_{m,h}} * GM\left(x_A^{(r)}\right) * x^{(r)}{}_{m\to j,B} * \cancel{GM(t_h)}}\right)$$

All information regarding genomic target size cancels out, leaving:

$$ln\left(\frac{x^{(r)}{}_{m\to j,A}}{GM\left(x_A^{(r)}\right)} * \frac{GM\left(x_B^{(r)}\right)}{x^{(r)}{}_{m\to j,B}}\right) = ln\left(\frac{\frac{x^{(r)}{}_{m\to j,A}}{GM\left(x_A^{(r)}\right)}}{\frac{\vec{x}_{m\to j,B}}{GM(\vec{x}_B)}}\right) = ln\left(\frac{x^{(r)}{}_{m\to j,A}}{GM\left(x_A^{(r)}\right)}\right) - ln\left(\frac{x^{(r)}{}_{m\to j,B}}{GM\left(x_B^{(r)}\right)}\right)$$

This is simply the difference between the count-based CLR values for species A and B.

A similar argument proves that a CLR distance computed from raw mutation counts is identical to the CLR distance computed from mutation type proportions. We reiterate that proportions should only be used as long as target sizes have been rescaled to the same genomic content – otherwise proportions, rates, or counts will reflect species-specific differences in genome content.

### Supplementary Figures

**Figure S1. Time-scaled phylogenetic tree**. Ultrametric tree in which branch lengths represent millions of years before present, from TimeTree (Kumar et al. 2022).

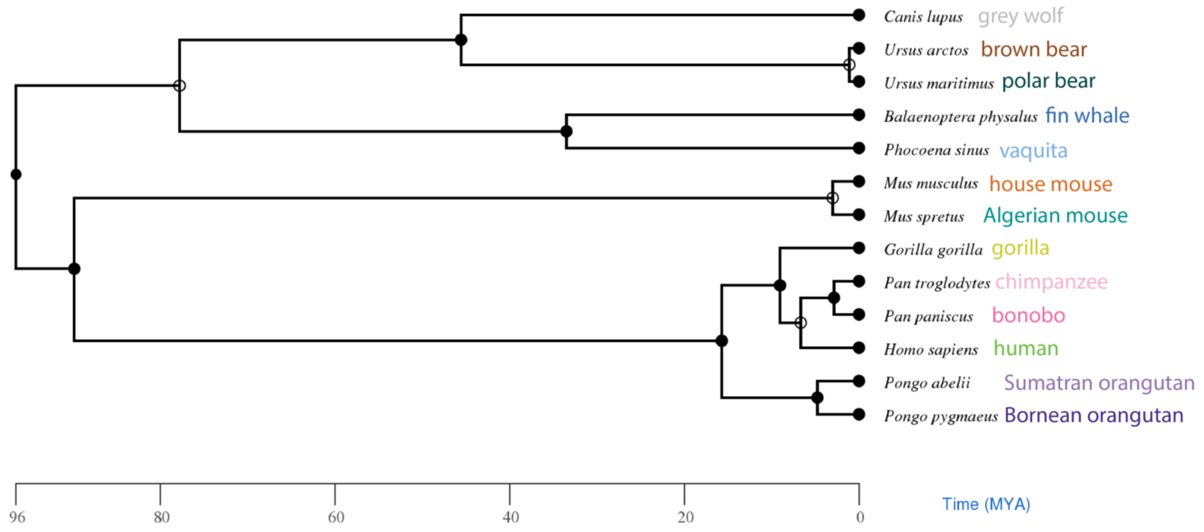

**Figure S2. Additional principal components.** Principal component analyses based on the 1-mer and 3-mer mutation spectra. Each point represents a single individual's mutation spectrum. Here, we plot additional PCs to show alternate clustering of points when the third principal component (PC3) is included.

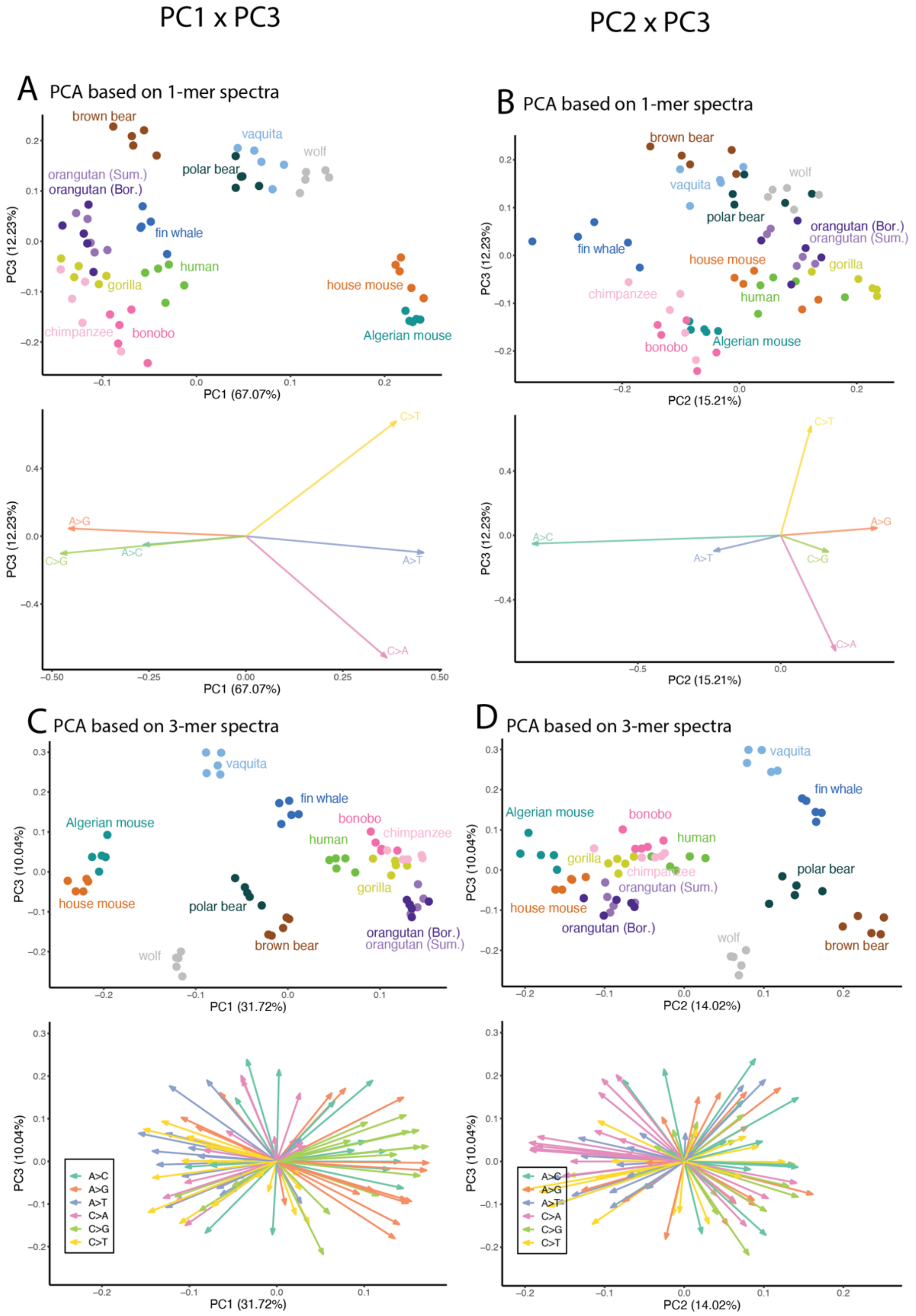

**Figure S3. PCA of isometric log-ratio (ILR) transformed mutation spectra resembles PCA of CLR transformed mutation spectra.** By some metrics, ILR is a potentially more robust compositional transformation than the CLR (Egozcue et al. 2003). We find that our results are qualitatively similar regardless of whether the ILR or CLR is used to calculate distances.

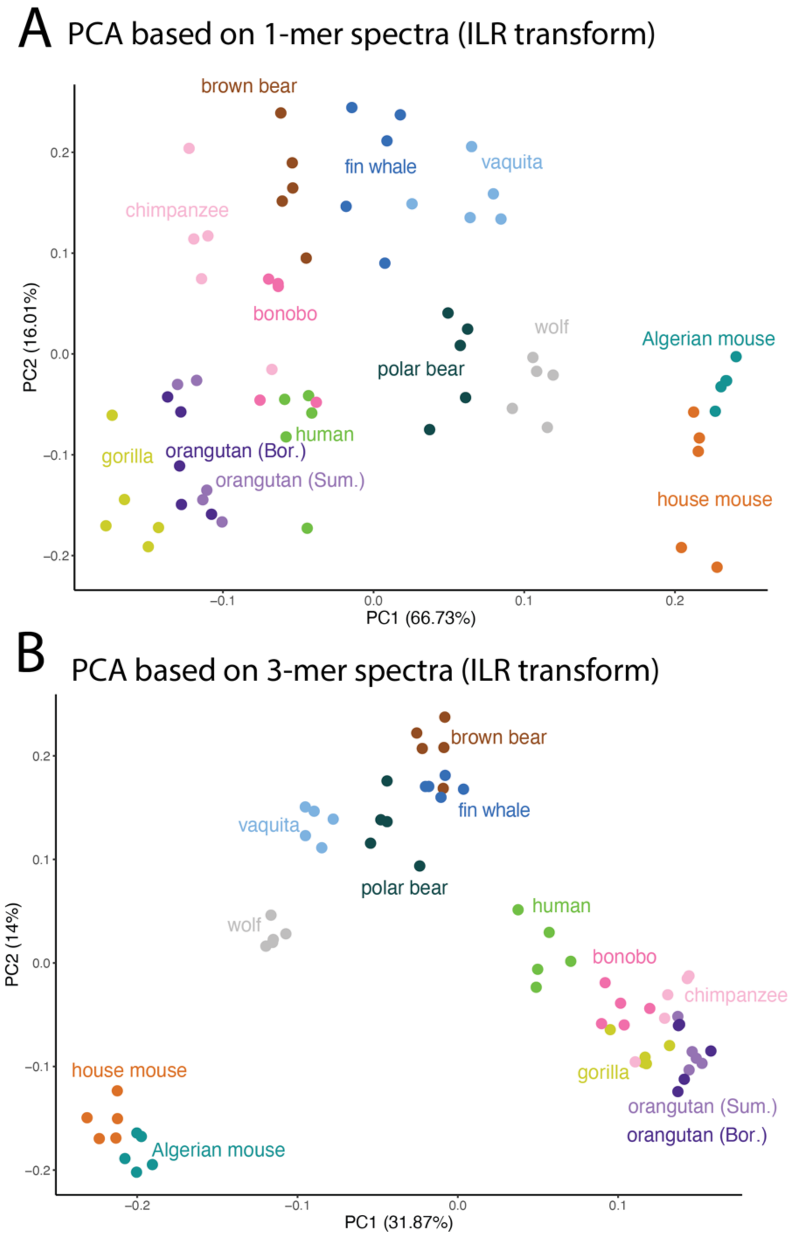

**Figure S4. Phylogenetic signal plots with labels.** These plots are identical to the distance plots shown in main text **Figure 3A** but with every point labeled with the specific species comparisons for ease of reference.

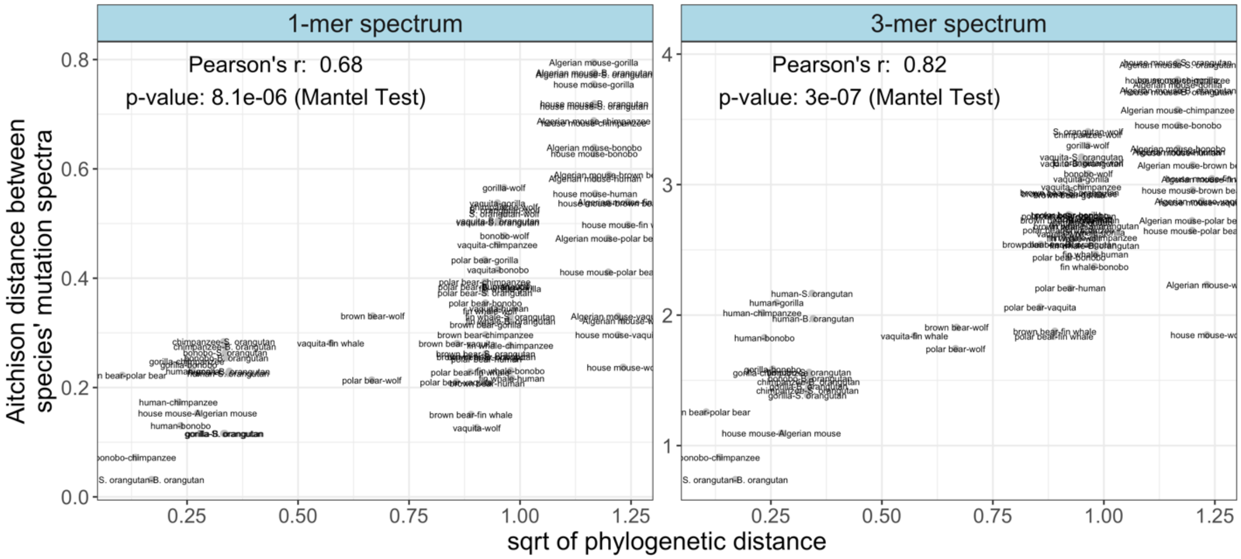

**Figure S5. Phylogenetic signal analyses based on an ultrametric timetree are consistent with distances based on the genetic alignment *RAxML* tree.** Distance plots with phylogenetic distance based on shared branch lengths from the ultrametric time tree (tree in **Figure S1**). Results are qualitatively similar to results based on the tree in which branch lengths represent expected substitutions per site (**Figure 3A**). *p*-values based on the Mantel test with 9,999,999 permutations.

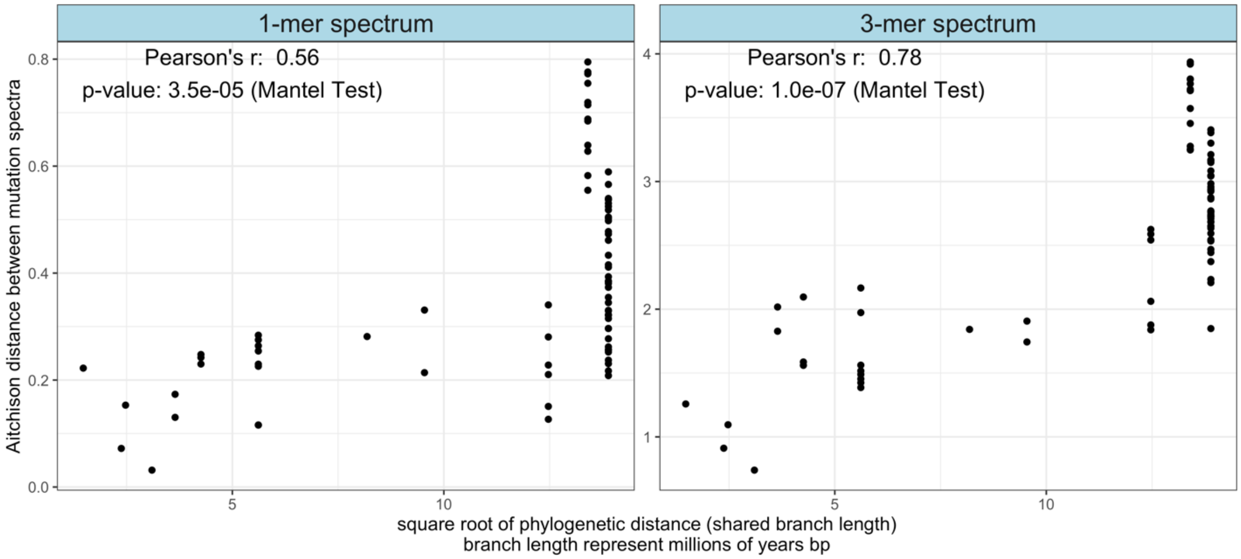

**Figure S6. Cosine distance is correlated with phylogenetic distance.** Plots showing the correlation between cosine distance (1-cosine similarity) and the square root of phylogenetic distance. *p*-values from the Mantel test with 9,999,999 permutations.

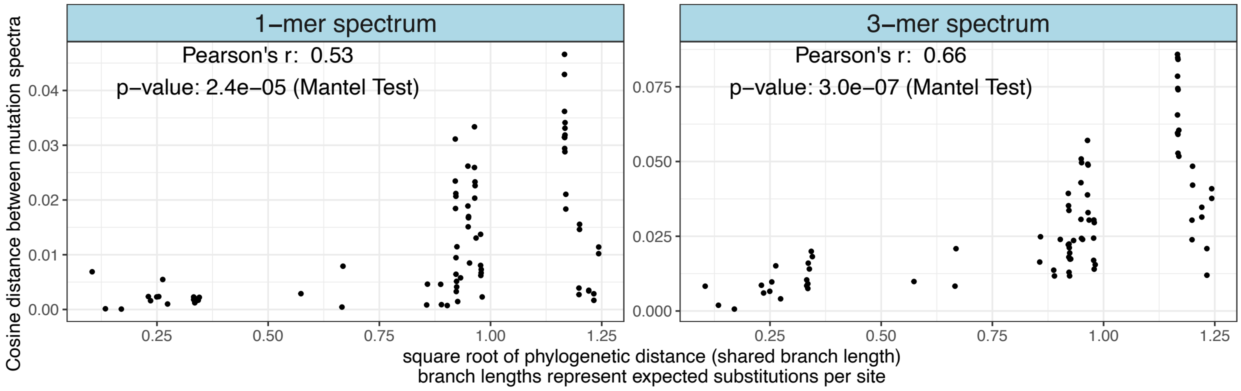


**Figure S7. Phylogenetic signal results are robust to using the isometric log-ratio transform (ILR) instead of the centered log ratio transform.** When carrying out Aitchison transformations, a potentially more robust transformation is the ILR. We find that our results are consistent whether the ILR or CLR is used to calculate distances. *p*-values based on the Mantel test with 9,999,999 permutations.

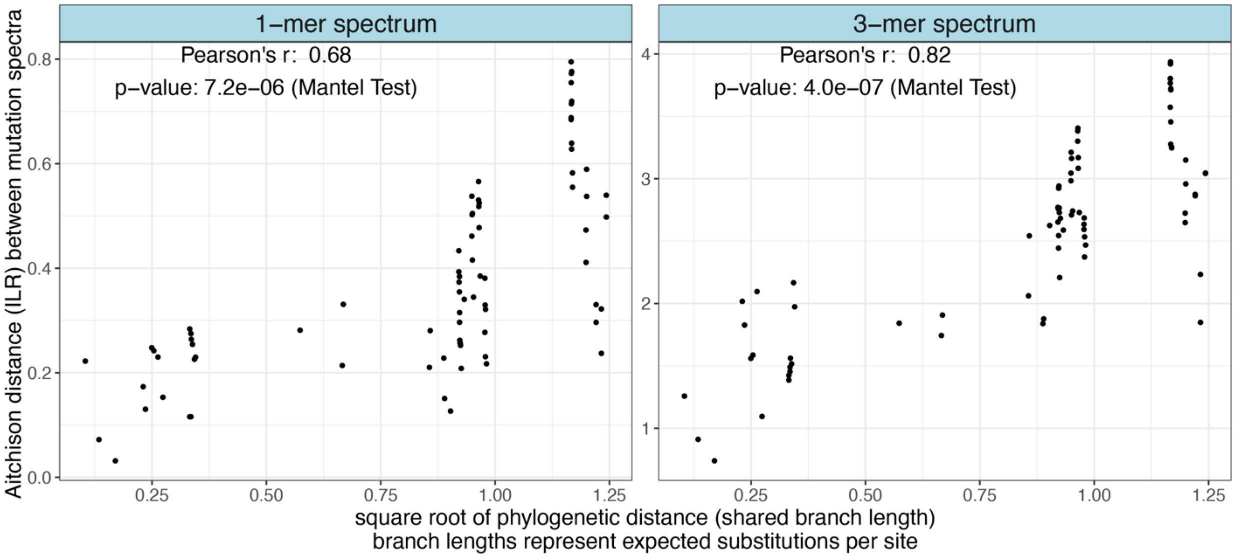

**Figure S8. Phylogenetic signal results are robust to mispolarization.** We measured the phylogenetic signal of a 'folded' mutation spectrum in which reverse mutation types are collapsed into the same category to determine whether the phylogenetic signal of the unfolded spectrum could be driven by mispolarization of ancestral alleles. For example, A**C**G>A**A**G and A**A**G>A**C**G are considered equivalent after folding. Despite the reduction in power caused by analyzing fewer overall mutation types, our findings of significant phylogenetic signal persist, indicating that they are robust to mispolarization of ancestral mutation types. *p*-values based on the Mantel test with 9,999,999 permutations.

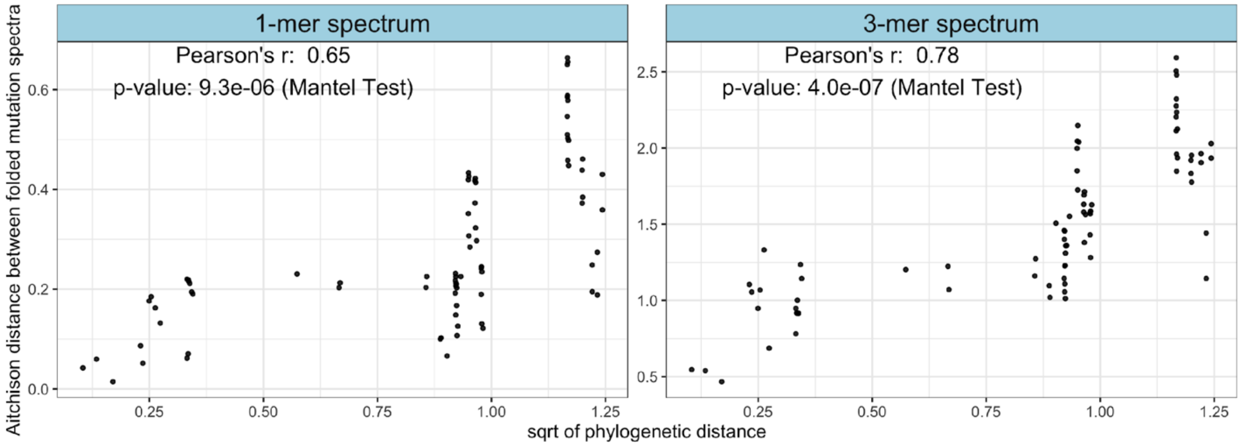

**Figure S9. 1-mer spectrum results are similar when CpG>TpG mutations are partitioned into a seventh mutation category, or when they are entirely excluded. A)** In main text 1-mer results, CpG>TpG mutations are included in the overall C>T mutation count. Here, they are instead separated into a 7[th] mutation type, yielding an augmented 1-mer mutation spectrum that is often used for analysis of de novo mutations (A>C,A>G, A>T, C>A, C>G, nonCpG C>T, CpG>TpG). The phylogenetic signal of this 1mer+CpG spectra remain significant ($r = 0.59$, $p < 3.1e\text{-}4$ with CpG>TpG as own category, compared to $r = 0.68$, $p < 8e\text{-}6$ when CpG>TpG are included as part of the C>T 1-mer category in **Figure 3**). **B)** The confounders correlated with the mutation spectrum when CpG>TpG mutations are separated out: all remain weaker than the phylogenetic signal, but genetic diversity is approaching the r-value of the phylogenetic signal. **C)** In this spectrum, CpG>TpG mutations are removed entirely from the spectrum, but the phylogenetic signal remains significant. **D)** As in (B), genetic diversity is approaching the *r*-value of phylogenetic signal.

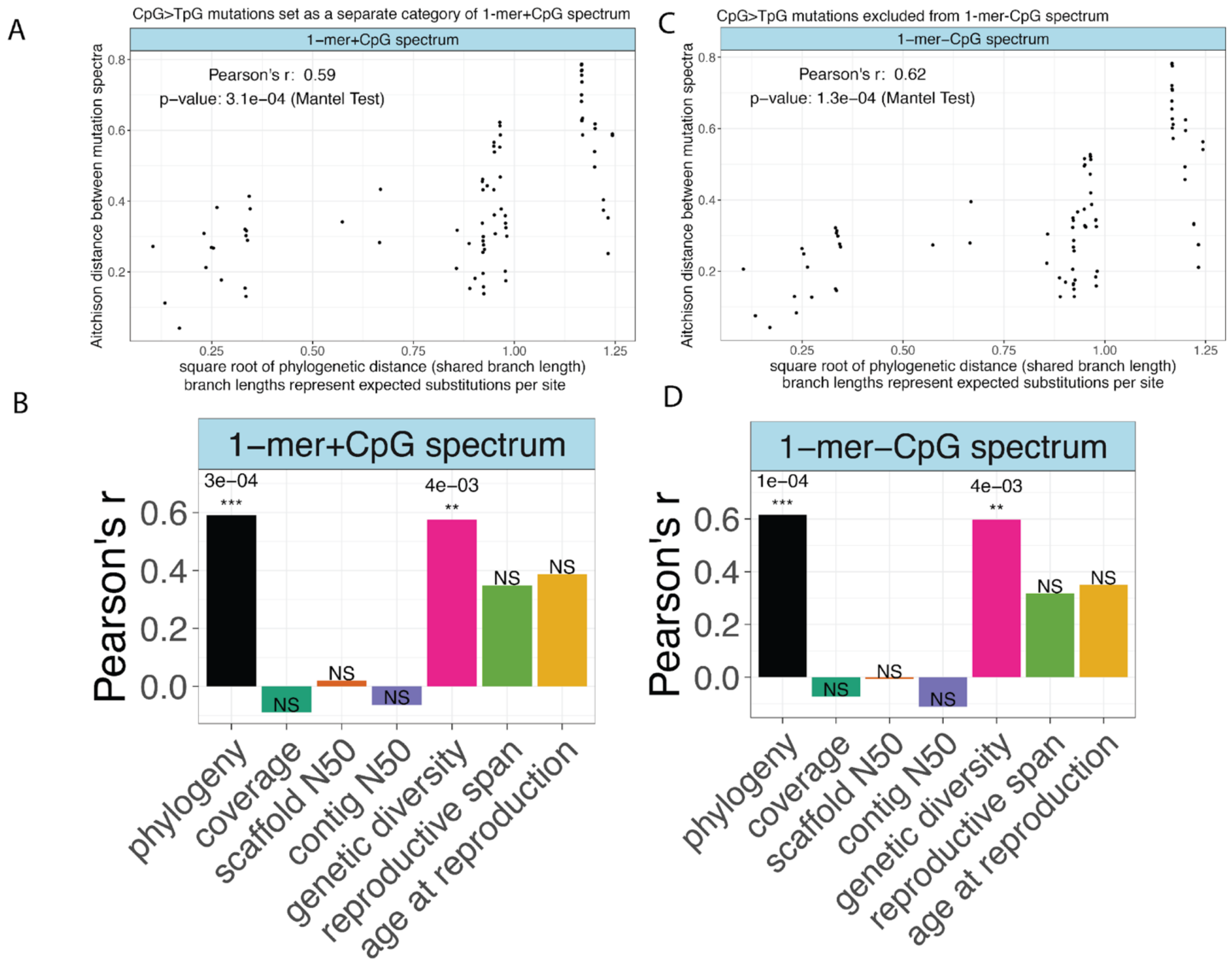

**Figure S10. $K_{mult}$ test.** Values of the $K_{mult}$ statistic, a multivariate version of Blomberg's $K$ (Adams 2014). Each mutation spectrum has a significant value of $K_{mult}$ (based on a permutation test with 999 permutations), but the value of $K_{mult}$ decreases with increased dimensionality, which may be due to lower fractions of components having phylogenetic signal in the higher-dimensional spectra.

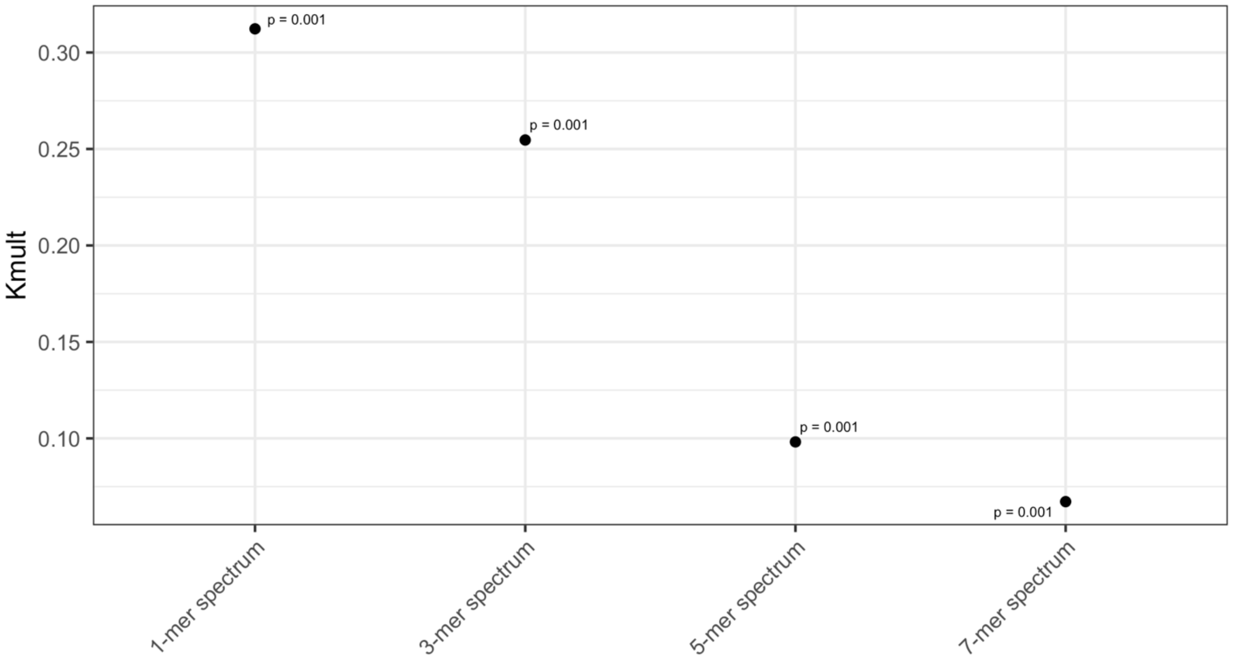

**Figure S11. Biased gene conversion is not the sole driver of mutation spectrum phylogenetic signal.** 3-mer spectrum distances were calculated based on 3-mers that were separated into GC-biased gene conversion (BGC) categories based on their mutating central basepair: BGC-conserved mutations (BGC_conserved), consisting of A>T and C>G mutations which are not affected by biased gene conversion; strong-to-weak mutations (BGC_SW), consisting of C>A and C>T mutations which are disfavored by BGC; and weak-to-strong (BGC_WS) mutations, consisting of A>C and A>G mutations which are favored by BGC. Importantly, the correlation between 3-mer spectrum distances and phylogenetic distance are highly significant in the BGC_conserved category, indicating that BGC is not the driver of the phylogenetic signal we observe. *p*-values based on the Mantel test with 9,999,999 permutations.

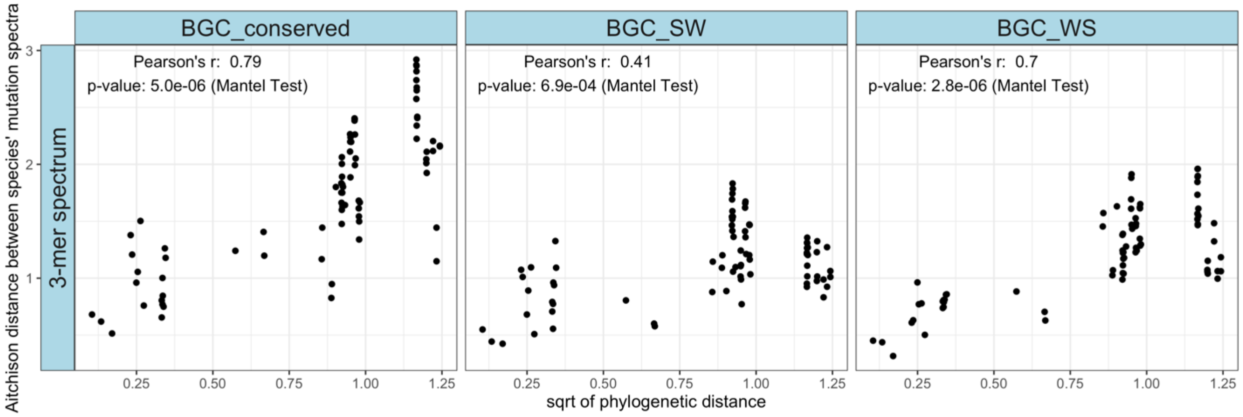

**Figure S12. Phylogenetic signal of technical confounders.** To determine whether any technical confounders could be contributing to our observed phylogenetic signal in the mutation spectrum, we used the Mantel test to determine whether differences between species' sequence coverage (a measure of dataset quality), contig N50, or scaffold N50 of the reference genomes (measures of genome assembly quality) had a strong phylogenetic signal. Coverage and scaffold N50 did not have a significant phylogenetic signal. Contig N50 showed a significant phylogenetic signal of smaller magnitude than what we observed for the phylogenetic signal of mutation spectra ($r = 0.42$ and $p < 7e\text{-}3$ for contig N50 compared to $r = 0.68$ and 0.82, and $p$-values < 8e-6 and 3e-7 for the 1- and 3-mer mutation spectra, respectively in **Figure 3A**). $p$-values based on the Mantel test with 99,999 permutations (note that fewer permutations can be used for the Mantel test here, since no test is hitting the minimum $p$-value for 99,999 permutations (1e-5)).

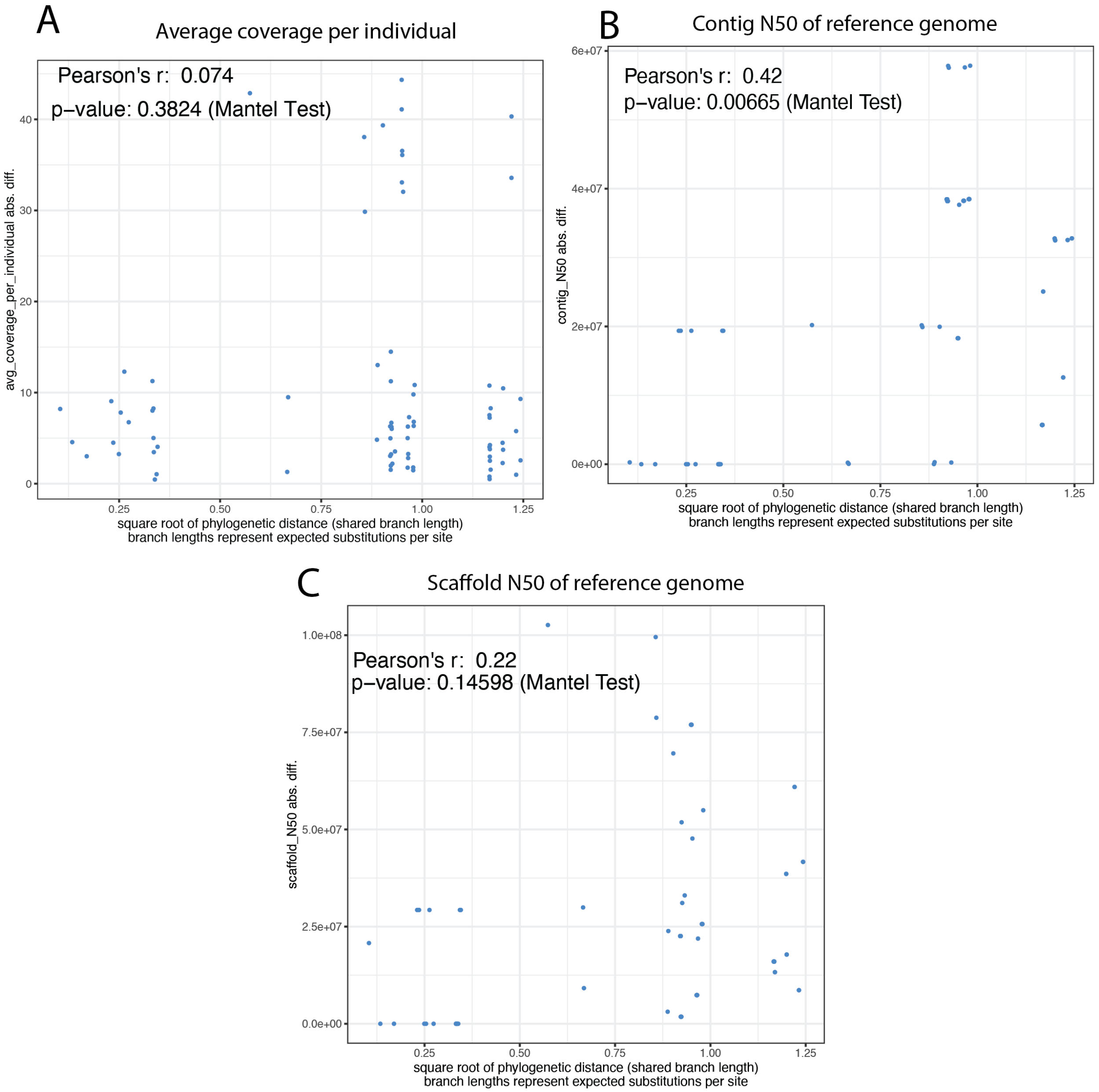

**Figure S13. Confounder correlation results are qualitatively similar when using a phylogenetically-aware Mantel test.** A version of the Mantel test called the phylogenetic permutation (PP) Mantel test can be used to test for significant correlations between variables which may share a common phylogenetic signal that could cause falsely significant correlations if not corrected for. The test permutes species that are closely related in the phylogenetic tree with higher probability to generate a null set of permutations that incorporate any shared phylogenetic signal. The results are largely consistent across confounders between the uncorrected Mantel test and the phylogenetically-aware Mantel test ("phyloMantel"), though genetic diversity and age at reproduction become slightly *more* significantly correlated with the 1-mer spectrum when using phyloMantel. $p$-values based on the Mantel test with 99,999 permutations (note that fewer permutations can be used for the Mantel test here, since no test is hitting the minimum $p$-value for 99,999 permutations (1e-5)).

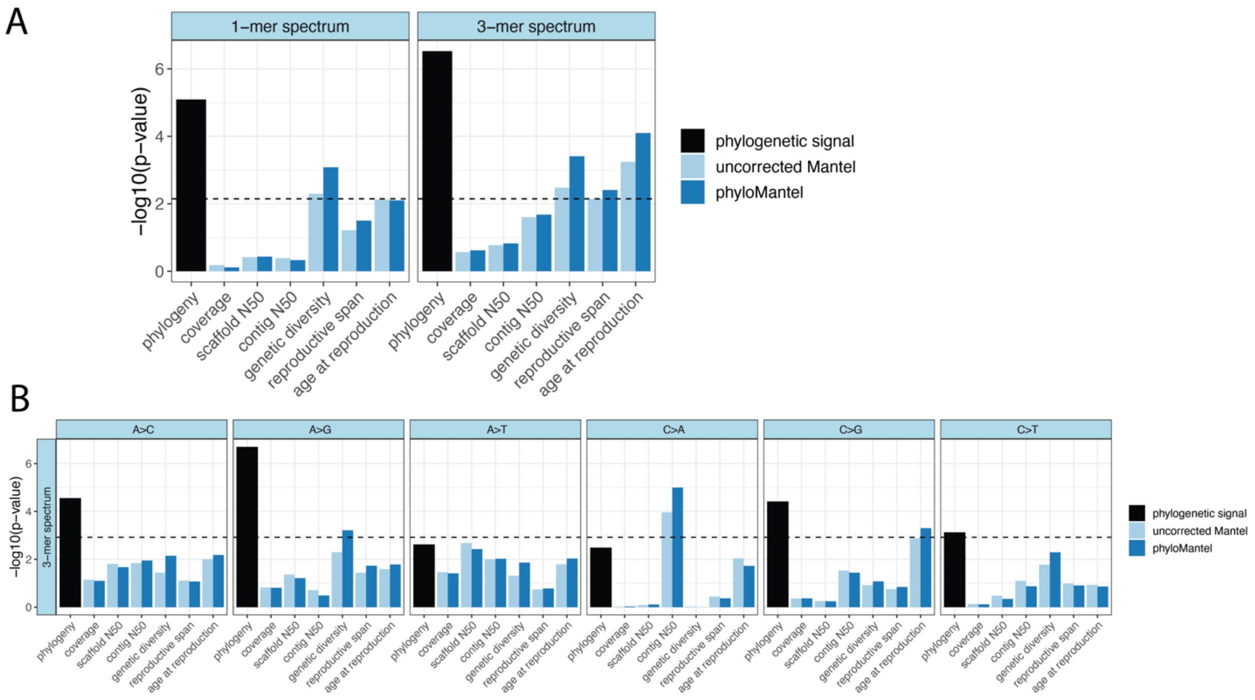

**Figure S14. Phylogenetic signal of biological confounders.** To determine whether biological confounders could be contributing to our observed phylogenetic signal in the mutation spectrum, we used the Mantel test to determine whether differences between **(A)** species' genetic diversity (measured as Watterson's $\theta$), **(B)** age at first reproduction, or **(C)** reproductive span had a strong phylogenetic signal. Both genetic diversity ($r$ =0.6, $p < 0.0013$) and reproductive lifespan ($r = 0.4$, $p < 0.005$) had a significant phylogenetic signal, though not as strong as the phylogenetic signal of the mutation spectrum ($r = 0.68$ and 0.82, and $p$-values < 8e-6 and 3e-7 for the 1- and 3-mer mutation spectra, respectively in **Figure 3A**). $p$-values based on the Mantel test with 99,999 permutations (note that fewer permutations can be used for the Mantel test here, since no test is hitting the minimum $p$-value for 99,999 permutations (1e-5)).

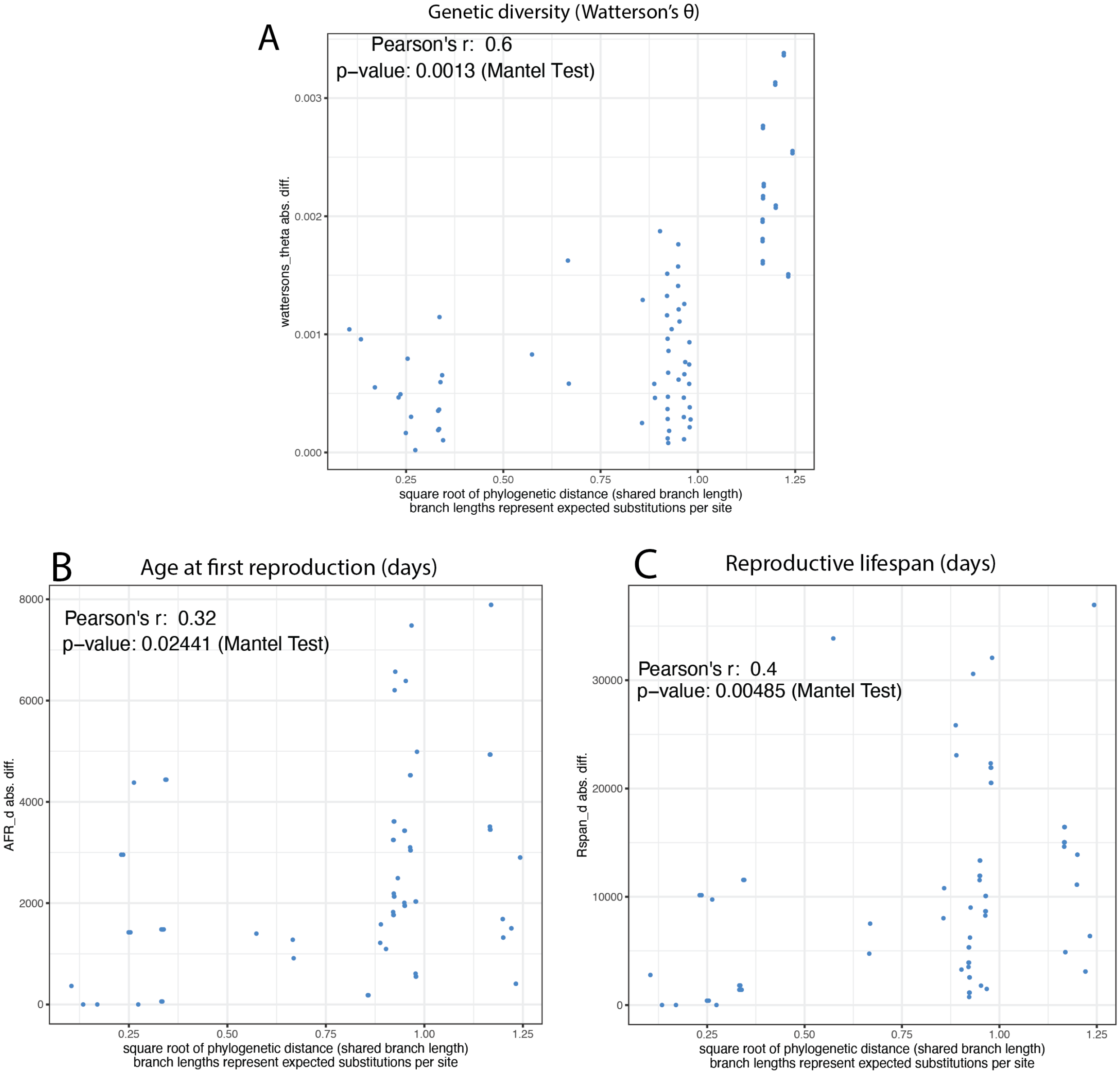

**Figure S15. Extended sequence context PCAs explain less variance than 1-mer and 3-mer PCAs but yield more visually distinct cladistic groupings.** As in Figure **2C-D**, plots of PCA based on individuals' mutation spectra. **A)** PCA based on the 5-mer spectrum. **B)** PCA based on the 7-mer spectrum. 5-mer and 7-mer PCA plots including PC3 are in **Figure S16**. **C)** Distributions of cosine similarities between 5-mer and 7-mer mutation spectra for every pair of species in our dataset. Horizontal lines denote the median.

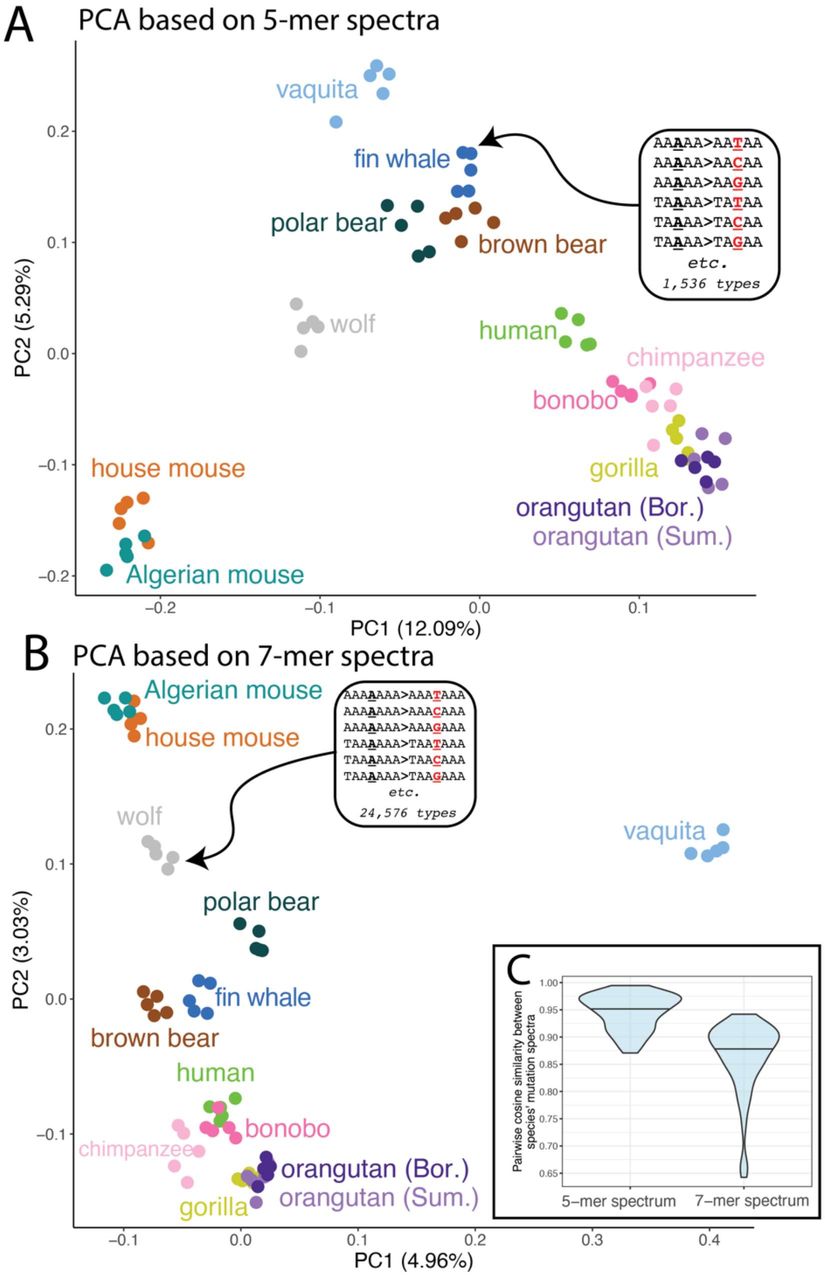

**Figure S16. Additional principal components.** Principal component analyses based on the 1-mer and 3-mer mutation spectra. Each point represents a single individual's mutation spectrum. Here, we plot additional PCs to show alternate clustering of points when the third principal component (PC3) is included.

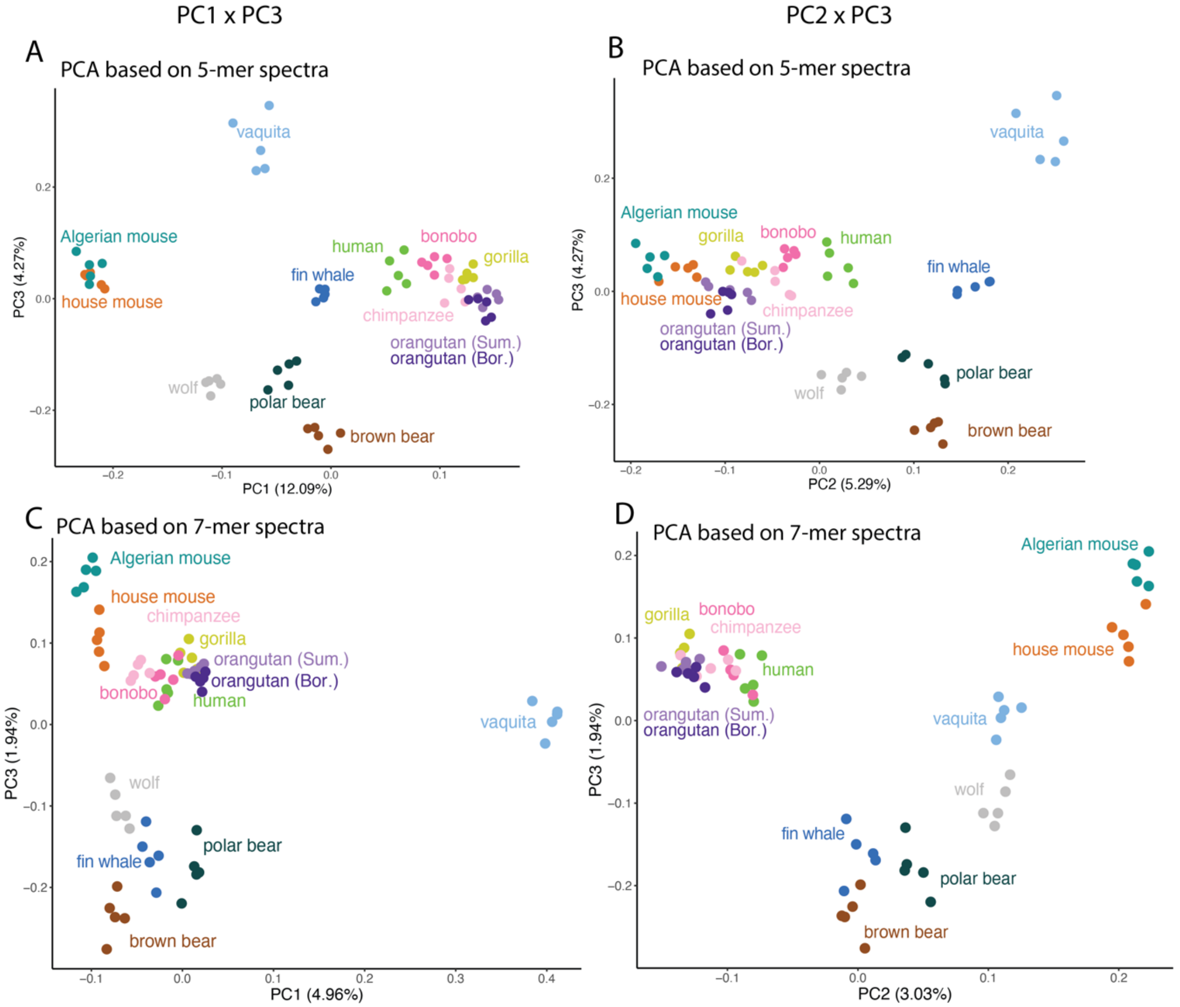

**Figure S17. Phylogenetic signal plots with labels.** These plots are identical to the distance plots shown in main text **Figure 4A** but with every point labeled with the specific species comparisons for reference.

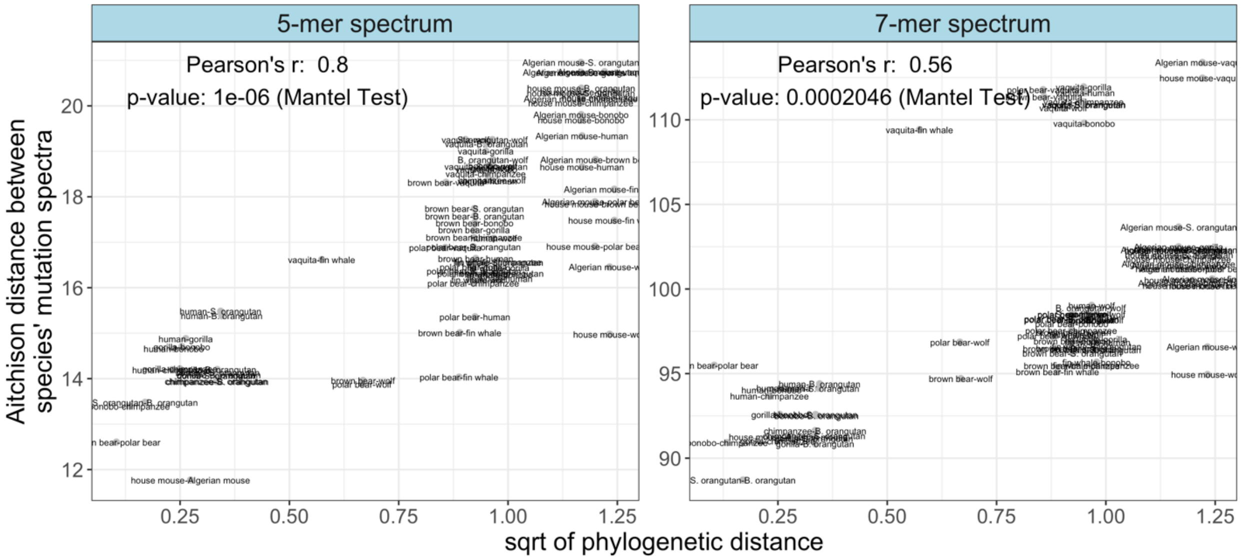


**Figure S18. Phylogenetic signal analyses based on an ultrametric timetree are consistent with distances based on the genetic alignment *RAxML* tree.** Distance plots with phylogenetic distance based on shared branch lengths from the ultrametric time tree (**Figure S1**). Results are qualitatively similar to results based on the tree in which branch lengths represent expected substitutions per site (**Figure 4A**). *p*-values based on the Mantel test with 9,999,999 permutations.

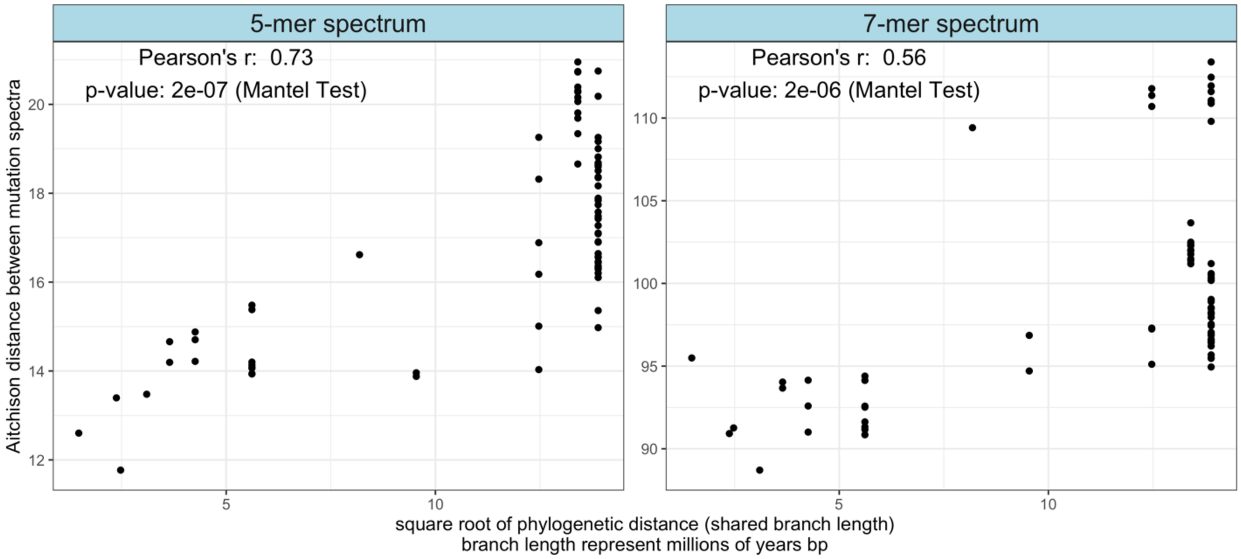

**Figure S19. Biased gene conversion does not drive phylogenetic signal. Distances based on the 5 and 7-mer spectra stratified by biased gene conversion mutation categories.** 5-mer and 7-mer spectrum distances were calculated based on *k*-mers that were separated into GC-biased gene conversion (BGC) categories based on their mutating central basepair: BGC-conserved mutations (BGC_conserved), consisting of A>T and C>G mutations which are not affected by biased gene conversion; strong-to-weak mutations (BGC_SW), consisting of C>A and C>T mutations which are disfavored by BGC; and weak-to-strong (BGC_WS) mutations, consisting of A>C and A>G mutations which are favored by BGC. Importantly, the correlation between 5- and 7-mer spectrum distances and phylogenetic distance are still significant in the BGC_conserved category, indicating that BGC is not causing the phylogenetic signal we observe. *p*-values based on the Mantel test with 9,999,999 permutations.

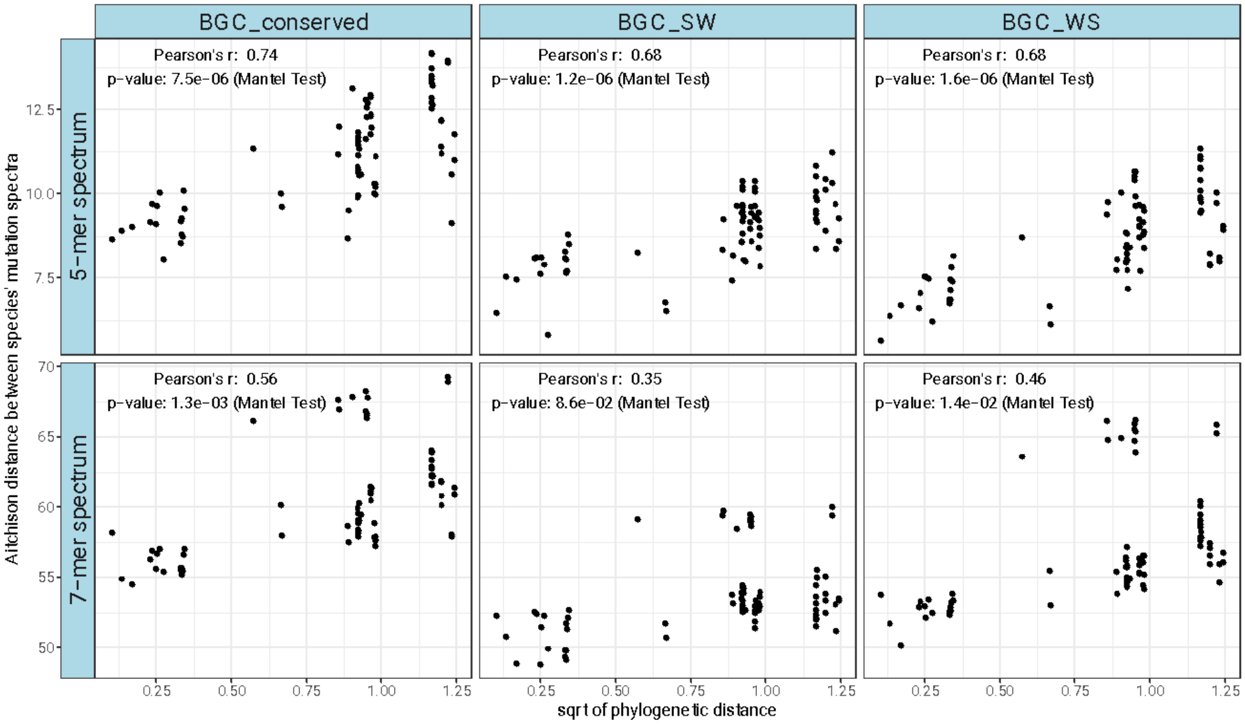

**Figure S20. Cosine distance is correlated with phylogenetic distance.** Plots showing the correlation between cosine dissimilarity (1-cosine similarity) and the square root of phylogenetic distance. *p*-values from the Mantel test with 9,999,999 permutations.

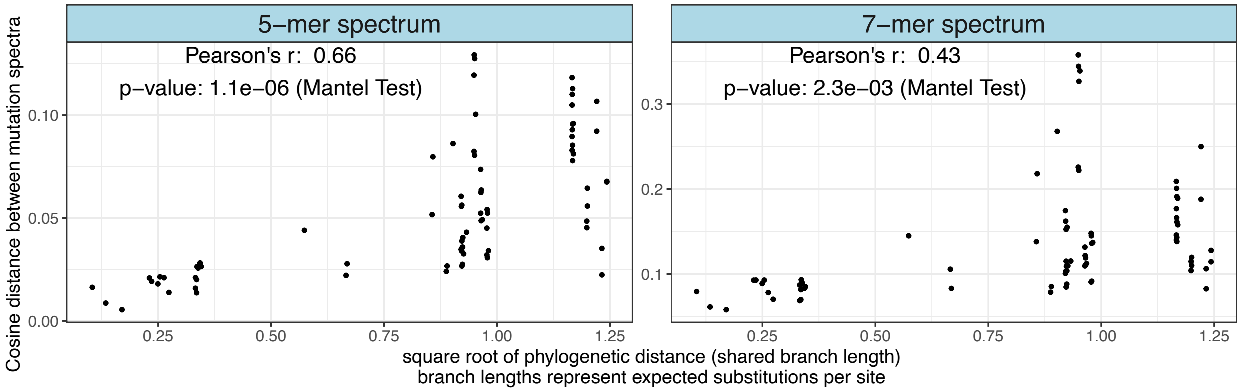


**Figure S21. Phylogenetic signal results are robust to mispolarization.** Repeating the analyses with a 'folded' mutation spectrum in which reverse mutation types are collapsed into the same category to determine whether the signal could be driven by mispolarization of ancestral alleles. For example, TA**<u>C</u>**AG>TA**<u>A</u>**AG and TA**<u>A</u>**AG> TA**<u>C</u>**AG are grouped inton the same 5-mer mutation category. Despite the reduction in power caused by analyzing fewer overall mutation types, our findings of significant phylogenetic signal persist, indicating that they are robust to mispolarization of ancestral mutation types. *p*-values based on the Mantel test with 9,999,999 permutations.

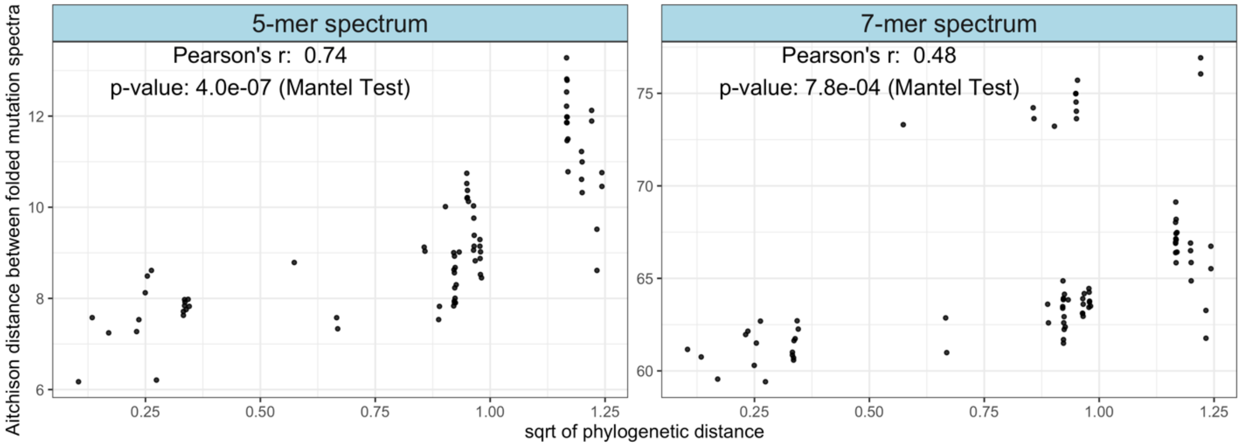

**Figure S22. Comparing the *k*-mer mutation spectrum to a mutation spectrum that has been permuted within (*k*-2)-mer mutation categories.** To determine whether the phylogenetic signal of 5-mers is entirely driven by underlying phylogenetic signal of 3-mers, we generated 5,000 control spectra for each species which randomized 5-mer mutation counts based on genomic target size within a central 3-mer to generate a pseudo-5mer spectrum that does not contain any phylogenetic signal beyond what is contained in the 3-mer spectrum. To demonstrate this principle, in the left panel we show the results of the Mantel test carried out on a single randomized replicate (red) compared to the empirical spectrum (black). In the right panel, we show a demonstration of a single randomized 7-mer replicate, for which we generated randomized mutation counts of 7-mers within central 5-mers to generate a 7-mer spectrum that does not contain any phylogenetic signal beyond the 5-mer spectrum. The full distributions of all 5,000 randomized datasets are shown in main text **Figure 4**. *p*-values calculated using the Mantel Test with 9,999,999 permutations.

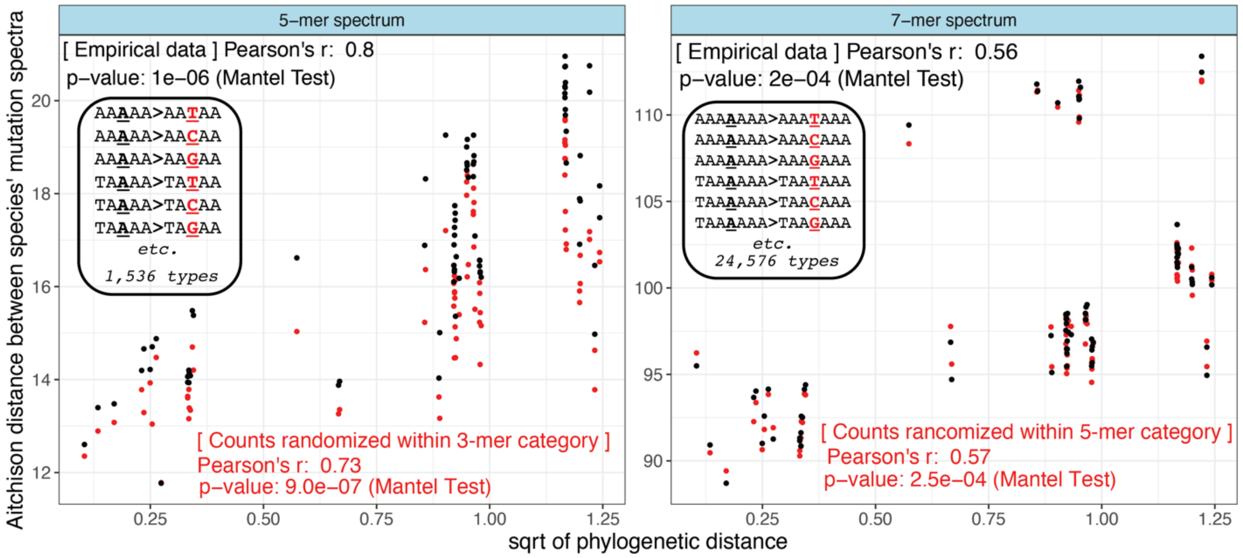

**Figure S23. Phylogenetic signal results are consistent when excluding low diversity species.** The low-diversity vaquita, polar bear and Gulf of California fin whale population result in needing to downsample all other species to ~130,000 SNPs. When mutation types are numerous this results in sparsely distributed data at the 5- and 7-mer spectrum level. In order to downsample in a less extreme manner, we excluded vaquita and polar bear, and switched to using the Eastern North Pacific fin whale population (ENP), which has higher diversity. This allowed us to downsample to ~890,000 SNPs instead of 130k, providing greater resolution on rarer mutation types, though also a reduction in power due to fewer species being included in the analysis. **A)** As in **Figure 4**, we compare the correlation between mutation spectrum distance and phylogenetic distance between the empirical dataset to a permuted dataset in which mutation counts are randomized across 5-mers within a central 3-mer, or across 7-mers within a central 5-mer. *p*-values based on Mantel test with 9,999,999 permutations. **B)** Distributions of *r* values from randomized control datasets, in which 5-mers have been randomized within their central 3-mer category (left panel), or 7-mers have been randomized within their central 5-mer category. Red lines denote empirical r values**.** **C)** Correlations between potential confounders and mutation spectrum distance. *p*-values based on the Mantel test with 99,999 permutations. **D)** Correlations between possible confounders and mutation spectrum distance, when the spectrum has been stratified by central basepair. *p*-values calculated using the Mantel test with 99,999 permutations. Note that fewer permutations can be used for the Mantel test in panels (C) and (D), since no test hits the minimum *p*-value for 99,999 permutations (1e-5)).

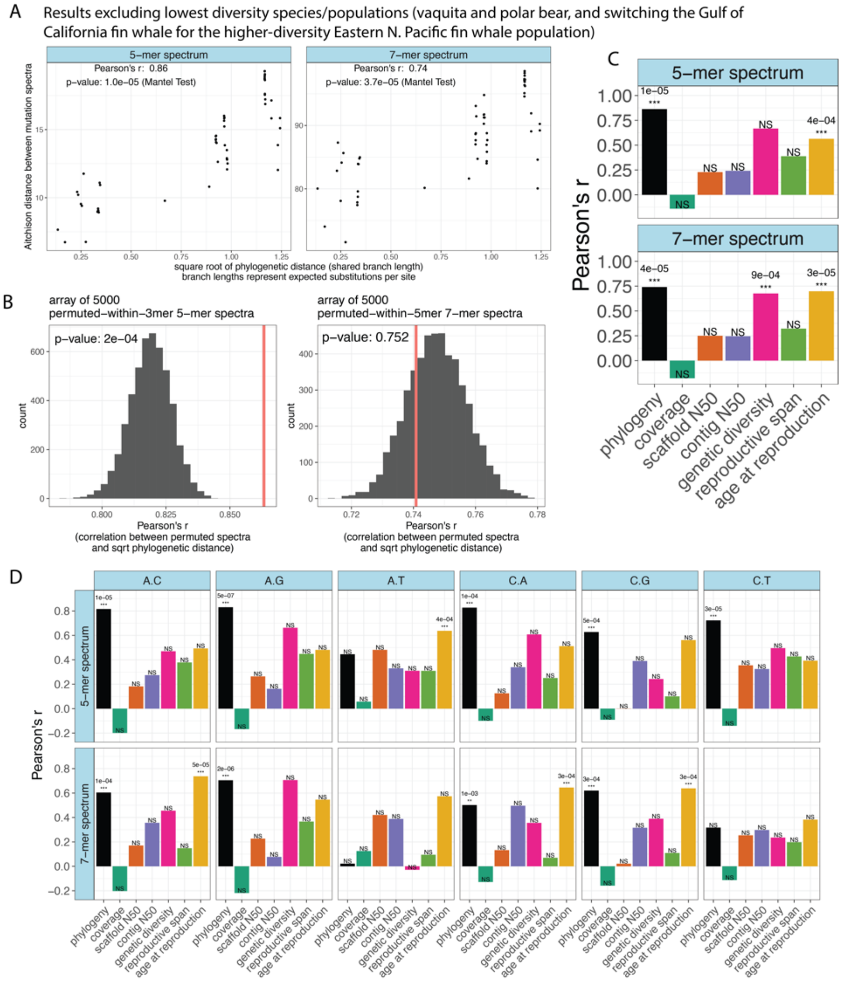

**Figure S24. Correlation of technical and biological confounders with the 5-mer and 7-mer mutation spectra.** Values of Pearson's *r* for the correlation in the differences between pairs of species' values for technical and biological variables and their 5-mer and 7-mer mutation spectrum distance, calculated using the Mantel test with 99,999 permutations. The black "phylogeny" columns represent the empirical phylogenetic signal *r*-values. "NS" (non-significant) denotes *p*-values that fell above the Bonferroni-corrected threshold $\alpha = 0.05/7$ confounders = 0.007. Results using a phylogenetically-aware Mantel test in **Figure S26**.

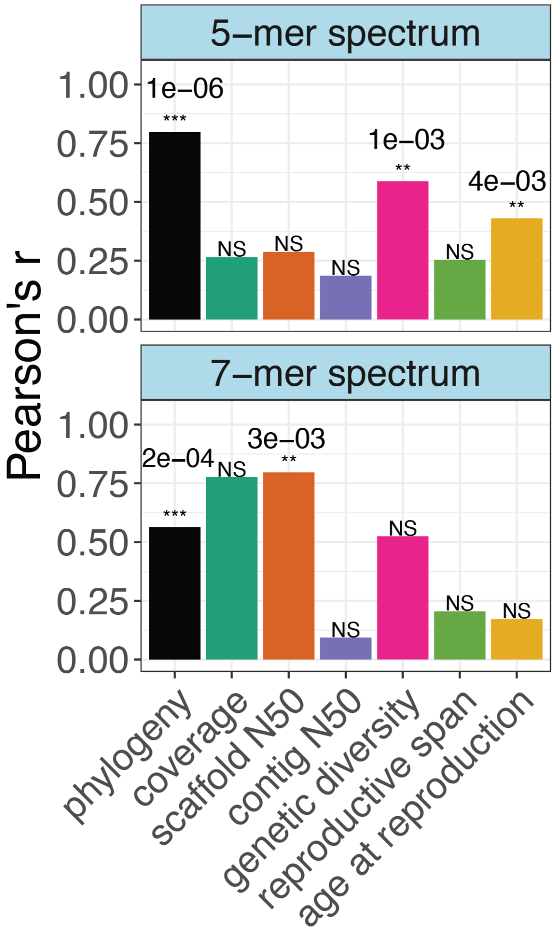

**Figure S25: 5-mer and 7-mer spectra faceted by central basepair still show phylogenetic signal and possibly more influence of technical confounders. A)** 5-mer and 7-mer mutation spectrum distances plotted against phylogenetic distance after stratifying by central 1-mer mutation type (e.g. in the "A>C" 5-mer panel, distances are calculated based on the 4,096 A>C 5-mers only). *p*-values based on Mantel test with 9,999,999 permutations. Mutation types without significant phylogenetic signal are grayed out per a Bonferroni-corrected significance threshold of 0.05/6=0.008. **B)** The significance of different variables to the 5-mer and 7-mer spectrum when they are stratified by central 1-mer type (as in (A)). The black "phylogeny" columns represent the r-values in (A). *p*-values calculated using the Mantel test with 99,999 permutations. *p*-values that fall above a significance threshold of $\alpha$ = 0.05/(7 variables * 6 mutation types) = 0.001) are noted as "NS" (non-significant). *p*-values that are < 0.001 are denoted "***" and the *p* -value is written above the corresponding column. Results using a phylogenetically-aware Mantel test in **Figure S26**.

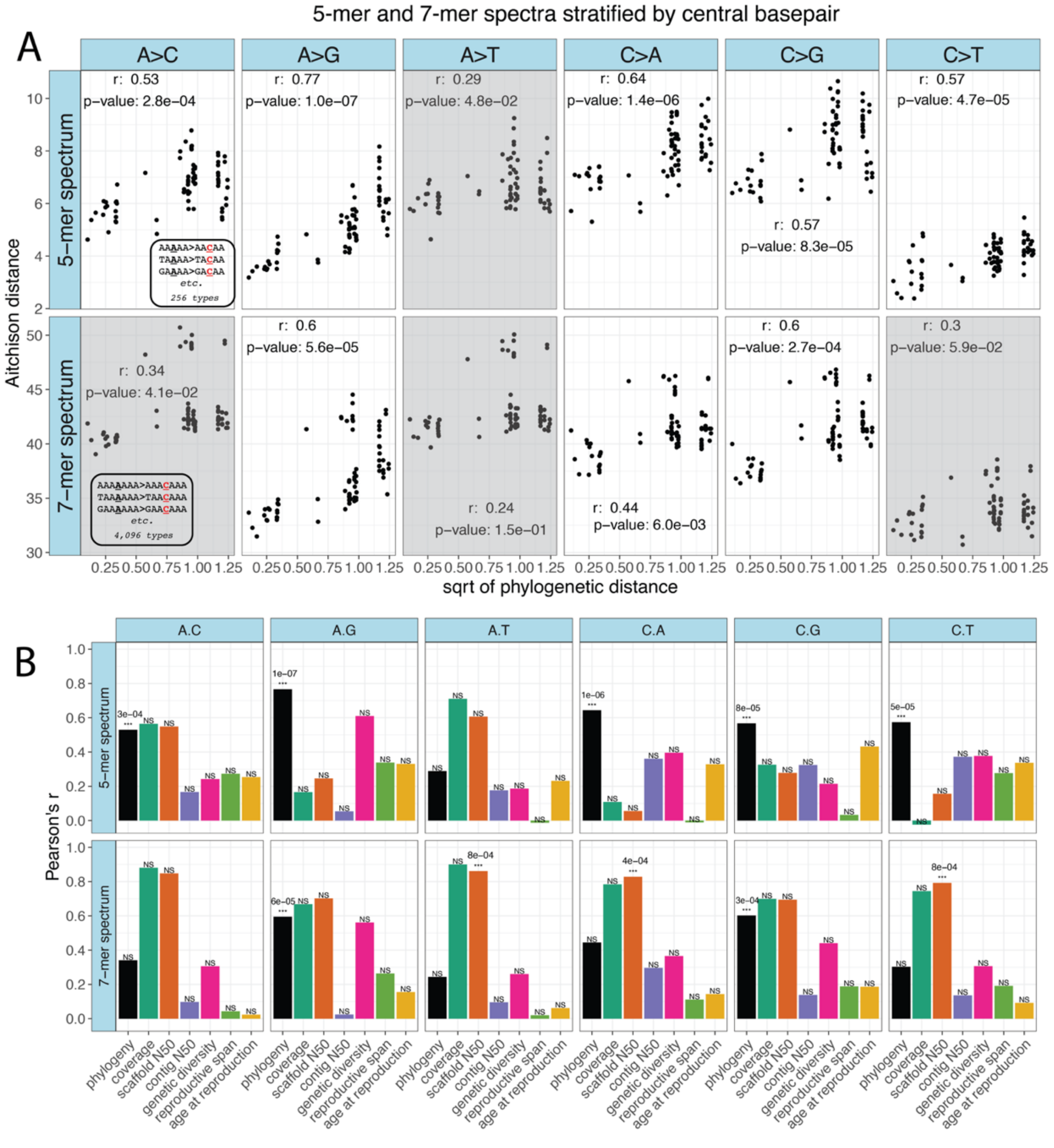

**Figure S26. Confounder correlation results are qualitatively similar when using a phylogenetically-aware Mantel test.** A version of the Mantel test called the phylogenetic permutation (PP) Mantel test ("phyloMantel") can be used to test for significant correlations between variables which may share a common phylogenetic signal that could cause falsely significant correlations if not corrected for. The test permutes species that are closely related in the phylogenetic tree with higher probability to generate a null set of permutations that incorporate any shared phylogenetic signal. The results are largely consistent across confounders between the uncorrected Mantel test and phyloMantel, though as seen in for the 1-mer and 3-mer spectra, genetic diversity becomes *more* significantly correlated with both spectra when using phyloMantel. *p*-values based on the Mantel test with 99,999 permutations (note that fewer permutations can be used for the Mantel test here, since no test is hitting the minimum *p*-value for 99,999 permutations (1e-5)).

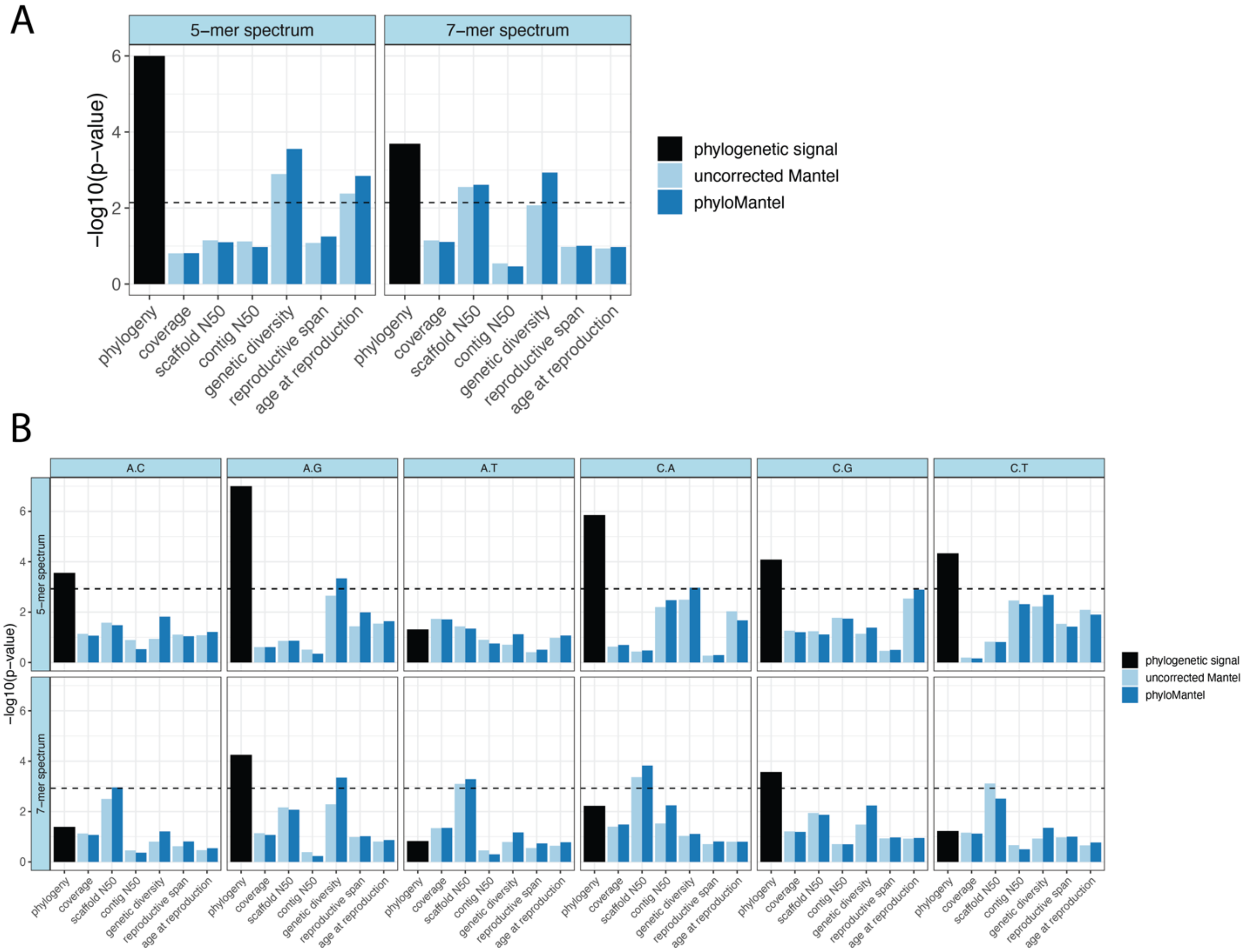

**Figure S27. 3-mer spectrum enrichment plots for all species and populations in the dataset.** As in main text **Figure 5**, but for 3-mer mutation types across all species in the dataset. The horizontal black dashed line represents the Bonferroni-corrected statistical significance threshold, and the red vertical dashed line is the species-specific mutability of CpG>TpG dimers relative to the background C>T rate. *k*-mers are colored by central mutation type.

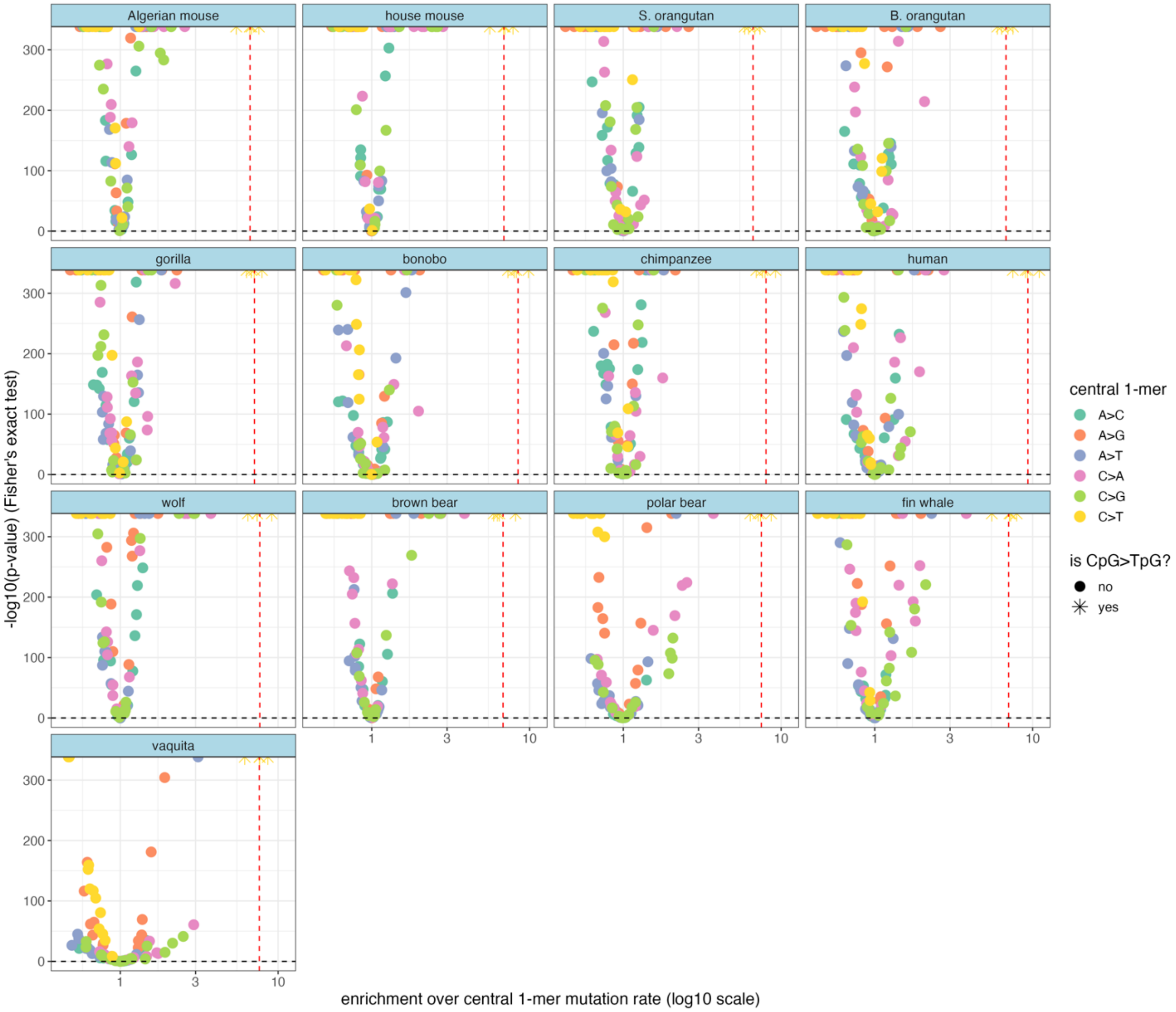

**Figure S28. 5-mer spectrum enrichment plots for all species and populations in the dataset.** As in **Figure 5**, the enrichment of 5-mers above their central 1-mer rate, here showing all species/populations in the full dataset. The 5 non-CpG>TpG 5-mers that have a higher level of enrichment than CpG>TpG dimers (red dashed line) are labelled. **Table S3** has a list of 5-mer mutation types that are enriched beyond species-specific CpG>TpG level.

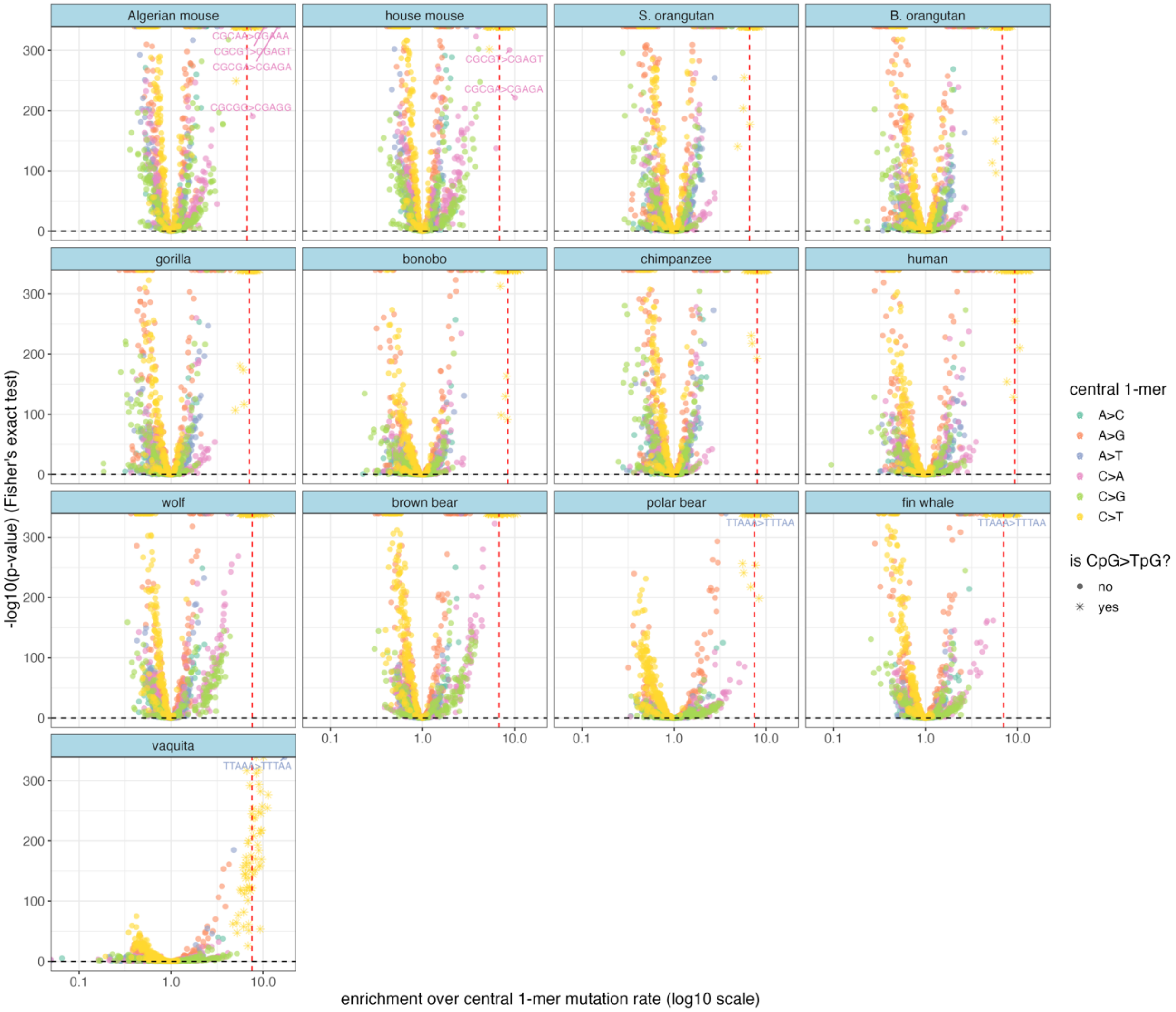

**Figure S29. 7-mer spectrum enrichment plots for all species and populations in the dataset.** As in **Figure 5**, the enrichment of 7-mers above their central 1-mer rate, here showing all species/populations in the full dataset. A small subset of the ~100 non-CpG>TpG 7-mers that have a higher level of enrichment than CpG>TpG dimers (red dashed line) are labeled. **Table S4** has a list of the significantly enriched 7-mer mutation types that are enriched beyond species-specific CpG>TpG level.

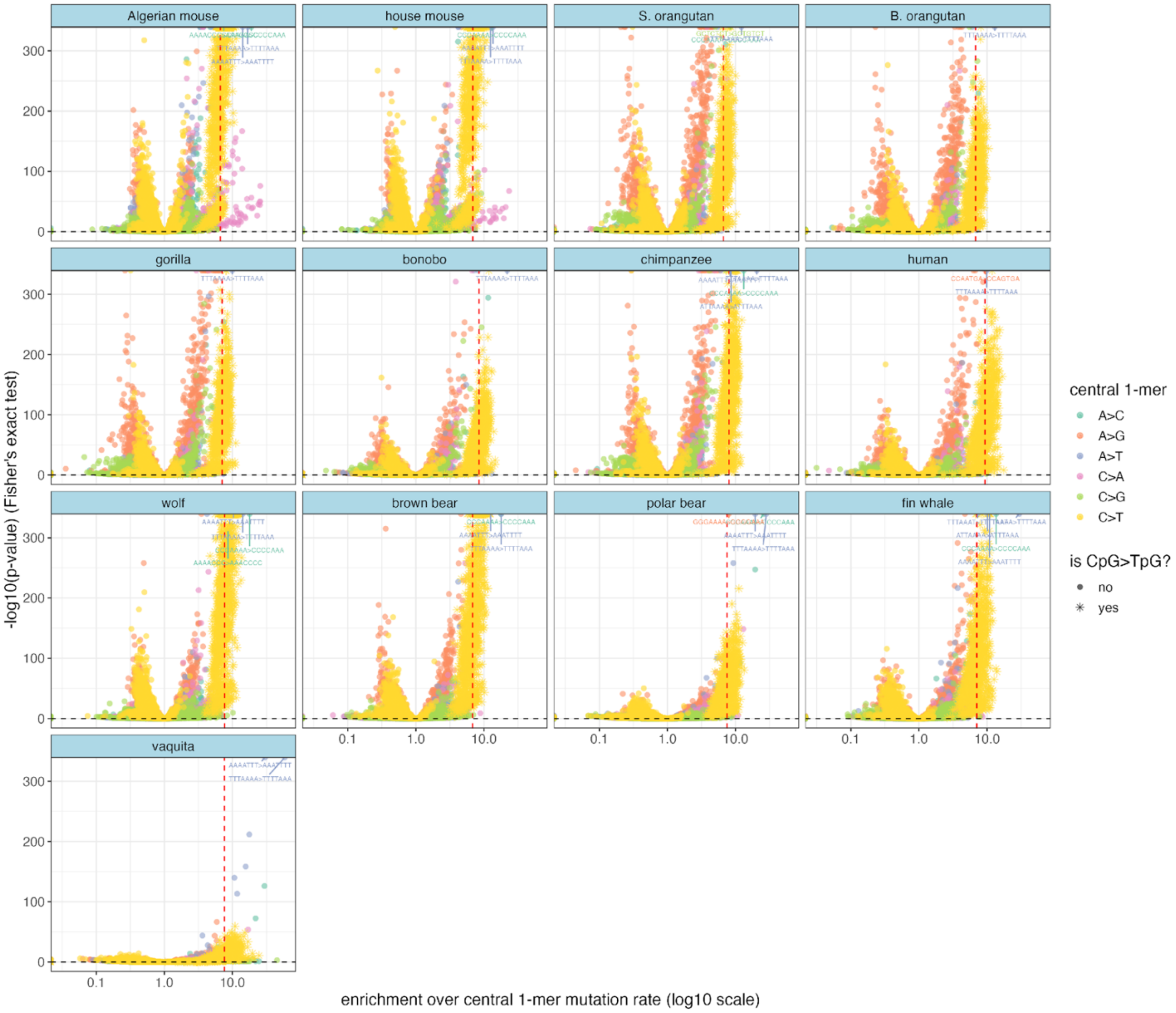

**Figure S30. SBS Signatures and novel 3-mer signature.** COSMIC single base substitution (SBS) cancer signatures: SBS1, associated with CpG>TpG mutations due to cytosine deamination, and SBS5, a signature of unknown etiology that is found in both germline and somatic mutation datasets and appears to accumulate in a clock-like manner in human somatic tissue. A third novel signature was extracted using *sigfit* from our species' empirical 3-mer spectra, while simultaneously fitting SBS1 and SBS5. Gray error bars in the novel signature represent the 95% credible interval.

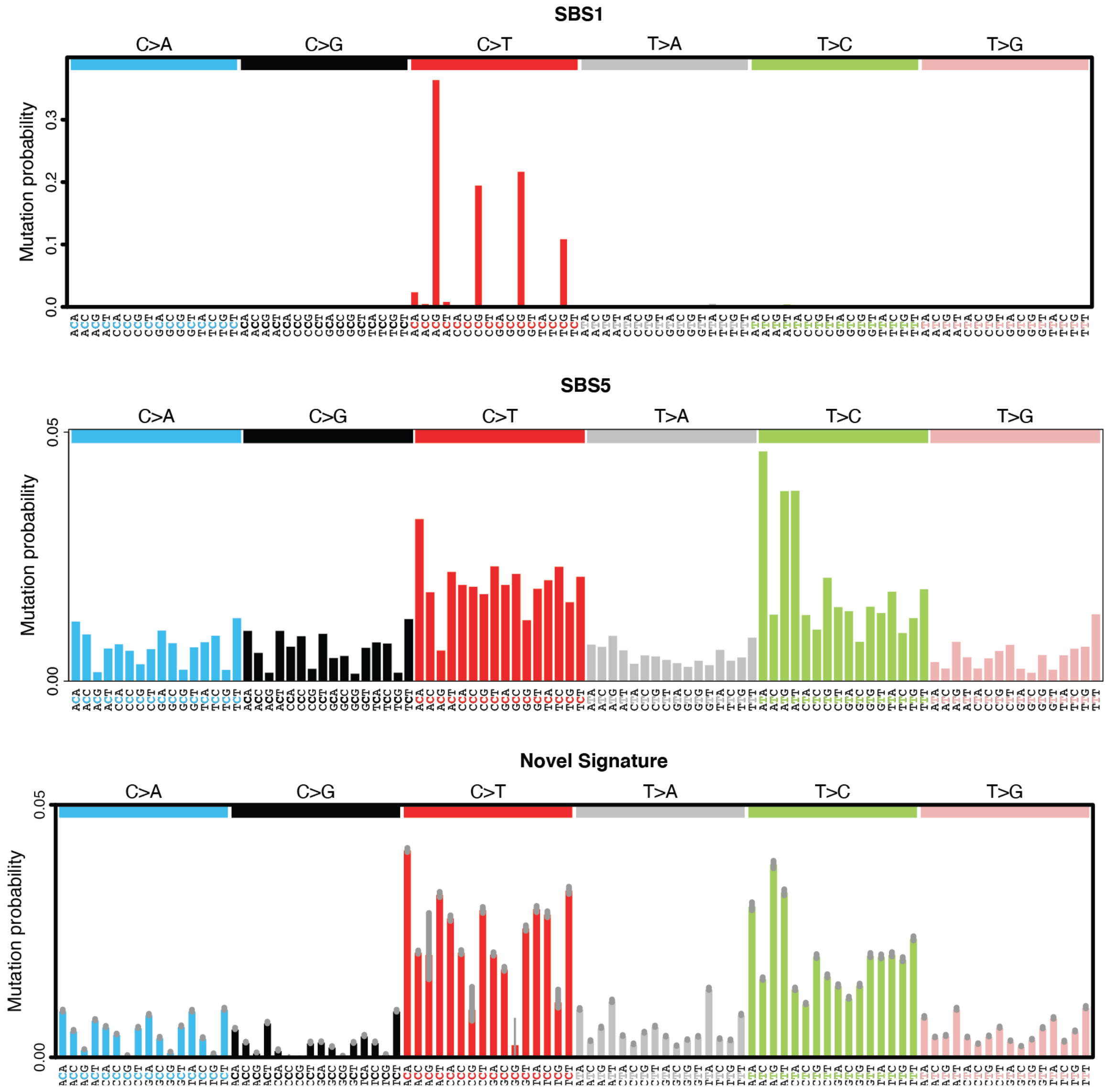

**Figure S31.** ***a priori*** **reproductive aging signatures and one novel signature extracted from the data (1-mer-CpG spectrum).** The "aging signatures" were calculated based on Poisson regressions carried out in Jónsson et al. (2017) based on Icelandic human family sequencing data. The maternal and paternal age signatures represent the signature of mutations associated with advancing maternal and paternal age at the time of conception. The "young parent" signature represents a signature associated with mutations accumulated prior to puberty. The novel signature was extracted from the empirical 1-mer-CpG (excluding CpG>TpG mutations) spectra of our species, while simultaneously fitting the three aging signatures.

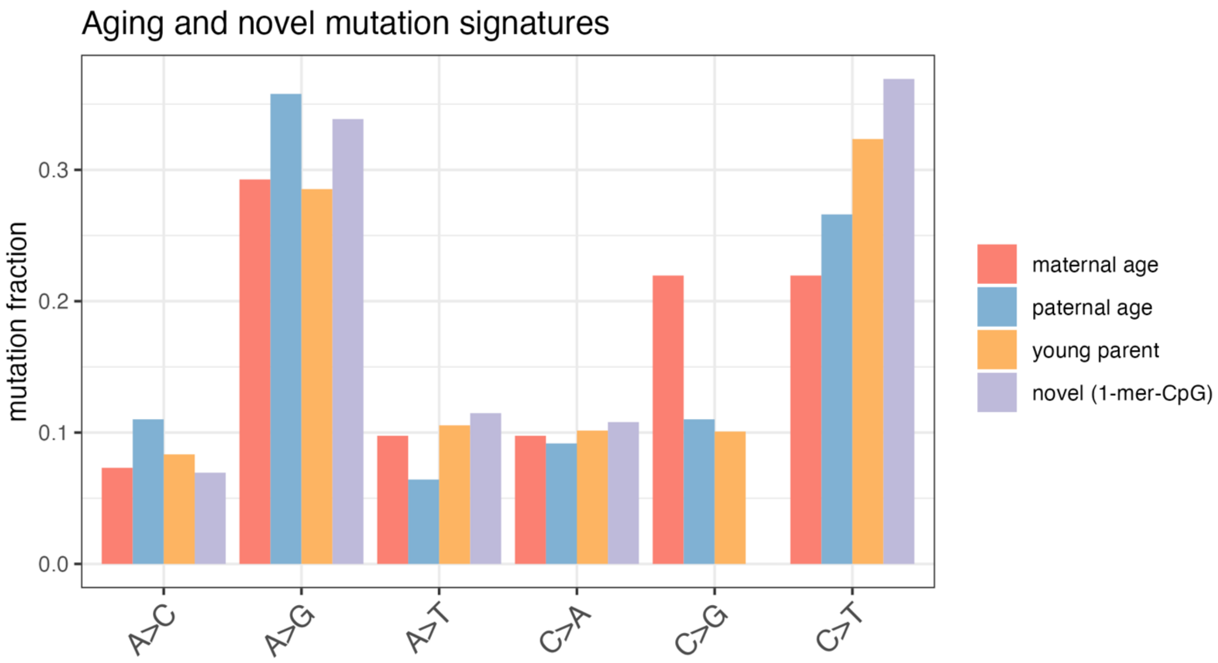

**Figure 32. Mouse-wolf similarities persist across independent datasets.** The Aitchison distance between grey wolf and mouse species' 1-mer and 3-mer spectra was lower than predicted given their phylogenetic distance. We set out to replicate this in independent datasets to assess whether underlying batch effects might be driving this pattern. We calculated spectrum distances between de novo mouse mutations from Lindsay et al. (2019) and an independent wolf dataset from Mooney et al. (2023) (labeled as wolf (UCLA)) to compare to our initial wolf dataset from the Broad Institute (labeled as wolf (Broad)). We found that the de novo mouse spectrum was closest to the mouse polymorphism spectra (which is reassuring), and the next-closest distance was to the two wolf datasets and the vaquita. We found that both wolf datasets had the closest distance to the vaquita, then to mouse species. Note that all the mouse DNM comparisons are elevated relative to their polymorphism-based counterparts, likely due to increased noise in the de novo spectrum and/or systematic differences between DNMS and polymorphisms, but the relative *ranking* of species ordered by increasing distance from the mouse DNMs or the mouse polymorphism datasets are highly similar. The "Lindsay mouse DNMs" are elevated for both mice and wolves above other more distantly related species for this reason as well.

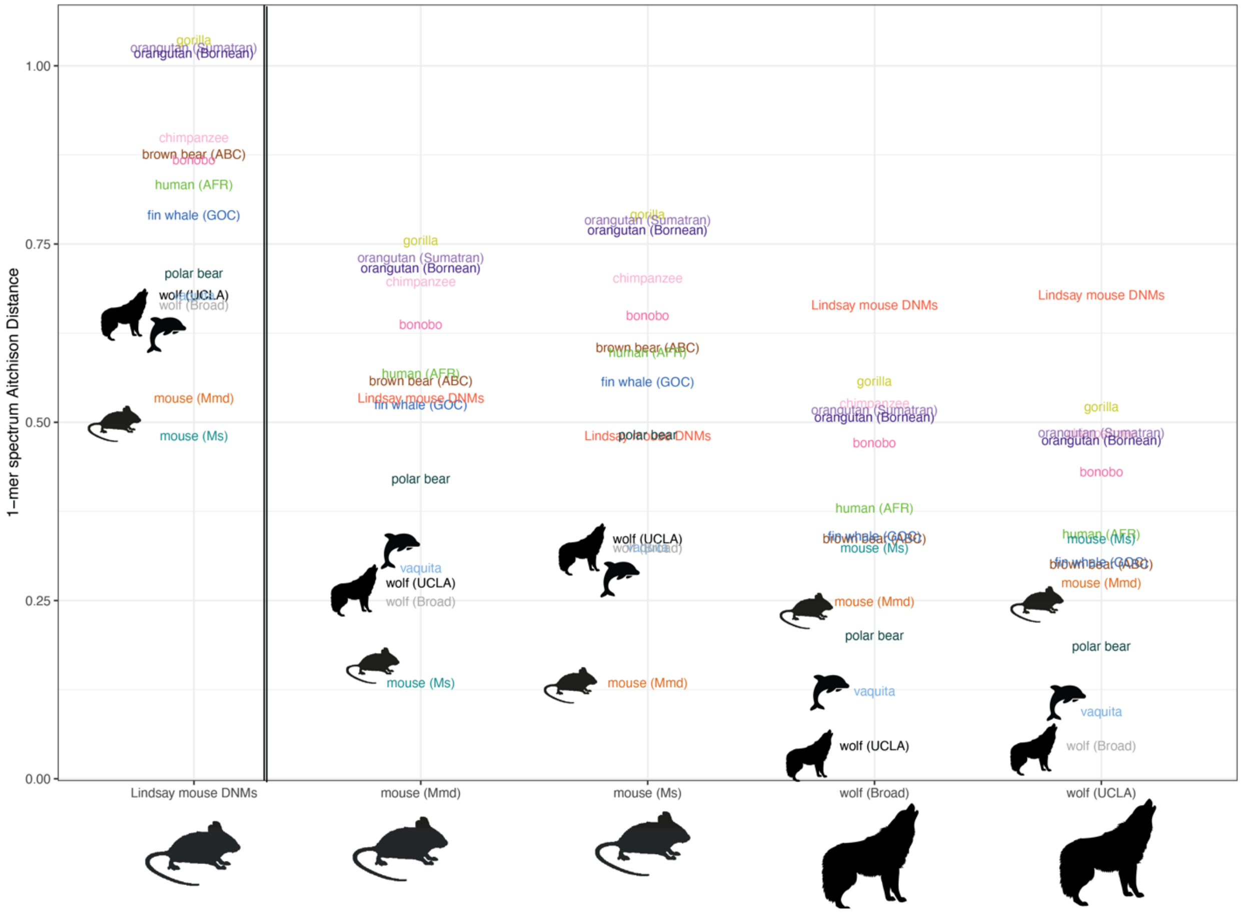

**Figure S33. Aitchison transformations allow for more informative spectrum comparisons. A)** Demonstration of differences in contribution of the different mutation types to the distance between two species (here human and gorilla), depending on which distance metric is used. In Aitchison distance, when each mutation type's rate has been centered log-ratio transformed, larger proportional differences in mutation abundance contribute more to the distance, whereas Euclidean distance is almost entirely dominated by high mutation rate NpCpG>NpTpG 3-mers. This is less of an issue if mutation fractions rather than rates are used, as CpGs make up a smaller fraction of the overall set of mutations.
**B)** Another commonly used metric in cancer biology is cosine similarity, which here is shown to be dominated by the most abundant 3-mers in the dataset.

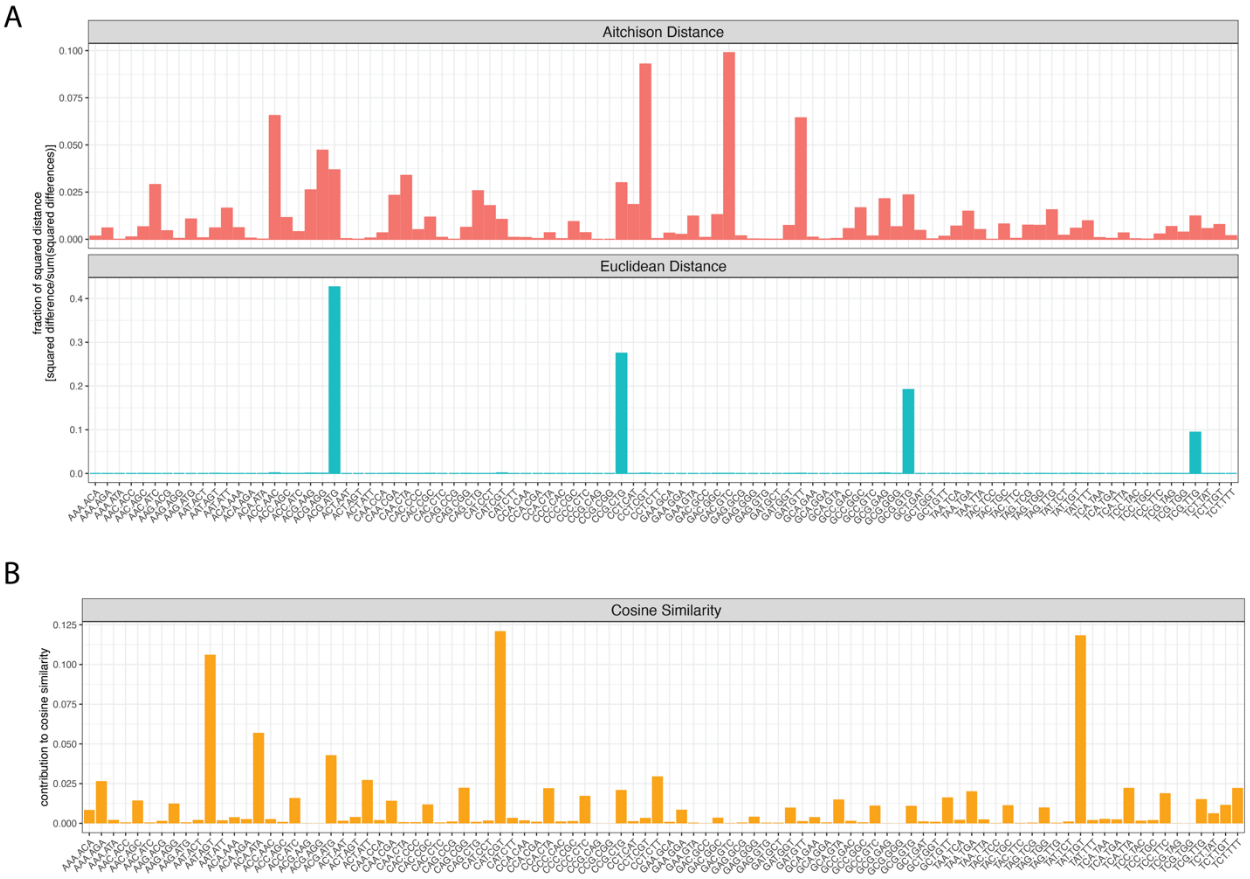

**Figure S34. After rescaling to the same genomic target content and carrying out the CLR transform, different ways of scaling count data yield the same results.** Once species' spectra have been scaled to have the same genomic content, and the centered log-ratio (CLR) transform has been performed, then mutation counts, mutation proportions (counts divided by sum of counts per species), or mutation 'rates' (counts divided by genomic target sizes) all yield the same results for PCA **(A)** or the distance between spectra **(B)**.

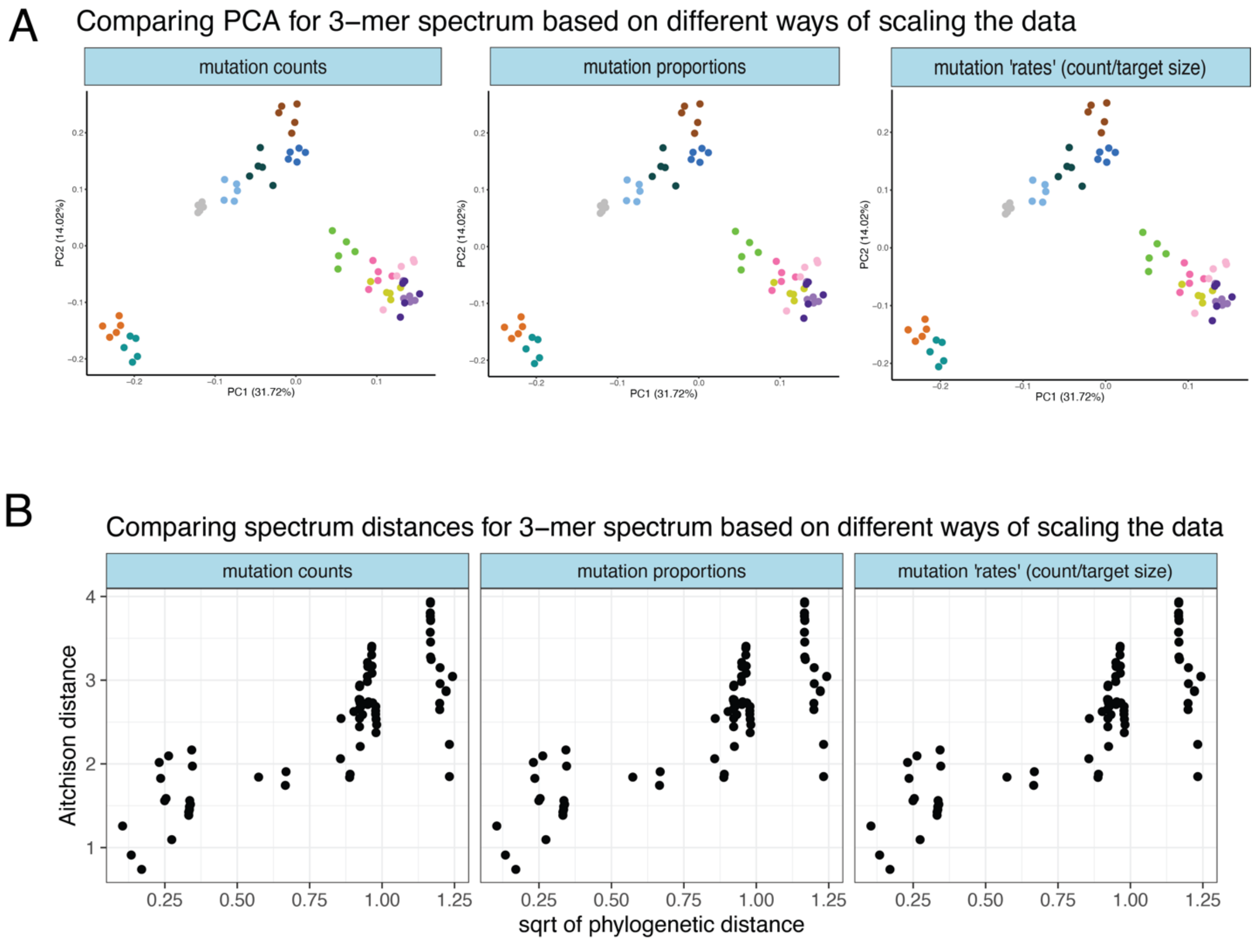
